# Multi-tissue metabolic GWAS and drought-responsive multi-omics reveal the genetic basis of the quinoa metabolome

**DOI:** 10.64898/2026.02.25.706111

**Authors:** Julia von Steimker, Elodie L. Rey, Clara Stanschewski, Regina Wendenburg, Annabella Klemmer, Markéta Macho, Venkatesh Thirumlaikumar, Noha O. Saber, Aleksandra Skirycz, Alisdair R. Fernie, Mark A. Tester, Saleh Alseekh

## Abstract

Quinoa (*Chenopodium quinoa*) is a nutrient-rich pseudocereal with diverse specialized metabolites, yet the genetic basis of this metabolic diversity is poorly understood. Here we integrate whole-genome sequencing and multi-tissue metabolic profiling of 603 quinoa accessions. We detected 4,688 metabolic features and identified over 1,000 metabolites in seeds, leaves, and roots. Using multi-tissue genome-wide association, we mapped the genetic architecture of quinoa metabolome by identifying 584 quantitative trait loci (QTL) and prioritized 219 candidate genes across 58 major QTL governing saponin, betalain, and flavonoid biosynthesis. Moreover, we constructed a drought-responsive multi-omics regulatory network and uncovered additional key genes involved in quinoa stress signalling and metabolic pathways. Finally, we cloned and functional validated the roles of CYP76AD1 in betalamate accumulation, UGT91C1 in flavonoid glycosylation, and CYP72A154 and soyasapogenol B glucuronide galactosyltransferase (SGT) in saponin biosynthesis. This multi-omic framework provides a high-resolution map of the quinoa metabolome and a foundation for breeding nutrient-rich and stress-resilient quinoa cultivars.

## Introduction

Quinoa (*Chenopodium quinoa* Willd.) is an allotetraploid crop (2n = 4x = 36) that originated in the Andean region of South America approximately 7,000 years ago (Jarvis et al., 2017; Risi and Galwey, 1989). It is emerging as a highly nutritious food, with seeds rich in gluten free protein and essential amino acids (Galwey *et al*., 1989; Nowak *et al*., 2016). Quinoa is adaptable to a wide range of agricultural systems due to its remarkable tolerance to multiple abiotic stresses, including salinity (Hariadi *et al*., 2011; Iqbal *et al*., 2020), drought (Fuentes and Bhargava, 2011), frost (Jacobsen *et al*., 2005; Jacobsen *et al*., 2007), and high ultraviolet radiation (González *et al*., 2009), rendering it a promising crop for ensuring food security under climate change.

Recent advances in quinoa research have led to the development of extensive genetic and genomic resources aimed at improving agronomic performance and stress resilience (Bodrug-Schepers *et al*., 2021; Rey *et al*., 2023; Yasui *et al*., 2016; Zou *et al*., 2017; Jarvis *et al*., 2017). These efforts have enabled the application of genome-wide association studies (GWAS) to uncover the genetic architecture of traits such as seed weight, mildew resistance (Patiranage *et al*., 2022), salinity tolerance (Zou *et al*., 2017), flowering time (Patiranage *et al*., 2022), and seed colour (Sandell *et al*., 2024).

The plant metabolome, characterized by vast chemical diversity, plays critical roles in development, environmental adaptation, and defense responses (Oksman-Caldentey and Saito, 2005; Pichersky and Lewinsohn, 2011; Raza, 2022). Many plant-derived metabolites, such as vitamins, flavonoids, phytosterols, phenolics, and carotenoids, also contribute significantly to human health as nutritional and medicinal compounds (Fraga *et al*., 2019; McChesney *et al*., 2007; Sreenivasulu *et al*., 2023). Over the past decades, integrative omics approaches – including metabolomics, genomics, and transcriptomics – have proven effective in elucidating gene function and metabolic pathways (Alseekh, Kostova, *et al*., 2021; Shen *et al*., 2023). Exploring metabolite diversity and its underlying genetic variation offers valuable insights for crop improvement, germplasm conservation, and the understanding of domestication (Alseekh, Scossa, *et al*., 2021; Fernie *et al*., 2021; Sreenivasulu *et al*., 2023).

GWAS and metabolome quantitative trait locus (mQTL) mapping have been successfully employed in numerous crops to reveal the genetic basis of metabolic variation (Fang *et al*., 2019) and to identify loci associated with quality traits and environmental responses (Tieman *et al*., 2017). As a result, biosynthetic and regulatory genes for key metabolite classes – such as flavonoids in rice (Xia *et al*., 2023; Tiozon *et al*., 2023) and maize (Zhou *et al*., 2019), steroidal glycoalkaloids in tomato (Zhao *et al*., 2023), and saponins (Fiallos-Jurado *et al*., 2016; Lim *et al*., 2020; Jarvis *et al*., 2017) – have been identified and functionally characterized.

Despite recent progress in quinoa genomics and phenotypic characterization, the genetic basis of metabolic diversity in quinoa remains largely unexplored. Notably, quinoa metabolites such as saponins, betalains, and flavonoids contribute to both seed quality – impacting flavor, nutrition, and processing – and plant stress tolerance (Vega-Gálvez *et al*., 2010; Zhou *et al*., 2021; Davies *et al*., 2018; Szakiel *et al*., 2011). For example, betalains, exclusive to the Caryophyllales order, are potent antioxidants that influence seed coloration and protect against drought, UV, and salinity (Timoneda *et al*., 2019). Their biosynthesis originates from L-tyrosine through pathways involving cytochrome P450-mediated reactions, yielding betalamic acid and *cyclo*-DOPA or through dopamine by decarboxylation, generating a highly structurally diverse compound class by spontaneous conjugation and condensation reactions (Timoneda *et al*., 2019; Nakatsuka *et al*., 2013).

Saponins are abundant in quinoa seed coats, contributing to bitterness and requiring removal prior to consumption (Melini and Melini, 2021; Otterbach *et al*., 2021). Nonetheless, they play protective roles against pathogens and pests (Kuljanabhagavad *et al*., 2008) and some have recognized health benefits (Yu *et al*., 2022). Their biosynthesis involves squalene epoxidation, followed by complex cyclization, oxidation, and glycosylation steps, which generate a diverse spectrum of saponins through these secondary modifications (Phillips *et al*., 2006; Thimmappa *et al*., 2014; Sawai and Saito, 2011). Saponin content is also thought to differentiate bitter and sweet quinoa varieties and is largely regulated by transcription factors such as TSARL1 and TSARL2 (Fiallos-Jurado *et al*., 2016; Lim *et al*., 2020; Jarvis *et al*., 2017), although recent studies suggest flavonoids and polyphenols as main bitterness driver (Song *et al*., 2024). Similarly, flavonoids are a major class of metabolites with diverse physiological functions in plants and implications for human health (Wang *et al*., 2019; Zhuang *et al*., 2023).

To date, most metabolomic studies in quinoa have examined a limited number of accessions, and no large-scale metabolic GWAS (mGWAS) has been conducted. Understanding the genetic basis of quinoa’s metabolome is therefore critical for leveraging natural variation in key functional metabolites. In this study, we integrated whole-genome sequencing with untargeted LC-MS-based metabolic profiling across three tissues – seeds, leaves, and roots – of 603 quinoa accessions. Using mGWAS, we dissected the genetic architecture of metabolite accumulation, with an emphasis on saponins, flavonoids, and betalains in a tissue-specific manner. Furthermore, through a multi-omics approach incorporating proteomics and metabolomics under drought stress, we identified key molecular components involved in stress adaptation. Our findings provide a foundational resource for dissecting metabolic regulation in quinoa and offer promising targets for breeding nutritionally superior and stress-resilient cultivars.

## Results

### The genomic diversity of the quinoa diversity panel

Previously we analysed a quinoa core collection of 310 accessions, reporting 2.9 million polymorphic single-nucleotide variants (Patiranage *et al*., 2022). Here, we made use of and characterized a quinoa diversity panel comprising 603 *C. quinoa* accessions originating primarily from Argentina, Bolivia, Chile, Peru, and the United States, spanning elevations from 12 to 4,899 meters above sea level (**Fig. 1a**). Whole-genome sequencing yielded 13.7 million SNPs, of which 1.45 million high-confidence variants were retained after stringent quality control (60,417 - 104,938 SNPs per chromosome; transition/transversion ratio = 1.53; **Fig. S1a**).

**Figure 1.**
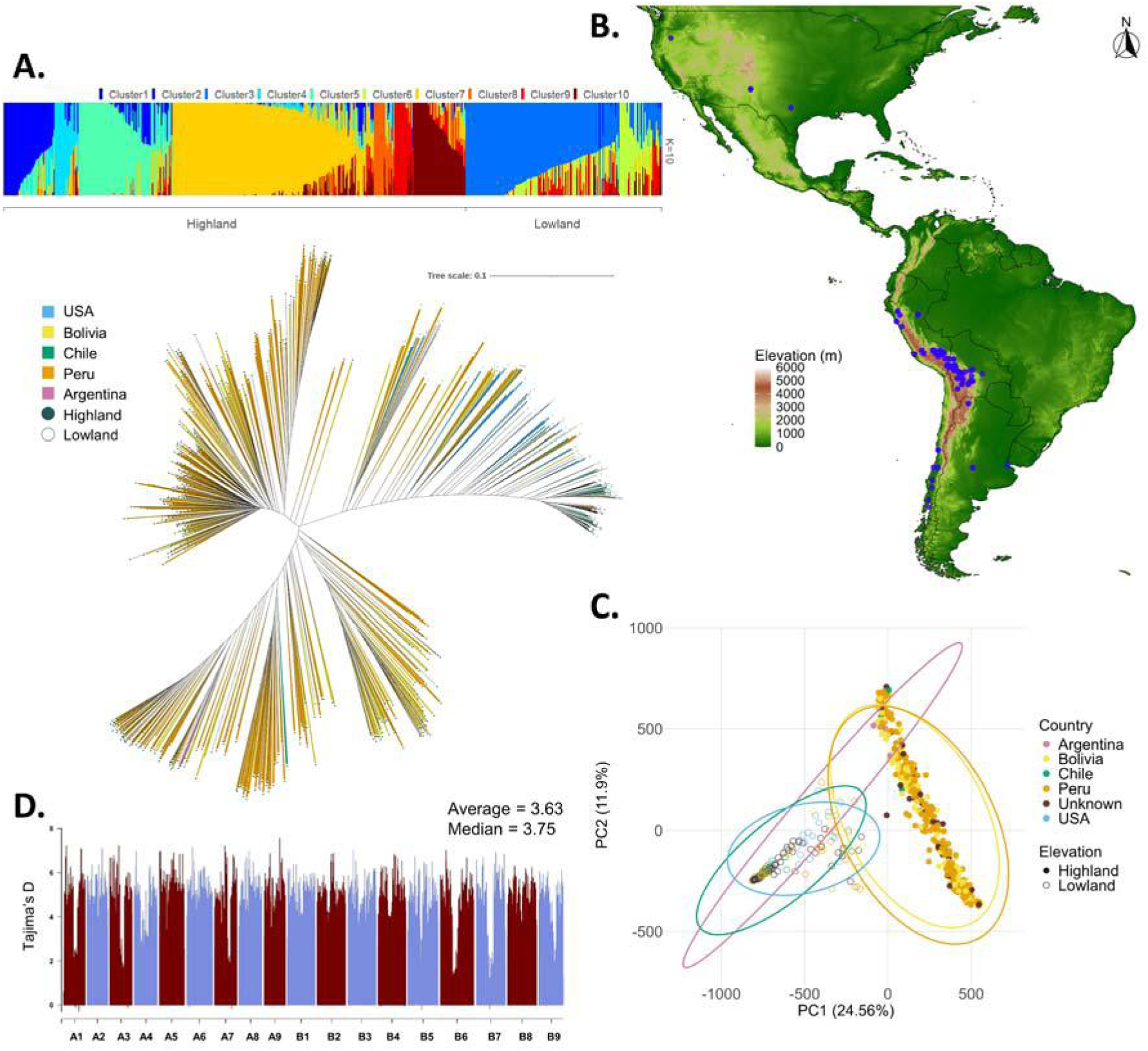
Genomic diversity of 603 C. quinoa accessions. (**A**) Neighbor-joining tree of 603 accessions with inferred population structure (K = 10, corresponding to the model complexity that maximized marginal likelihood), grouped by geographic origin. (**B**) Geographic distribution of quinoa accessions across the Americas, with collection sites (blue dots) shown in relation to altitude. (**C**) Principal component analysis (PCA) of 603 accessions, color-coded by geographic origin and shaped by elevation, illustrating genetic stratification largely along an altitudinal gradient. (**D**) Genome-wide Tajima’s D values calculated in 100 kb non-overlapping bins, showing predominantly positive values indicative of high genetic diversity and balancing selection within the population.

Population structure analysis resolved ten ancestral components, yet a single-component model captured most of the variance, indicating that diversity within the panel is shaped by fine-scale, rather than deeply divergent population differentiation. Geographic assignments revealed that clusters 2, 4, 5, and 7 – 10 were enriched for highland accessions, whereas clusters 3 and 6 largely lowland (**Fig. 1a**). Geographic assignments revealed that clusters 4, 5, 7, and 10 were enriched for Bolivian and Peruvian accessions, whereas cluster 6 was largely Chilean (**Fig. S1b**). Cluster 5 was predominantly associated with bitter quinoa accessions, while cluster 7 was enriched for sweet types. By contrast, saponin content did not show a clear correspondence with population structure.

Genomic principal component analysis (PCA) explained 24.6% of the total genetic variance, primarily reflecting geographic origin: Chilean and U.S. accessions formed a distinct cluster from Bolivian and Peruvian accessions (Patiranage *et al*., 2022), a pattern closely correlated with the elevation of collection sites (**Fig. 1b,c**). This suggests altitude as a major driver of population differentiation. Within-group variance accounted for 11.9% of the total genetic variation, a pattern that was recapitulated using a representative subset of accessions for later GWAS analysis (**Fig. S1c**). Genome-wide diversity was further supported by a strongly positive Tajima’s D (mean = 3.63, median = 3.75; **Fig. 1d; Table S1**), indicating an excess of intermediate-frequency alleles consistent with balancing selection or population admixture. Data completeness was high, with 532 accessions retaining > 90% of SNP calls (**Fig. S1d**), and inbreeding coefficients exceeded 0.72 in 464 accessions (**Fig. S1d; Table S2**), confirming extensive homozygosity and validating the suitability of this panel for high-resolution genome-wide association studies.

### Multi-tissue metabolic diversity in the quinoa GWAS panel

We profiled specialized metabolites and lipids in seeds of 588 *C. quinoa* accessions, and specialized metabolites in roots and leaves of a subset of 166 accessions using liquid chromatography–mass spectrometry (LC–MS). In total, we detected 4,688 polar secondary metabolites and 4,949 apolar lipid features in seeds, 1,203 metabolic features in roots, and 608 in leaves (**Fig. 2a**). PCA revealed clear tissue-specific metabolic variation, with seeds explaining 13.1% and 6.9% of variance along PC1 and PC2, respectively, leaves 18.4% and 10.6%, and roots 17.1% and 10.8%. Importantly, secondary metabolite variation in seeds mirrored genomic structure, with highland accessions separating from lowland accessions (**Fig. 1c, 2a**). This trend was observed in the polar but not the apolar seed fraction (**Fig. S2**).

**Figure 2.**
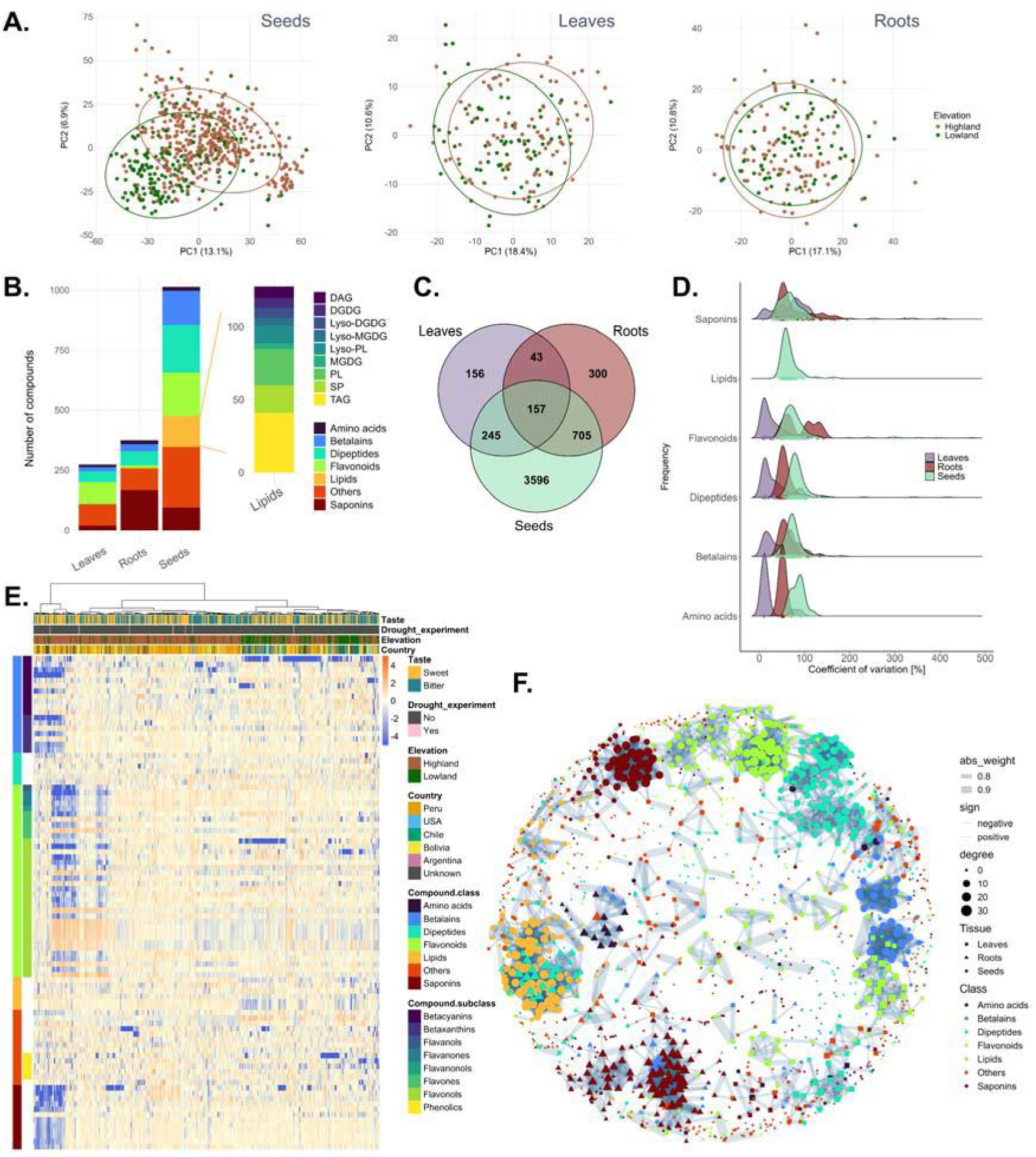
Metabolic diversity of the *C. quinoa* diversity panel across seeds, leaves, and roots. (**A**) Principal component analysis of 4,706 seed, 609 leaf, and 1,210 root metabolic features from 573, 166, and 167 accessions, respectively. Accessions are color- or shape-coded by geographic origin or elevation. (**B**) Distribution of compound classes per tissue (excluding unknowns): 275 in leaves, 375 in roots, and 1,014 in seeds, including 128 lipids. (**C**) Venn diagram of shared and tissue-specific metabolites across seeds, leaves, and roots. (**D**) Ridgeline plot of the coefficient of variation [%] for leaf, root, and seed secondary metabolites. (**E**) Heatmap of 96 annotated seed metabolites with significant genomic associations, including 18 betalains, 7 dipeptides, 38 flavonoids, 6 lipids, 12 saponins, and 15 other metabolites like sugars or organic acids. Accessions are annotated by taste type (sweet/bitter), origin, and elevation; those used in the drought experiment are highlighted in pink (Ames-13760, LM2, CHEN-199, D-12393, CHEN-384, D-12165, PI-665276). (**F**) Network analysis of annotated seed, leaf, and root metabolites and lipids based on Pearson correlation (|r| > 0.7, p < 0.05), visualized using the Kamada–Kawai layout.

We annotated 1,014 metabolites in seeds, 375 in roots, and 275 in leaves (**Fig. 2b; Tables S3, S4**). In seeds, the dominant classes included lipids (12.6%, mainly triacylglycerols), betalains (14.1%, primarily betacyanins), flavonoids (17.7%, mostly flavonols), and dipeptides (19.5%). Roots were enriched in saponins (44.2%), while leaves were dominated by flavonoids (32.7%, largely flavonols). Across tissues, 157 metabolites were shared, comprising diverse chemical classes such as dipeptides (15.9%), saponins (9.5%), betalains (8.9%), and flavonoids (5.7%) (**Fig. 2c, Table S5**). Pairwise comparisons revealed 43 metabolites shared between leaves and roots, 245 between leaves and seeds (notably flavonoids), and 705 between roots and seeds (enriched for saponins).

Variation within compound classes, quantified by the coefficient of variation (CV), highlighted amino acids, betalains, and dipeptides as highly variable (mean CV: 85.1%, 75.3%, and 86.2%, respectively). Flavonoids showed greatest variability in roots (90.9%) and seeds (85.9%), while saponins were highly variable in both roots (78.7%) and seeds (78.3%) (**Fig. 2d**). Heatmap analyses underscored extensive diversity across accessions, particularly in dipeptides, flavonoids, and saponins (**Fig. 2e, S3**). In seeds, a distinct subgroup of accessions diverged strongly from the main population, driving separation along PC1 (**Fig. 2a, 2e**).

Network analysis resolved 19 major metabolic clusters (≥10 compounds) using Louvain community detection (**Fig. 2f; Table S6**). These included three saponin clusters (two root-enriched, one seed-associated), four dipeptide clusters (two seed, one root, one leaf), three lipid clusters (seed), and four flavonoid clusters (two seed, two leaf). Additional clusters comprised betalains (two), amino acids (one), and mixed compound classes (two). For example, ferulate strongly correlated with paselate methyl ester (r = 0.78), while together with ferulate isorhamnetin-3,7-*O*-glucoside correlated with *p*-coumaroylmalate (r = 0.80, 0.71). The other mixed leaf cluster highlighted strong correlations among arginine, glutamyl-arginine, histidine, pyroglutamate, and proline-betaine (r ≥ 0.71).

We further identified a secondary metabolite (m/z 505.227, retention time 4.68 min) as a major determinant of sweet versus bitter taste, consistent with previously described accessions (Patiranage *et al*., 2022; Kollmar *et al*., 2025). Using this marker, we classified accessions according to seed taste and identified compounds correlated with sweetness and bitterness (**Fig. S4; Table S7**). In seeds, differential abundance was observed for six amino acids, 58 betalains, 74 dipeptides, 114 flavonoids, 47 saponins, 48 lipids, and 90 other metabolites including for instance sugars, organic acids and amino acid derivatives. In roots, 53 saponins, 13 other compounds, seven betalains, six dipeptides, and one flavonoid distinguished sweet and bitter accessions. In leaves, differences were detected for five dipeptides, 13 flavonoids, four saponins (including basellasaponin-B and soyasaponin-II), and 14 other metabolites. The most intriguing features were the higher abundance of phytolaccoside E in leaves (FC = 0.47) of bitter varieties and a higher abundance of quercetin-dihexoside (FC = 2) in seeds, three flavonoids (Quercetin-3-*O*-2-*O*-galloyl-arabinoside, kampferol-hexoside, luteolin), and soyasaponins-II in leaves (FC > 2), and seven saponins in roots (hederagenin-pen-3x hex, goyasaponin-III, licoricesaponin A3, cloversaponin-I, gomphrenin-I, isogomphrenin-I, licoricesaponin E2; FC > 2).

To investigate metabolite divergence between domesticated quinoa and its wild ancestor, we conducted untargeted metabolic profiling of *Chenopodium suecicum*. Among a total of 828 shared polar metabolic features, 208 compounds were structurally annotated (**Fig. S5**). Of these, 88 exhibited significant differences in abundance between *C. suecicum* and *C. quinoa* 600 panel. PCA positioned *C. suecicum* within the broader cluster of *C. quinoa* accessions, with variation along PC1 (16.1%) largely driven by saponin abundance. This pattern suggests that divergence in saponin content either originated from *C. pallidicaule* or arose following the hybridization event leading to modern quinoa. When comparing *C. suecicum* with a single quinoa accession (CHEN-543), however, PCA revealed a much clearer separation, explaining 89.6% of the variance (**Fig. S6**). Overall, 474 metabolites differed significantly in abundance between *C. suecicum* and *C. quinoa* (adjusted *p* < 0.05, |log₂FC| > 1), of which 119 could be assigned to metabolic classes including amino acids, betalains, flavonoids, dipeptides, and saponins.

Together, these results reveal extensive metabolic diversity across tissues in the quinoa diversity panel, highlight the tissue-specific clustering of major metabolite classes, and identify key compounds underlying the sweet–bitter taste distinction. Moreover, the comparative metabolic profiling of *C. suecicum* supports the hypothesis that metabolic divergence – particularly in saponin-related traits – either arose post-hybridization or stemmed from other wild progenitors, offering further insight into the evolutionary history of the specialized metabolite repertoire of quinoa.

### Genome-wide association study of the quinoa metabolome

We performed GWAS on 4,688 polar secondary metabolite features and 4,949 lipid-related apolar features in seeds, as well as 1,203 and 608 polar secondary metabolite features in roots and leaves, respectively. Six complementary models were applied (see materials and methods). Compounds were first pre-filtered using FaST-LMM with genotype binning and HE as the variance component, and only associations surpassing a Bonferroni threshold were carried forward. Subsequent analyses employed MLM, CMLM, MLMM, FarmCPU, and BLINK (**Fig. S7**). To ensure robustness, only SNPs identified by at least four methods were retained (**Table S8-S9**). This strategy yielded 615 high-confidence marker-trait associations (MTAs), encompassing 119 seed apolar metabolites (82 MTAs), 652 seed polar metabolites (365 MTAs), 209 root metabolites (137 MTAs), and 44 leaf metabolites (46 MTAs; **Table S9-11**). Of these, 295 MTAs corresponded to annotated compounds (26 leaf, 66 root, 205 seed MTAs; **Tables S9–S10**). In seeds, significant associations were observed for 22 betalains, 34 cholesterol derivatives, seven dipeptides, 39 flavonoids, 163 saponins, six lipids, and 46 additional metabolites. In roots, associations involved one amino acid, seven betalains, five dipeptides, three flavonoids, 53 saponins, and ten other metabolites. In leaves, we detected associations for one betalain, three dipeptides, nine flavonoids, three saponins, and eight other metabolites.

To identify candidate genes associated to metabolic traits, we integrated GWAS with comparative genomics (OrthoFinder, **Fig. S8**), published transcriptomic resources, and new transcriptomic and proteomic datasets generated from a drought experiment on seven genetically diverse accessions selected based on seed metabolic profiles (**Fig. 6**). Using weighted gene co-expression network analysis (WGCNA), we identified 46 distinct co-expression modules that organized into four major clusters. Together, these data provide a multi-omic framework that integrates metabolic GWAS with transcriptomic and proteomic variation, thereby strengthening the identification of candidate genes underlying metabolic diversity in quinoa.

### Genomic hotspots and candidate genes underlying metabolic diversity

To validate the robustness of our mapping, we recovered TSARL1 and TSARL2 as candidate genes in a major hotspot QTL on chromosome B5 (**Fig. 3a, S9a, S10b,c; Tables S8–S11**). This is consistent with previous reports identifying this locus as a key QTL for saponin abundance (Jarvis *et al*., 2017; Patiranage *et al*., 2022). Interestingly, adjacent to these genes, we detected six UDP-glycosyltransferases (UGTs; *CQ004358*, *CQ004359*, *CQ004360*, *CQ004361*, *CQ004362*, *CQ004365*), three of which were differentially expressed under drought conditions and belong to the black (*CQ004358*), tan (*CQ004359*) and blue module (*CQ004361*).

**Figure 3.**
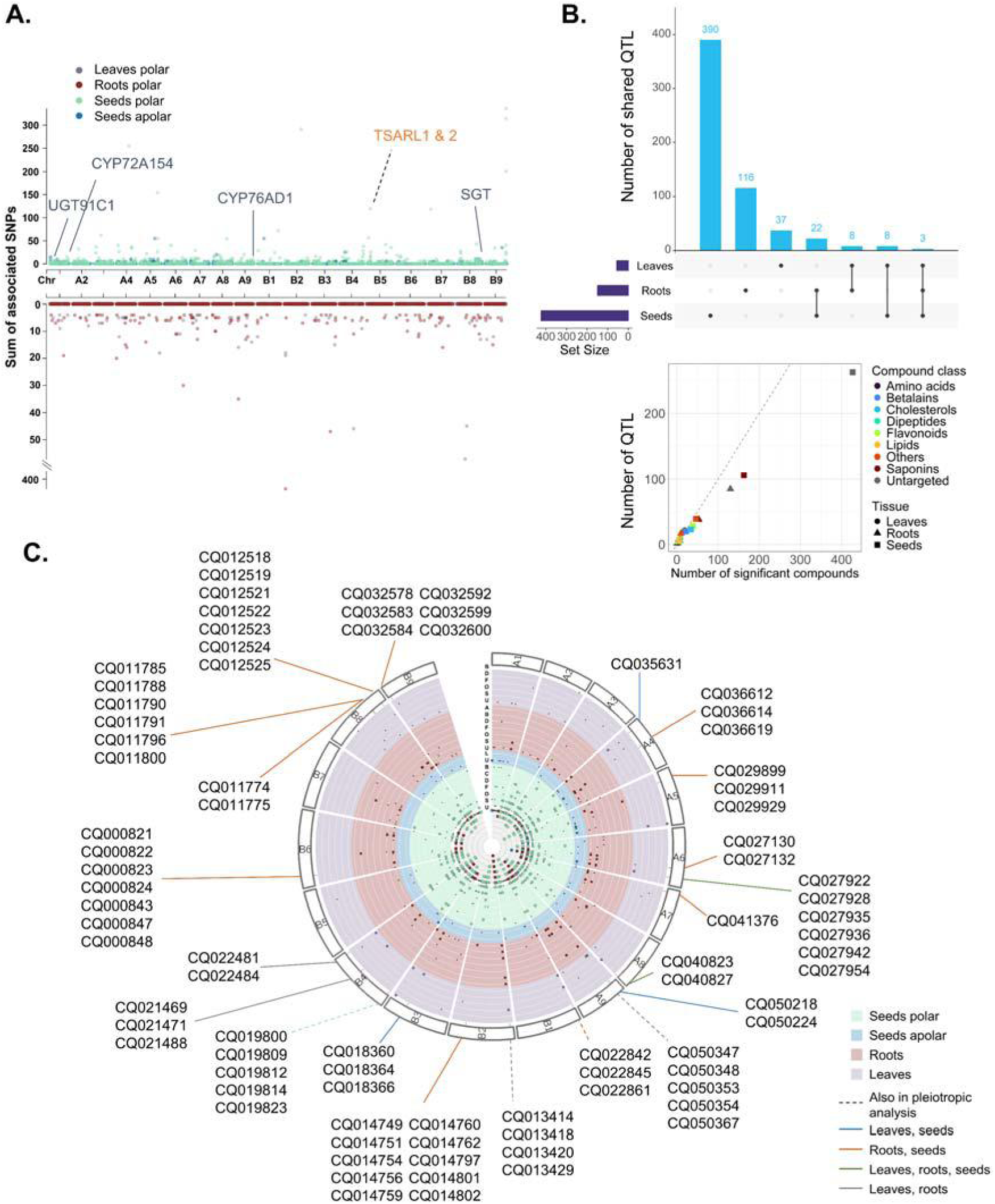
Multi-tissue QTLome of *C. quinoa* secondary metabolites. (**A**) Number of metabolic features with significant genomic associations across tissues: 652 seed secondary metabolites, 119 seed lipid features, 44 leaf secondary metabolites, and 209 root secondary metabolites, detected in 573, 166, and 167 quinoa accessions, respectively. In total, 365, 82, 46, and 137 marker–trait associations (MTAs) were identified for seed secondary metabolites, seed lipids, and leaf and root secondary metabolites, respectively. (**B**). Intersections of QTL in a 50 kb sliding window across tissues. As input the sum of the marker-trait associations across compound classes was used. In total 584 QTL were detected with 41 overlaps. Number of identified QTL in a 50 kb sliding window versus number of compounds with a GWAS association grouped by compound class and shaped by tissue. (**C**) Pleiotropic analysis in a 50 kb window size of seed polar and apolar metabolites, root metabolites, and leaf metabolites with their candidate genes. The outer ring represents the chromosomes; the inner rings show compound classes per tissue; and the dots indicate SNPs associated with genomic regions. The innermost circle summarizes the total number of associations per QTL. Compound class abbreviations: A = amino acids, B = betalains, D = dipeptides, F = flavonoids, L = lipids, O = other compounds, S = saponins, U = unknown.

In addition, GWAS for elevation revealed three QTL overlapping with the genomic PC1 reported by Patiranage et al. (2022), largely reflecting adaptation to altitude (**Fig. 9b**). Given the role of oxygen limitation at high elevations (Loreti and Perata, 2020; Giuntoli and Perata, 2018), we further investigated hypoxia-related pathways and identified 35 candidate genes involved in hormone signaling, development, and transcriptional regulation (**Table S12**).

In total, we identified 584 QTL associated with secondary metabolites across seed, root, and leaf tissues (**Fig. 3b**). Among these, 41 QTL were shared between tissues, while 390, 116, and 37 QTL were uniquely detected in seeds, roots, and leaves, respectively. Notably, for seed and root saponins as well as untargeted compounds, the number of detected QTL was lower than the number of significant metabolites, suggesting that multiple compounds co-localize to common genomic regions – indicative of regulatory QTL exerting pleiotropic control over entire metabolic pathways. Most QTL, SNPs, and significantly associated compounds were linked to saponins in roots and seeds, and to flavonoids in leaves (**Fig. S10a**). Across compound classes, we detected 493, 157, and 49 QTL in seeds, roots, and leaves, respectively, of which 148, 46, and 6 QTL were shared among multiple compound classes (**Fig. S11a**). The genomic distribution of QTL revealed major hotspots on chromosomes A1, A4, A5, and B8 for seed metabolites; A4, A6, A7, B3, B6, and B7 for root metabolites; and A1, A4, A9, B8, and B9 for leaf metabolites (**Fig. S11b**).

Genome-wide hotspot mapping, based on the density of significant associations and pleiotropic signals across tissues, revealed 58 novel QTL comprising 219 candidate genes (**Fig. 3c, S12; Table S12**). These included 26 UGTs, 20 transporters, 34 cytochrome P450 enzymes, 34 transcription factors (including nine basic helix–loop–helix [bHLH]), and 23 sugar-, methyl-, or acetyltransferases. Tissue-specific candidates comprised 10 candidate genes in leaves, 11 in roots, and 112 in seeds, while pleiotropic analyses identified 14 candidates shared between leaves and roots, 50 between roots and seeds, six between seeds and leaves, and eight common to all three tissues. Two of which, an arginase (*CQ047623*), and fructose-bisphosphate aldolase (*CQ007727*) were differentially regulated under drought belonging to the blue module of the drought experiment.

Examples of key loci include: (i) a leaf QTL on chromosome 5A at 61.7 Mb associated with ferulate and decarboxybetanin, harboring two CYPs (*CQ032038*, *CQ032039*), two bHLH transcription factors (*CQ032045*, *CQ032046*), and three amino acid transporters (*CQ032055*, *CQ032056*, *CQ032058*) (**Fig. S12a**); (ii) a multi-tissue QTL on chromosome A9 at 9.2 Mb containing three ABC transporters (*CQ050347*, *CQ050348* [grey module], *CQ050353*), a 4-hydroxy-3-methylbut-2-en-1-yl diphosphate synthase (*CQ050354,* [blue module]) involved in isoprenoid biosynthesis, and CYP81D1 (*CQ050367*), associated with betalains and saponins in leaves and roots (**Fig. 4, S12b**); and (iii) two major seed QTL: one on chromosome 4 at 43.2 Mb harboring seven CYPs, including 11-oxo-β-amyrin 30-oxidase (*CQ036986*), three CYP72A219 (*CQ036984*, *CQ036987*, *CQ036990*), and three CYP72A15 (*CQ036985*, *CQ036996*, *CQ037000*) (**Fig. S12c**); and another on chromosome B8 at 9.2 Mb comprising 12 CYPs (*CQ010896* - *CQ010921* [*CQ010911* pink module; *CQ010912* brown module; *CQ010921* turquoise module]) and two bHLH transcription factors (*CQ010905*, *CQ010906*), associated with sterol- and saponin-derived metabolites.

**Figure 4.**
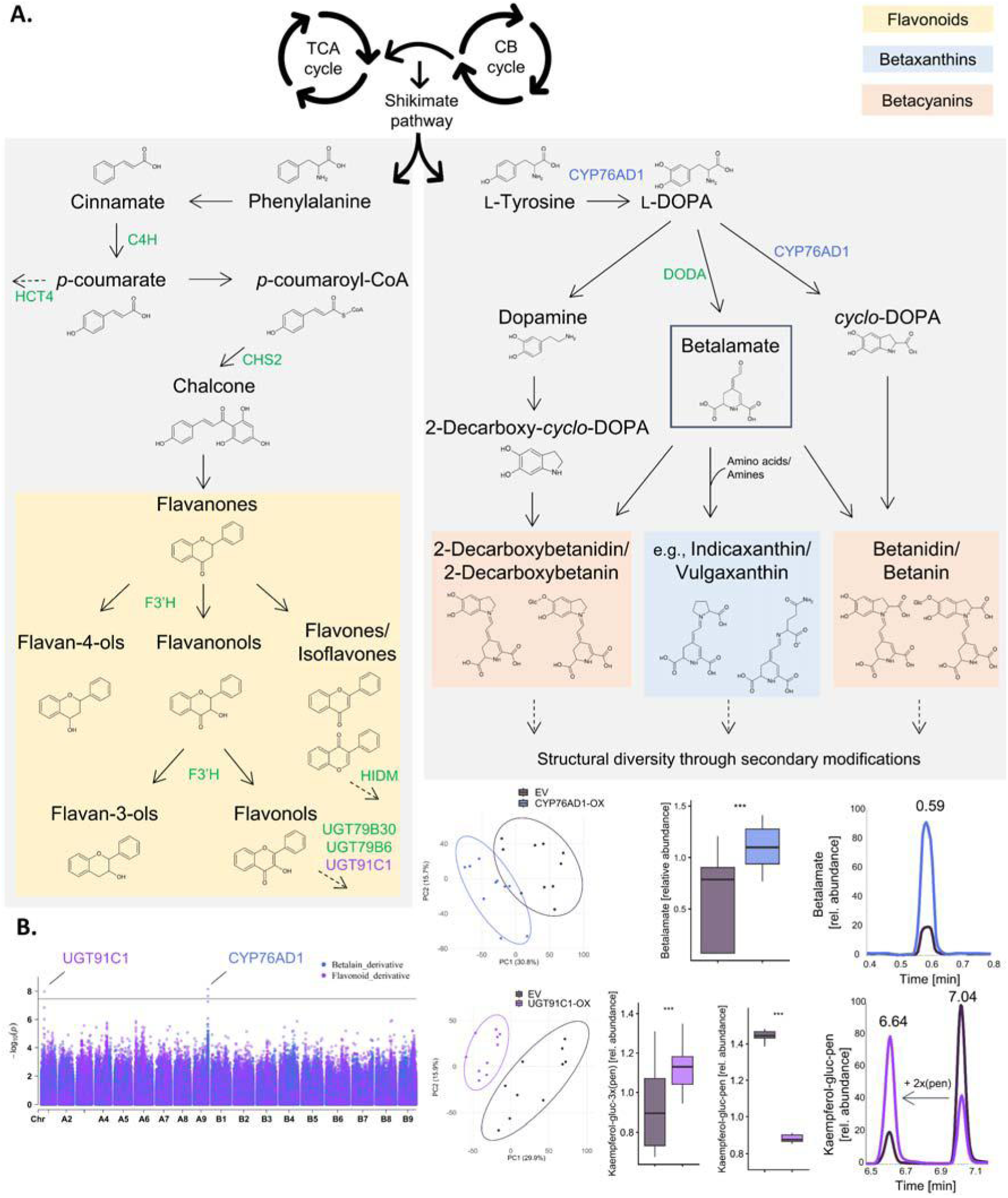
Refinement of the flavonoid and betalain pathways in *C. quinoa*. (**A**) The flavonoid pathway originates from phenylalanine and proceeds through phenylpropanoid intermediates such as cinnamate, *p*-coumarate, *p*-coumaroyl-CoA, and chalcone. From these precursors, diverse flavonoid structures are synthesized, including flavanones, flavan-4-ols, flavanonols, flavones and isoflavones, flavan-3-ols, and flavonols. The betalain pathway, in contrast, derives from L-tyrosine via L-DOPA, *cyclo*-DOPA, betalamate, and dopamine, leading to the formation of betacyanins (e.g., betanidin, betanin) and betaxanthins (e.g., indicaxanthin, vulgaxanthin). These core molecules serve as substrates for secondary modifications such as glycosylation and oxidation. Candidate genes are marked in green, validated genes are colored in purple and blue. (**B**) Overlay Manhattan plots for a representative betalain derivative and a flavonoid derivative highlight the QTL associated with CYP76AD1 and UGT91C1, respectively. Principal component analysis of transiently overexpressed (OX) lines compared to empty vector (EV) controls in *C. quinoa* demonstrates clear metabolic differentiation. Both genes were functionally validated, as shown by the boxplots and chromatograms of betalamate and the addition of two pentosides (pen) to kaempferol-glucuronide-pentoside (kaempferol-gluc–pen). Asterisks indicate significance \*\*\**p*<0.001 (Student’s *t*-test or Wilcoxon test based on data normal distribution). Abbreviations: TCA, tricarboxylic acid cycle; CB, Calvin-Benson cycle; C4H, cinnamate 4-hydroxylase; HCT4, hydroxycinnamoyltransferase 4; CHS, chalcone synthase; F3’H, flavonoid 3′-hydroxylase; HIDM, 2-hydroxyisoflavanone dehydratase; UGT, UDP-glycosyltransferase; L-DOPA, L-3,4-dihydroxyphenylalanine; DODA, L-DOPA 4,5-dioxygenases; CYP, cytochrome P450.

### Candidate genes underlying flavonoid, betalain, and saponin biosynthesis

To investigate the genetic architecture underlying metabolic biosynthetic pathways in quinoa, next, we focused on three key classes of specialized metabolites – flavonoids, betalains, and saponins (**Figs. 4, 5**). For flavonoids, we identified seven candidate genes involved in flavonoid and phenylpropanoid biosynthesis. On chromosome B2, a QTL shared between roots and seeds encompassed cinnamate 4-hydroxylase (C4H; *CQ014797*), which catalyzes the conversion of *trans*-cinnamate to *p*-coumarate, and hydroxycinnamoyltransferase 4 (HCT4; *CQ014802*), a key enzyme in lignin precursor formation. Additional QTL on chromosomes B8 and B9 contained one and four chalcone synthase (CHS) genes (*CQ011785*, *CQ034822*, *CQ034825*, *CQ034827*, *CQ034828*) associated with flavonoid accumulation in seeds and roots. The same region on B9 also included 2-hydroxyisoflavanone dehydratase (HIDM; *CQ034839*), which catalyzes the dehydration of isoflavones. In seeds, a major QTL on A5 harbored flavonoid 3′-monooxygenase (F3′H; *CQ031304*), while another on B5 (adjacent to TSARL1 and 2) comprised five UGT79B30 glycosyltransferases (*CQ004358*, *CQ004359*, *CQ004360*, *CQ004361*, *CQ004365*) previously implicated in flavonol glycosylation in *Glycine max* (Di *et al*., 2015). Similarly, a QTL on B6 contained three UGT79B6 genes (*CQ000822*, *CQ000823*, *CQ000824*), known for flavonol glycosylation in *Arabidopsis thaliana* (Yonekura-Sakakibara *et al*., 2014), co-localizing with one bHLH (*CQ000843*) and two ICE transcription factors (*CQ000847*, *CQ000848*), suggesting coordinated transcriptional control of flavonoid modification and accumulation. We further identified UGT91C1 (*CQ048026*; chromosome A1, 11 Mb), associated with a flavonoid derivative and annotated as a soyasaponin III rhamnosyltransferase orthologue in beetroot (*Beta vulgaris*). To provide more insight and functionally validate their roles in the biosynthetic pathways, transient overexpression in both *Nicotiana benthamiana* and *C. quinoa* resulted in a metabolic alteration, which was more pronounced in *C. quinoa* (**Fig. 4b, S13**). While no significant changes were observed in saponin levels (**Fig. S14**), differential effects were evident among flavonoids: kaempferol-glucuronide-pentoside accumulated more in the empty vector control, whereas kaempferol-glucuronide-3x(pentoside) was enriched in UGT91C1-OX lines, supporting a role for UGT91C1 in flavonoid glycosylation and decoration.

For betalains, we identified a QTL on B4 contained five L-DOPA 4,5-dioxygenases (DODA; *CQ020378*, *CQ020379*, *CQ020383*, *CQ020384*, *CQ020385*), which convert L-DOPA to betalamate, co-localizing with an extradiol ring-cleavage dioxygenase (*CQ020383*) and four UGTs (*CQ020393*, *CQ020394*, *CQ020395*, *CQ020397*), forming a tightly co-regulated cluster likely responsible for betalain biosynthesis and modification. Additionally, two major seed QTL on chromosomes A8 and A9, each containing CYP76AD1 (*CQ039659*, *CQ052677*, *CQ052678*, *CQ052696*, *CQ052698*, *CQ052697*), which catalyzes the sequential oxidation of L-tyrosine to L-DOPA and *cyclo*-DOPA. The A9 QTL, harboring the functionally cloned *CQ052697*, also included two bHLH transcription factors (*CQ052675*, *CQ052676*) at 53.7 Mb. This locus was associated with 2-decarboxybetanin and several unknown polar metabolites. By transient overexpression in *C. quinoa,* we were able to functionally validate its role in betalain biosynthesis (**Fig. 4b**). PCA revealed that PC1 accounted for 30.8% of the variation, clearly separating the overexpression line from the control, whereas in *N. benthamiana* no distinct separation was observed, even when co-infiltrated with DODA (**Fig. S13**). In total, 56 metabolites – including betalains, flavonoids, and saponins – were differentially abundant upon overexpression, of which 14 were betalains (**Fig. S14**). Notably, betalamate levels were significantly higher in the overexpression line compared to the empty vector control (**Fig. 4b**), confirming the gene’s role in betalain biosynthesis.

For the triterpenoid saponin biosynthetic pathway, we identified four genes directly involved in saponin formation (**Fig. 5**). A prominent QTL shared between roots and seeds on chromosome B2 harbored CYP716A15 (*CQ014759*), a β-amyrin 28-monooxygenase implicated in oleanolate synthesis in *Vitis vinifera* (Fukushima *et al*., 2011). Another QTL on chromosome A8 (33.4 Mb) contained CYP88D6 (*CQ039380*), encoding a β-amyrin 11-oxidase that introduces a hydroxyl group at the C-11 position of the triterpene scaffold. Here, two additional candidate genes were cloned and functionally validated through transient overexpression in *C. quinoa* (**Fig. 5b**). These included: (i) CYP72A154 (*CQ049330*; chromosome A1, 47 Mb), which catalyzes the subsequent oxidation at C-30 to produce serijanate, and (ii) a soyasapogenol B glucuronide galactosyltransferase (SGT; *CQ012413*; chromosome B8, 67.8 Mb), responsible for the galactosylation of soyasapogenol B. Both genes conferred discrete metabolic signatures upon overexpression, as evidenced by PCA in *N. benthamiana* (**Fig. S13**) and an even more pronounced effect in *C. quinoa* (**Fig. 5b**). Their expression significantly altered the abundance of 76 and 45 metabolites, respectively, including betalains, flavonoids, and saponins (**Fig. S15**), with serijanate and soyasaponin I being significantly increased in the CYP72A154-OX and SGT-OX lines relative to empty vector controls (**Fig. 5b**).

**Figure 5.**
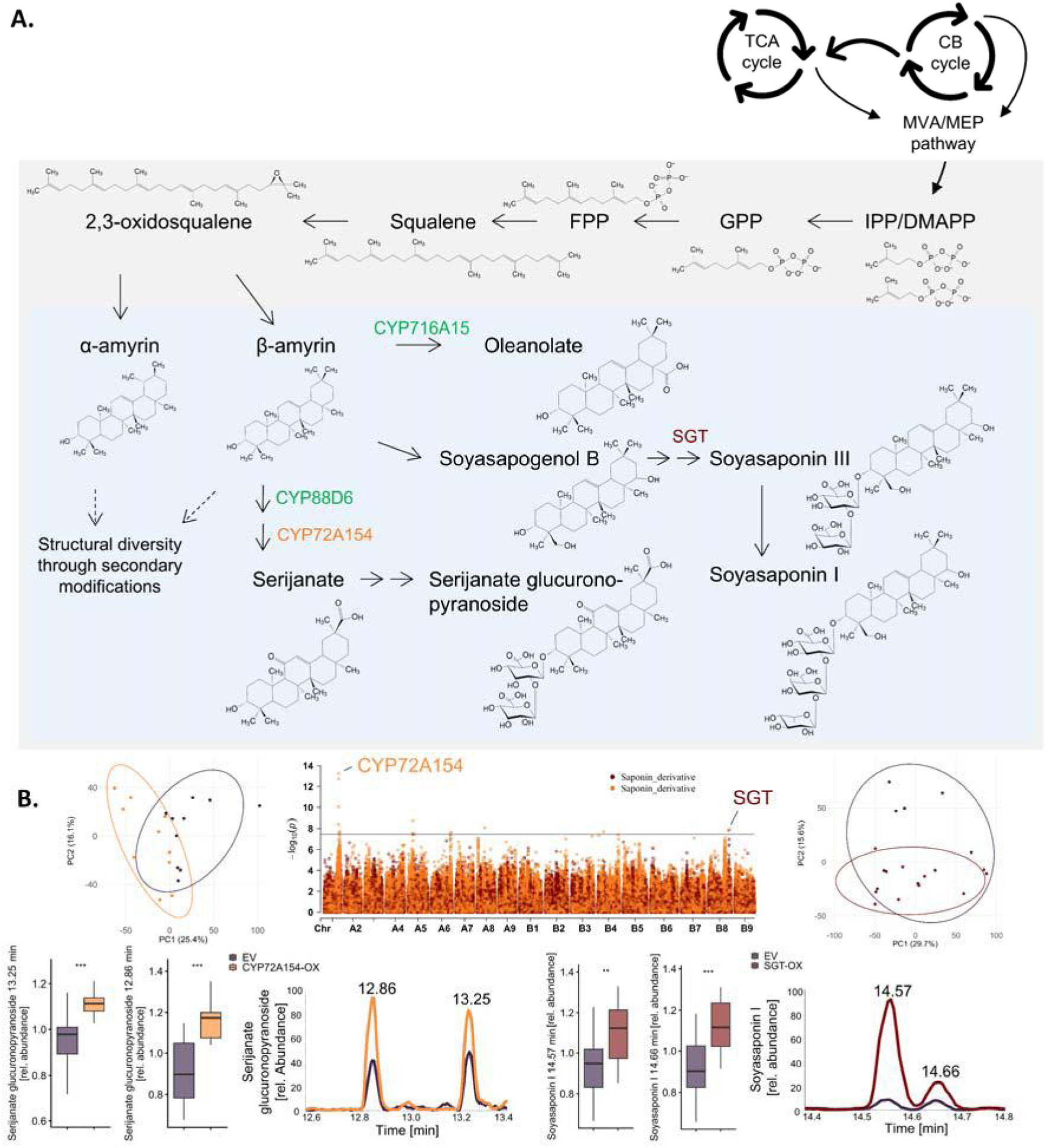
Refinement of the saponin pathway in *C. quinoa*. (**A**) The saponin pathway originates from the mevalonate (MVA) and methylerythritol phosphate (MEP) cytosolic and plastidic pathways, respectively, and proceeds through multiple condensation reactions of isopentenyl pyrophosphate (IPP)/ dimethylallyl pyrophosphate (DMAPP) forming the saponin precursor molecules α- and β-amyrin. Example of structural diversity through secondary modification are oleanolate which is synthesized through C-28 hydroxylation by CYP716A15, and serijanate glucuronopyranoside and soyasaponin by hydroxylation and glycosylation by soyasapogenol B glucuronide galactosyltransferase (SGT), CYP88D6, and CYP72A154. Candidate genes are marked in green, validated genes are colored in red and orange. (**B**) Overlay Manhattan plots for a representative saponin derivatives highlight the QTL associated with CYP72A154 and SGT. Principal component analysis of transiently overexpressed (OX) lines compared to empty vector (EV) controls in *C. quinoa* demonstrates metabolic differentiation. Both genes were functionally validated, as shown by the boxplots and chromatograms of serijanate glucuronopyranoside and soyasaponin I. Asterisks indicate significance \*\**p*<0.01, \*\*\**p*<0.001 (Student’s *t*-test or Wilcoxon test based on data normal distribution). Abbreviations: TCA, tricarboxylic acid cycle; CB, Calvin-Benson cycle; GPP, geranyl pyrophosphate; FPP, farnesyl pyrophosphate; UGT, UDP-glycosyltransferase; CYP, cytochrome P450.

While these functionally validated loci provide a foundation for understanding saponin biosynthesis in quinoa, it is likely that additional genes among the 219 candidates identified across 58 QTL contribute to the biosynthesis, transport, or modification of flavonoids, betalains, and saponins. These findings underscore both the complexity of specialized metabolism in quinoa and the current limitations of genome annotation for this species.

### Multi-omic validation of drought-responsive candidate genes

To further support candidate gene identification from our GWAS, we conducted a drought experiment using seven quinoa accessions selected based on seed metabolic diversity (**Figs. 2e, S14**). Plants were grown under both greenhouse and polytunnel conditions, experiencing temperatures from 6.7 to 44.9 °C and a maximum photosynthetically active radiation (PAR) of 1,717 µmol m⁻² s⁻¹ (**Fig. S15**). Drought stress caused significant reductions in growth, fresh and dry biomass, and altered photosynthetic activity in some lines (**Figs. S16–S20**).

Integration of multi-omics datasets revealed tissue- and environment-specific drought signatures (**Table S13**). Metabolomics (LC–MS polar/apolar and GC–MS primary metabolites) separated polytunnel-grown leaves from greenhouse leaves, roots, and seeds, explaining 36.5% of variance, while greenhouse leaves separated from other tissues by 24%, and seeds diverged from roots along PC3 (11.7%; **Fig. 6a**). Under drought, leaves and roots accumulated sugars and organic acids, while flavonoids, dipeptides, and triacylglycerides (TAGs) were differentially abundant in leaves, cholesterols, saponins, and phospholipids (PLs) in roots, and betalains, saponins, and PLs in seeds (**Figs. S21–S23**). PCA indicated that secondary metabolites primarily drove tissue separation, rather than primary metabolites or lipids (**Fig. S24a**). Proteomics of greenhouse leaves and roots explained 59.8% of variance along PC1, with 13.7% within-tissue variation (**Figs. 6b, S24b**), whereas transcriptomics separated roots and leaves from seeds (31.9% along PC1) and further distinguished tissues along PC2 (23.4%; **Figs. 6c, S24c**).

**Figure 6.**
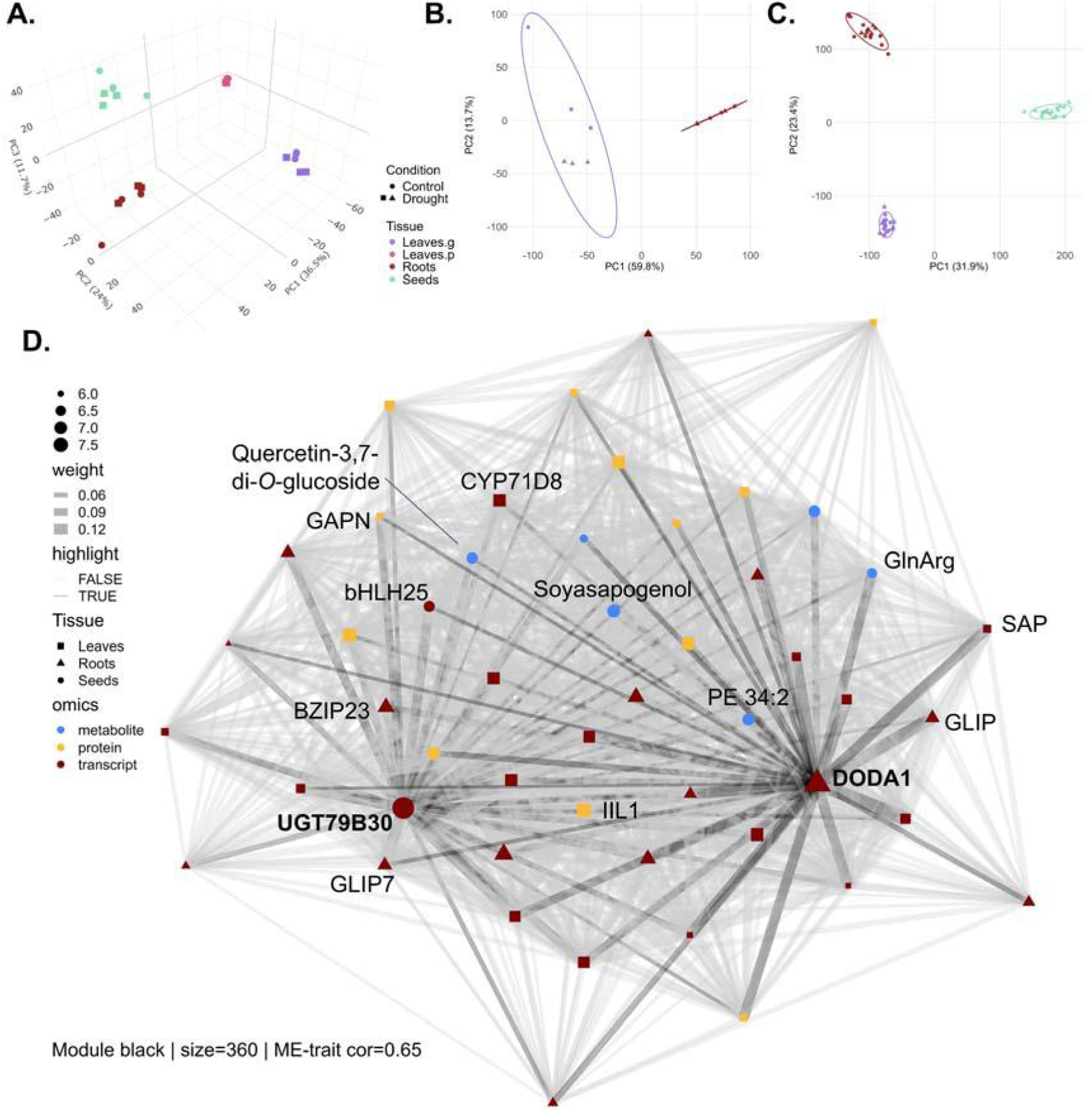
Multi-omic diversity of three quinoa accessions across tissues. (**A**) 3D principal component analysis (PCA) of primary metabolites, secondary metabolites, and lipids in leaves (p = polytunnel, g = greenhouse), roots, and seeds of accessions CHEN-384, LM2, and PI-665276. (**B**) PCA of proteomic profiles from greenhouse-grown leaves and roots of the same accessions. (**C**) PCA of transcriptomes from leaves, roots, and seeds under control and drought conditions. (**D**) Top 50 features of black module of weighted gene co-expression network analysis (WGCNA) of 15,809 metabolic, transcriptomic and proteomic features across leaf, root and seed tissues of CHEN-384, LM2, and PI-665276 identifying 46 modules of four block-wise clusters. DODA1 and UGT79B30 were identified as candidate genes in GWAS analysis, meaningful correlated features were highlighted. CYP, cytochrome P450; GAPN, NADP-dependent glyceraldehyde-3-phosphate dehydrogenase; GLIP, GDSL esterase/lipase; BZIP, basic leucine zipper; bHLH, basic helix-loop-helix; SAP, sterile apetala; GlnArg, glutamyarginine; PE, phosphatidylethanol; IIL, 3-isopropylmalate dehydratase large subunit.

WGCNA identified 46 modules (**S25–S26, Table S14**), six of which showed the strongest associations with drought response: turquoise (1,663 traits, r = 0.92, |GS| = 0.60), black (360 traits, r = 0.65, |GS| = 0.51), tan (183 traits, r = 0.69, |GS| = 0.49), purple (238 traits, r = 0.65, |GS| = 0.48), skyblue (86 traits, r = 0.72, |GS| = 0.46), and greenyellow (184 traits, r = 0.61, |GS| = 0.44). Functional enrichment revealed distinct biological signatures: the turquoise module was enriched in transcripts and proteins linked to cellular and metabolic processes, strongly correlated with lipids, dipeptides, and saponins; black and tan modules were dominated by proteins associated with metabolic and oxidoreductase activity and molecular/cellular processes, co-varying primarily with saponins, lipids, and dipeptides; the purple module combined lipids, saponins, and flavonoids with transferase-related functions; the skyblue module featured seed metabolites including phenylalanine, quercetin-3,4-di-*O*-glucoside, and miraxanthin-I, associated with oxidoreductase activity and energy metabolism; and the greenyellow module comprised only proteomic and transcriptomic features enriched in energy generation and small-molecule metabolism (**Figs. S27–S31**).

The black module included two candidate genes identified via GWAS: DODA1 (*CQ020379*) in roots and UGT79B30 (*CQ004358*) in seeds (**Fig. 6d**). Both genes showed co-expression with the seed metabolites soyasapogenol, glutamyarginine, phosphatidylethanol, and quercetin-3,7-di-*O*-glucoside, as reflected by topological overlap in the WGCNA network. In addition, several other network members exhibited tissue-specific co-expression: leaf proteins NADP-dependent glyceraldehyde-3-phosphate dehydrogenase (*CQ055725*) and 3-isopropylmalate dehydratase large subunit (*CQ021690*); root transcripts GDSL esterase/lipase (*CQ011072*, *CQ015414*) and basic leucine zipper 23 (*CQ021500*); seed transcript bHLH25 (*CQ015892*); and leaf transcripts CYP71D8 (*CQ012278*) and the transcriptional regulator sterile apetala (*CQ050157*).

Integration of these multi-omic datasets validated some GWAS candidate genes and highlights additional genes and metabolites for future functional studies, providing a framework to dissect the regulatory networks underlying drought tolerance in quinoa.

## Discussion

### The quinoa diversity panel reveals high genomic and metabolic variation

The re-sequenced *C. quinoa* diversity panel, comprising 603 accessions, was subjected to integrated genomic and metabolic analyses followed by GWAS. This panel included 121 accessions previously profiled for seed metabolites (Tabatabaei *et al*., 2022) and 310 accessions earlier characterized by sequencing, population structure, and GWAS-based QTL mapping (Patiranage *et al*., 2022). Similarly, population structure clusters could be assigned to the altitude of the accessions collection side (**Figs. 1a, S1**). Genome-wide positive Tajima’s D values indicate balancing selection, consistent with genetically distinct subgroups and potentially multiple domestication events (**Fig. 1d**; Carlson et al., 2005). This interpretation is supported by high inbreeding coefficients (F > 0.5) in 92% of accessions (**Fig. S1d**). Notably, genomic variation clearly separated highland accessions from those originating in lowland (**Fig. 1c**), suggesting two centers of domestication.

This genomic stratification was mirrored in the metabolic landscape, particularly among secondary metabolites, with saponins emerging as the most prominent discriminators, corroborating earlier observations (Tabatabaei *et al*., 2022). Tissue-specific profiling revealed root saponins, leaf flavonoids, and both seed saponins and flavonoids as the most abundant metabolite classes (**Fig. 2**), consistent with their ecological roles in defense and photoprotection (Kuljanabhagavad *et al*., 2008; Wang *et al*., 2019). In seeds, TAGs, sphingolipids, and PLs dominated the lipidome, reflecting their conserved functions in energy storage, membrane stability, and desiccation tolerance (Colin and Jaillais, 2020; Yang and Benning, 2018; Michaelson *et al*., 2016). The highest CV values were observed for root and seed flavonoids and saponins, highlighting their dynamic evolutionary turnover under both natural and artificial selection (Kuljanabhagavad *et al*., 2008; Wang *et al*., 2019; Yu *et al*., 2022). This variability likely reflects local ecological adaptation as well as human-driven selection for taste, palatability, and reduced anti-nutritional content.

We classified quinoa accessions as bitter or sweet based on a discriminant compound eluting at 4.68 minutes. Notably, this feature appears to belong to the flavonoid class rather than to saponins, aligning with recent work questioning the long-held assumption that saponins are the principal drivers of quinoa bitterness. Instead, flavonoids and other polyphenols have been proposed as major contributors to bitter taste in quinoa (Song *et al*., 2024). Moreover, growing evidence shows that di- and oligopeptides can elicit bitter or umami sensations (Kohl *et al*., 2013; Maehashi *et al*., 2008), indicating that quinoa taste is likely shaped by a complex interplay of multiple metabolite classes rather than by saponins alone. These findings underscore that breeding strategies aimed at reducing bitterness should take flavonoid, polyphenol, and dipeptide composition, alongside saponin content, into account.

Consistent with previous analyses of quinoa leaf drought responses (Huan *et al*., 2022), we observed a significant downregulation of trehalose in roots under drought stress considering all accessions (**Fig. S24**). Glucose, however, displayed contrasting dynamics, showing leaf-specific upregulation in our study while being downregulated in variety Dianli 129, indicating context-dependent sugar signaling roles in stress responses. These divergent patterns underscore the pleiotropic functions of sugars in drought signaling and their integration within broader metabolic regulatory networks. Notably, seed metabolism remained comparatively stable, in agreement with findings in other species (Leonova *et al*., 2020), although recent evidence in quinoa highlights cultivar-specific transcriptional and metabolic shifts in sugar metabolism under drought (Wang *et al*., 2024). The subdued metabolic response in seeds cannot be attributed to insufficient drought exposure (applied for two weeks, interrupted for two weeks, and then resumed until harvest), but instead may indicate metabolic buffering, developmental canalization, or delayed stress signaling in this tissue. However, in all cases, a genotype-specific difference in metabolome, proteome and transcriptome could be observed.

Our study provides a comprehensive view of the drought regulatory network in quinoa, integrating multi-omic and multi-tissue data from three diverse varieties. This approach not only revealed novel metabolites and gene candidates underpinning drought tolerance but also highlights the power of combining genomic, transcriptomic, proteomic, and metabolomic datasets to dissect complex traits. While the primary aim here was to support candidate gene identification in the GWAS framework, these results lay a foundation for future functional studies and for breeding quinoa varieties with enhanced resilience to environmental stress.

### GWAS identifies QTL linked to adaptation, secondary metabolism, and seed pigmentation

Among QTL associated with elevation – previously shown to explain PC1 variation in a subset of 296 genotypes (Patiranage *et al*., 2022) – candidate genes included amongst the 35 identified genes *MAP1A*, *Arginase 2*, an ERF transcription factor, and *EER5*, all involved in hypoxic stress responses crucial for high-altitude adaptation in Andean crops like quinoa (Galwey, 1992; Giuntoli and Perata, 2018; Loreti and Perata, 2020). Notably, a cluster of 21 SAUR-like auxin-responsive genes – implicated in plant development and hypoxia responses (Hsu *et al*., 2011; Stortenbeker and Bemer, 2019) – was also linked to elevation. This aligns with known altitude-related growth constraints such as reduced CO₂ partial pressure and increased UV radiation (Korner and Diemer, 1994; Gale, 2004).

The saponin regulators *TSARL1* and *TSARL2* (on chromosome B5), previously identified by Jarvis et al. (2017), were validated in this study using the LC-MS. GWAS of metabolite levels identified a set of candidate genes with direct functional relevance to secondary metabolism, including 26 UGTs, 20 transporters, 34 cytochrome P450s, 34 transcription factors (including nine bHLH), and 23 sugar-, methyl-, or acetyltransferases. These gene families collectively mediate biosynthetic tailoring, chemical modification (**Fig. 5**), transport, and transcriptional control of specialized metabolites (Wang *et al*., 2019), providing a mechanistic explanation for the observed variation in flavonoid and saponin content. Their association with metabolic traits highlights the power of integrating genomic variation with biochemical phenotypes to uncover regulatory architecture.

Four candidate genes were functionally implicated in the saponin biosynthetic pathway, including CYP88D6 and CYP72A154, previously characterized in serijanate glucuronopyranoside (also known as glycyrrhizin) biosynthesis (Seki *et al*., 2008; Seki *et al*., 2011; Biazzi *et al*., 2015), as well as a SGT as glycosyltransferase responsible for converting soyasapogenol B to soyasaponin III (Shibuya *et al*., 2010). Serijanate glucuronopyranoside, widely used as a natural sweetener, also exhibits anti-inflammatory properties (Hayashi et al., 2003). Moreover, group B saponins such as soyasaponin I, which lack the C22 moiety present in group A saponins, are less bitter and considered health-promoting (Yu *et al*., 2022). While soyasaponins have been considered Fabaceae-specific (Yu *et al*., 2022), our results suggest that structurally similar group B saponins may also occur in Amaranthaceae, highlighting the need for further structural and functional characterization.

Flavonoids and betalains are key specialized metabolites in quinoa with important ecological and nutritional functions. Flavonoids contribute to UV protection, defense, and pollinator attraction (Wang *et al*., 2019; Zhuang *et al*., 2023), yet their biosynthesis and modification remain poorly explored in *C. quinoa* (Qian *et al*., 2023; Jiang *et al*., 2025). Among the 21 candidate genes implemented in the flavonoid pathway in our study, UGT91C1 mediates the glycosylation of a kaempferol derivative, a modification that enhances solubility and stability. Notably, UGT91C1 also glycosylates diterpenes such as steviol in rice and *Arabidopsis* (Zhang *et al*., 2021; Huang *et al*., 2021), highlighting its broad substrate versatility and potential role in modulating diverse secondary metabolites.

Betalains, pigments unique to the Caryophyllales order, provide antioxidant and anticancer benefits (Escribano *et al*., 2017; Timoneda *et al*., 2019) and are increasingly valued as natural food colorants. Engineering betalain accumulation in crops has been demonstrated in tomato, potato, and tobacco via targeted expression of biosynthetic genes (Nakatsuka *et al*., 2013; Grützner *et al*., 2021). Our identification and functional validation of CYP76AD1 in quinoa provide robust targets for the bioengineering of betalain-enriched quinoa varieties, opening opportunities to enhance both nutritional quality and market value of this increasingly popular crop.

Integration of multi-omic drought-response analyses revealed additional candidate genes through correlation-based approaches, providing a framework for the systematic identification of metabolic genes and their co-regulators that contribute to stress resilience.

Overall, our findings provide a roadmap for breeding programs aimed at improving nutritional quality, stress tolerance, and palatability. More broadly, this study demonstrates the utility of combining genomic and metabolomic approaches to dissect complex traits in an orphan crop with high agronomic and ecological relevance.

## Conclusion

Our quinoa diversity panel enabled the identification of 219 candidate genes across 58 high-confidence QTL, while also providing a scalable framework for uncovering additional loci linked to the secondary metabolome in leaves, roots, and seeds. By integrating multi-method GWAS with stringent SNP selection, we prioritized robust candidates, yet variants detected by fewer methods may reveal additional biologically relevant associations. By releasing the full mapping data for numerous unknown metabolites – including mass-to-charge ratios (*m/z*) and retention times – we furthermore enable researchers to perform their own annotations and discover QTL for metabolites not captured in our study. This dataset therefore functions both as a curated candidate-gene resource and a flexible toolkit for expanding the genetic and metabolic understanding of quinoa.

## Material and Methods

### Plant materials, sequencing, and SNP identification

Seeds of 603 Chenopodium quinoa accessions were provided by Prof. Mark Tester from the King Abdullah University of Science and Technology (KAUST). The re-sequenced 603 accessions were used in study (Stanschewski et al., submitted) of which 296 accessions were previously published (Patiranage *et al*., 2022). Whole-genome sequencing (WGS) was performed following (Jarvis *et al*., 2017). In total, 13,743,142 polymorphic sites were identified by SNP calling following https://github.com/IBEXCluster/Wheat-SNPCaller (Abrouk *et al*., 2020). After filtering using vcftools (https://vcftools.github.io/man_latest.html) eliminating SNPs with 90% of missing data, considering 5% minor allele frequency and 30 as quality score threshold 1,450,189 high confidence SNPs remained and used to conduct the genomic and GWAS analysis on the full 603 accessions. For the GWAS of quinoa leaf and root samples, triplicates of 166 accessions were germinated using 0.1% agarose containing 500 µM gibberellic acid A3 and stratified for two days at 4 °C. Plants were grown using sandy soil and 1% NovaTec® classic fertilizer in the greenhouse and leaf and root tissues were harvested after 4 weeks of growth.

### Drought experiment: Growth conditions and sampling

Seeds were surface sterilized following the protocol of Hesami et al. (2018). Briefly, seeds were incubated in 70% ethanol for 15 seconds, rinsed with double-distilled water (ddH₂O), and then treated for 5 minutes with 20% sodium hypochlorite containing 0.001% Tween-20. After four washes with ddH₂O, seeds were stratified for two days at 4 °C in 0.1% agarose supplemented with 500 µM gibberellic acid A₃. Seedlings were grown in sandy soil supplemented with 1% NovaTec® Classic fertilizer under controlled conditions in both the polytunnel and greenhouse starting in May. The following quinoa accessions were grown in the polytunnel: Ames-13760, PI-665276, LM2, CHEN-199, CHEN-384, and D-12393. In the greenhouse, the accessions CHEN-199, D-12165, Ames-13760, CHEN-384, PI-665276, and LM2 were cultivated. Drought stress was applied by modifying the irrigation regime: in the polytunnel, plants were watered every second day instead of daily; in the greenhouse, the amount of water was reduced by half every other day. Irrigation volumes were adjusted based on the full turgor of control plants. After two weeks of drought stress, leaf and root samples were harvested from quinoa accessions in the greenhouse at five weeks of age. At the same time, fresh and dry weight of the above-ground tissue was recorded. Leaf samples from polytunnel-grown plants were harvested at the same developmental stage. Photosynthetic parameters were recorded using the MultispeQ device (PhotosynQ), and plant height was measured. Drought stress was then paused for two weeks and subsequently re-applied. Plants were then grown to maturity under continued drought conditions, and final measurements of plant height and dry weight were taken.

### Genomic analysis

For the analysis of the genomic diversity Tassel 5 (Bradbury *et al*., 2007) and vcftools (Danecek *et al*., 2011) was used. Tajima’s D plot was designed using the R package ‘CMplot’. The phylogenetic tree was designed using iTOL (https://itol.embl.de/). The population structure of the GWAS population was determined using the software fastSTRUCTURE v1.0 (Raj *et al*., 2014) was utilized with K 1 – 15. The optimal K value was determined by fastSTRUCTUREs chookeK.py implementation and visualized with the R package ‘pophelperShiny’.

### Metabolite profiling using UHPLC-MS

Metabolites were extracted using a methanol–chloroform–water biphasic protocol based on (Bligh and Dyer, 1959) and adapted for plant tissues (Weckwerth *et al*., 2004). In brief, 50 mg of dried seeds were ground for 1 min at 23 Hz, after addition of 700 µl of 100% methanol spiked with 0.4 µg/ml isovitexin, the material was ground for another minute at 17 Hz, shaken for 10 min at 28 °C, sonicated for 5 minutes and 500 µl of chloroform (containing 0.5 µg/ml phosphocholine 17:0) was added before adding 700 µl of ddH_2_O. The extract was centrifuged for 5 minutes at 10,000 *g* and 100 µl of the semi-polar phase was transferred to LC-MS vials for analysis. For lipid analysis 300 µl of the apolar phase was concentrated and resuspended in 200 µl acetonitrile: isopropanol [7:3]. After vortexing the samples were ultrasonicated for 2 or 5 min for polar and lipid extracts, respectively, centrifuged (5 min, room temperature, 10,000 *g*) and 100 µl of the supernatant was used for ultra-high-pressure liquid chromatography electrospray ionization mass spectrometry (UHPLC-ESI-MS).

Analysis of semi-polar metabolites was conducted on a UHPLC-ESI-MS machine as described previously (Alseekh *et al*., 2015). For semi-polar compounds a technical injection was performed in order to exclude machine errors. In brief, the UHPLC system was equipped with an HSS T3 C18 reverse-phase column (100 × 2.1 mm internal diameter, 1.8 μm particle size; Waters) that was operated at a temperature of 40°C. The mobile phases consisted of either 0.1% formic acid in water (solvent A) and 0.1% formic acid in acetonitrile (solvent B) for semi-polar compounds and of 1% 1 M NH_4_-acetate, 0.1% acetic acid in UHPLC grade water (solvent A) and 1% 1 M NH_4_-acetate, 0.1% acetic acid in UHPLC grade acetonitrile:isopropanol [7:3] (solvent B) for the lipophilic fraction. The flow rate of the mobile phase was 400 μL/min, and 3 μL of the sample was loaded per injection. The UHPLC instrument was connected to an Exactive Orbitrap focus (Thermo Fisher Scientific) via a heated electrospray source (Thermo Fisher Scientific). The spectra were recorded using full-scan in positive ion-detection mode, covering a mass range from m/z 100 to 1,500 with an ESI approach (capillary conditions at 3 kV, 200 °C; drying gas at 350 °C; sheet and auxillary gas flow at 60 U and 20 U, respectively; skimmer and tube lens at 25 V and 130 V, respectively) (Perez de Souza *et al*., 2019).The resolution was set to 70,000 and the maximum scan time was set to 250 ms. The sheath gas was set to a value of 60 while the auxiliary gas was set to 35. The transfer capillary temperature was set to 150 °C while the heater temperature was adjusted to 300 °C. The spray voltage was fixed at 3 kV, with a capillary voltage and a skimmer voltage of 25 V and 15 V, respectively. MS spectra were recorded from 0 to 19 minutes of the UPLC gradient. Processing of chromatograms, peak detection, and integration was performed using RefinerMS (version 5.3; GeneData).

### Metabolite profiling using GC-MS

Primary metabolic features were extracted as described above. An aliquot of 100 μl of the polar phase was dried in a SpeedVac concentrator and stored at −20 °C. For derivatisation of the primary metabolites, 55 μl of 30 mg/ml methoxyaminhydrochloride in pyridine was added to the concentrate and incubated (120 min, 37 °C, 950 rpm). Following, 110 μl N-Methyl-N-(trimethylsilyl) trifluoracetamid (MSTFA) was added, shaken (30 min, 37 °C, 950 rpm) and 100 μl was used for GC-MS analysis. GC-MS was performed according to (Lisec *et al*., 2006). Separation took place by injection of 1 μl of samples in a splitless mode with helium as carrier gas with a flow set to 2 ml min^-1^ (constant with electronic pressure control) by usage of an autosampler setup. For high-abundance metabolites (glucose, fructose, saccharose, malic acid, glutamic acid), injection was executed in a split mode with a 1:20 ratio. For metabolite profiling, a 30-m MDN-35 capillary column with an isothermal temperature program (2 min at 80 °C, 15 °C per minute increase to 330 °C and 6 min at 330 °C, followed by rapid cooling; transfer line temperature at 250 °C matching ion source conditions). The instrument recorded 20 scans per sec in a mass range of m/z 70 to 600 (remaining chromatography with a 170 sec solvent delay with filaments turned off). The manual mass defect was set to 0, and detector voltage to 1700 – 1850 V with a filament bias current at −70 V. Processing of chromatograms, peak detection, and integration was performed using RefinerMS (version 5.3; GeneData).

### Compound annotation, filtering, normalization, and transformation

Metabolite identification and annotation were performed using in-house standard compounds, tandem MS (MS/MS) fragmentation, and metabolomics databases. When using the in-house reference compound library, we allowed for a 5-ppm mass error and a dynamic retention-time shift of 0.1. Additionally, we conducted a literature and integrated metabolic data analysis from previous studies, for example, betalain annotation from Dini et al., 2006; Xie and Chen, 2021; Escribano et al., 2017; saponin annotation from Madl et al., 2006; Tabatabaei et al., 2022; Escribano et al., 2017; and flavonoids based on Wang et al., 2019. Metabolite data is reported following updated standards for metabolite reporting (Alseekh, et al., 2021).

Data were filtered removing features with quality controls (QCs) that contain the pooled samples > 5%, the coefficient of variation of QCs > 60%, missing values of samples > 80% and the signal/blank ratio < 10%. Only features from 1 – 17 minutes were used. Data were imputed by 10% of the minimum value of the trait before normalization based on the sample weight, the internal standard isovitexin, the feature mean, the QCs per batch and a day mean standardisation. For GWAS, samples were log_2_ transformed.

### Metabolite network analysis

Metabolite networks were constructed from normalized metabolite abundances across accessions, with compounds annotated by tissue and class. Pairwise Pearson correlations between metabolites were calculated, and edges were retained for correlations |r| > 0.7 with *p* < 0.05. Networks were built using ‘igraph’, with node degree calculated to identify hub metabolites. Networks were visualized using ‘ggraph’ with the Kamada–Kawai layout. All analyses were performed in R (4.3.2.) using the ‘tidyverse’, ‘igraph’, ‘ggraph’, and ‘viridis’ packages.

### GWAS analysis

To screen for significant associations using GWAS the R package ‘rMVP’ (Yin *et al*., 2021) was used employing Haseman-Elston regression (HE) as variance component method and as method for genotype binning, factored spectrally transformed linear mixed model (FaST-LMM) was utilized. To correct for population structure effects with a kinship matrix and three PCs as fixed effects. Single-locus MLM (mixed linear model) was used as an association testing method for the identification of significant associations between genetic variants (SNPs) and the phenotype of interest. Missing values were excluded from the GWAS. We used the Bonferroni correction to adjust the significance threshold for multiple comparisons. Significant compounds were reanalyzed using efficient mixed-model association (EMMA) as variance component and binning method, and MLM and farmCPU for association testing method. Candidate genes were identified by scanning the genetic area ± 100 kb of the lead SNP and determining the biological relevance by including the results of OrthoFinder (Emms and Kelly, 2015; Emms and Kelly, 2018). Protein sequences were obtained from the Joint Genome Institute (https://data.jgi.doe.gov/; *Arabidopsis thaliana* - TAIR10, *Spinacia oleracea* - Spov3, *Amaranthus hypochondriacus* - v2.1, *Arachis hypohaea* cv. Tifrunner, *Beta vulgaris* - EL10_1.0, *Cicer arietinum* - v1.0, *C. quinoa* - v1.0, *Chlamydomonas reinhardtii* - v5.6, *Crocus sativus* - v1.0, *Gossypium hirsutum* - v2.1, *Glycine max* - Wm82.a2.v1, *Lupinus albus* - v1, *Lens culinaris* - v1, *Medicago truncatula* - Mt4.0v1, *Oryza sativa* - v7.0, *Phaseolus vulgaris* - v2.1, *Sorghum bicolor* - v3.1.1, *Solanum lycopersicum* - ITAG4.0, *S. tuberosum* - v6.1, *Zea mays* - V4) and NCBI (https://www.ncbi.nlm.nih.gov/protein/; *S. lycopersicum* - ITAG4.1). In addition, transcriptomic data of previous quinoa studies were integrated for candidate gene identification (Rollano-Peñaloza *et al*., 2021; Liu *et al*., 2022; Ma *et al*., 2021; Zhao *et al*., 2022; Huan *et al*., 2022). Furthermore, the differential abundance of primary and secondary metabolites, lipids, proteins and RNA during the drought experiment was an additional proxy for the involvement in the secondary metabolome.

Common QTL across tissues were assigned in a ± 50 kb sliding window using ‘IRanges’ and ‘GenomicRanges’. The pleiotropic maps were designed with the circos representation Fuji plot (Krzywinski *et al*., 2009). Common QTL were assigned in a ± 50 kb window following the method previously described (von Steimker *et al*., 2024; von Steimker *et al*., 2025).

### Proteomics

Proteins from seeds, leaves and roots of LM2, CHEN-384 and PI-665276 grown under drought and control conditions were extracted using the methyl tert-butyl ether (MTBE) extraction method (Giavalisco *et al*., 2011; Salem *et al*., 2016). In brief, 50 mg of seed, leaf or root material was incubated for 10 minutes at 4 °C on an orbital shaker after adding 1 ml of −20 °C pre-cooled extraction solvent mixture 1 (MTBE/methanol [3:1]). After 10 minutes of ultrasonication 500 μl of extraction solvent, mixture 2 (methanol/water [3:1]) was added and samples were centrifuged (5 min, 4 °C, 11,000 g). The supernatant was removed and the pellets dried at room temperature. Proteins were digested using in-solution digestion protocol with modifications described in (Rappsilber *et al*., 2007; Thirumalaikumar *et al*., 2023). Briefly, proteins were digested using a trypsin/Lys-C mixture (Mass Spec Grade; Promega, V5073) according to the manufacturer’s instructions. Digested peptides were desalted on C18 SEP-Pak columns (Teknokroma, TR-F034000), in which peptides were eluted using centrifuge. The dried peptides were resolubilised and subsequently, analysed by LC-MS/MS using a 480-Exploris mass spectrometer coupled to an Ultimate U3000 nano LC (Thermo fisher, Scientific). The gradient and mass spec settings were kept as same as (Wagner et al. (2025). Raw data were processed using MaxQuant software (Cox and Mann, 2008). The Hela digest was used before and after the run to monitor the accuracy of the experiment.

### Transcriptomics

Total RNA of leaves and roots from the greenhouse and seeds from the polytunnel of LM2, CHEN-384 and PI-665276 grown under drought and control conditions was extracted using NucleoSpin® RNA plant kit (Macherey-Nagel). RNA sequencing was performed by Novogene (Munich, Germany). Total RNA quality was assessed prior to library preparation. Polyadenylated RNA was enriched from total RNA using poly-T oligo–attached magnetic beads, fragmented, and reverse-transcribed using random hexamer primers. Second-strand cDNA synthesis was performed using dUTP to generate strand-specific libraries. Libraries were constructed following standard Illumina protocols, including end repair, A-tailing, adapter ligation, size selection, PCR amplification, and purification. Library quality and concentration were assessed using Qubit fluorometry, quantitative PCR, and an Agilent Bioanalyzer. Sequencing was performed on an Illumina NovaSeq X Plus platform using paired-end sequencing (PE150). Raw reads were processed to remove adapter contamination, reads containing more than 10% ambiguous nucleotides, and reads with low base quality, as previously described (Yan *et al*., 2013). Clean reads were aligned to the reference genome using HISAT2, a graph-based splice-aware aligner optimized for RNA-seq data (Mortazavi *et al*., 2008). Transcript assembly and novel transcript identification were performed using Cufflinks (Pertea *et al*., 2015). Gene expression levels were quantified based on mapped reads and normalized as fragments per kilobase of transcript per million mapped reads (FPKM) to account for sequencing depth and gene length (Mortazavi *et al*., 2008; Bray *et al*., 2016; Trapnell *et al*., 2010).

### Gene enrichment analysis

Gene Ontology (GO) is an acknowledged bioinformatics tool for representing gene product properties across all species through defined GO terms. The functions of genes and their products are represented by GO terms and effectively predicted through GO annotation (Ashburner *et al*., 2000). Functional enrichment analysis of significant proteins of drought versus control with cut-off criterion *p*-value < 0.05 and |log_2_ fold change| ≥ 1 were identified using the database g:Profiler (https://biit.cs.ut.ee/gprofiler/gost) with the gene IDs of *C. quinoa* - v1.0 matched through OrthoFinder.

### Weighted Gene Co-expression Network Analysis (WGCNA)

Co-expression networks were constructed using the ‘WGCNA’ R package. Multi-omics features (metabolites, proteins, transcripts) were clustered based on pairwise correlations, and modules of highly correlated features were identified using hierarchical clustering and dynamic tree cutting. Module eigengenes (first principal component) summarized module expression, and correlations with phenotypic traits were used to assess module significance. Hub features within each module were defined by high module membership (kME) and trait relevance (gene significance, GS). Network visualizations were generated in R using igraph and ggraph, with node color representing omics type and shape indicating tissue. Cytoscape-compatible node and edge tables were exported for downstream analysis and are available upon request.

### Statistical analysis

All data were analyzed in the R environment version 4.3.2. PCA was performed using ‘ggbiplot’. Heatmaps were designed using ‘pheatmap’ with median normalization and log_2_ transformation using Ward’s clustering method. Venn diagrams were computed using the R package ‘VennDiagram’, barplots, boxplots, volcano plots and histograms were designed using ‘ggplot2’ and ‘viridis’ for color selection. In order to compute volcano plots between conditions, data were tested for normal distribution and significance was tested by either Student’s t-test or Wilcoxon rank-sum test. Multiple comparison of accessions was performed either by two-way ANOVA and post hoc Tukey HSD test or Kruskal-Wallis test and post hoc Dunn’s test, ‘multcomp’ was utilized for computation of significance letters.

### Candidate gene cloning

Cloning of candidate genes was performed as described in (von Steimker *et al*., 2024). In brief, the total RNA from leaf, root and seed tissue was isolated using a NucleoSpin® RNA plant kit (Macherey-Nagel) according to the manufacturer’s instructions. First-strand cDNA was synthesized using 1.5 mg RNA and Prime Script™ RT reagent Kit with gDNA eraser (Takara) according to the manufacturer’s instructions. Full-length cDNA of quinoa seeds was amplified using 250 ng (**Table S15**). The entry clone was obtained through recombination of the PCR product with pDONR207 (Invitrogen). By LR recombination error-free clones were introduced into pK7FWG2 (Karimi *et al*., 2002). The transformation of three leaves of four *Nicotiana benthamiana* plants and five leaves of two *C. quinoa* plants of varieties CHEN-369 and Cauquenes was performed following (Zhang *et al*., 2020) and (Xiao *et al*., 2022) with *Agrobacterium tumefaciens* (AGL1) containing vector pBin61-p19, infiltrated with an OD600 of 0.5. DM6000B/SP5 confocal laser scanning microscope (Leica Microsystems, Wetzlar, Germany) was used for verification of expression. Metabolic shifts were analyzed after 3 days using 50 mg of agro-injected leaves and subjected to LC-MS analysis using chloroform:water:methanol extraction (Bligh and Dyer, 1959; Weckwerth *et al*., 2004) as described previously.

## Data availability

The authors declare that the data supporting the findings of this study are available within the paper, its supplementary information files, and upon request. Raw sequence data for the 610-accession quinoa diversity panel (of which 603 were used in this study) are distributed across two BioProjects in the NCBI Sequence Read Archive: PRJNA1357073 (Stanschewski et al., submitted) and PRJNA673789 (Patiranage *et al*., 2022).

## Acknowledgements

ARF and SA acknowledge BG16RFPR002-1.014-0003-C01 project, financed by the European Regional Development Fund through the Bulgarian Program for Research, Innovation, and Digitalisation for Smart Transformation (PRIDST) Operational Programme. S.A. acknowledges the NATGENCROP project: HORIZON-WIDERA-2022- TALENTS-01, No. 101087091. ELR, CS, NOS, and MAT acknowledge the baseline funding from King Abdullah University of Science and Technology (KAUST). JvS acknowledges the International Max Planck Research Schools for Molecular Plant Science.

## Author contribution

SA and MAT conceptualized the experiment. JvS and SA wrote the manuscript with input from all authors. ARF provided guidance on experimental strategy. ELR, CS, NOS, and MAT provided the germplasm and the sequencing data. AK filtered sequencing data. VT and AS performed proteomic analysis. JvS performed data analysis. JvS, RW and MM performed experiments.

## Supplementary Data

**Supplementary Data 1.** Tajima’s D across 603 *Chenopodium quinoa* accessions using 1.45 M SNPs in 100 kb nonoverlapping bins.

**Supplementary Data 2.** Inbreeding coefficient F and transition/transversion (Ts/Tv) ratio of 1.45 M SNPs across 603 *Chenopodium quinoa* accessions.

**Supplementary Data 3.** Annotated compounds in seed, leaf and root tissue.

**Supplementary Data 4.** Number of all annotated compounds and compounds with a significant association through genome-wide association study (GWAS).

**Supplementary Data 5.** Common compounds detected in seed, leaf and root tissue or pairwise in roots and seeds, leaves and seeds and leaves and roots.

**Supplementary Data 6.** Network analysis of annotated compounds.

**Supplementary Data 7.** Correlation of *C. quinoa* accessions assigned to sweet and bitter taste with metabolic compounds in seeds, leaves and roots.

**Supplementary Data 8.** Mapping results of seeds polar and apolar fraction, leaves and roots using either ≥2 common methods, ≥3, ≥4, ≥5 or 6 common genome-wide association study methods (BLINK, farmCPU, EMMA-MLM, HE-MLM, MLMM, CMLM).

**Supplementary Data 9.** Masterfile containing GWAS results, proteome, transcriptome and OrthoFinder results. In addition, previously published data are integrated.

**Supplementary Data 10.** Annotated metabolic features with a genomic association of leaves, roots and seeds using ≥4 common genome-wide association study methods.

**Supplementary Data 11.** Unannotated metabolic features with a genomic association of leaves, roots and seeds using ≥4 common genome-wide association study methods.

**Supplementary Data 12.** Candidate genes of leaves, roots and seeds identified using ≥4 common genome-wide association study methods.

**Supplementary Data 13.** Drought-responsive metabolites, transcripts and proteins identified in the accessions LM2, CHEN-384, and PI-665276 in leaves, roots and seeds.

**Supplementary Data 14.** Module ranking and feature meta data of WGCNA of drought multi-omics experiment.

**Supplementary Data 15.** Primers used for transient overexpression in *Nicotiana benthamiana* and *Chenopodium quinoa*.

---

**Figure S1.**
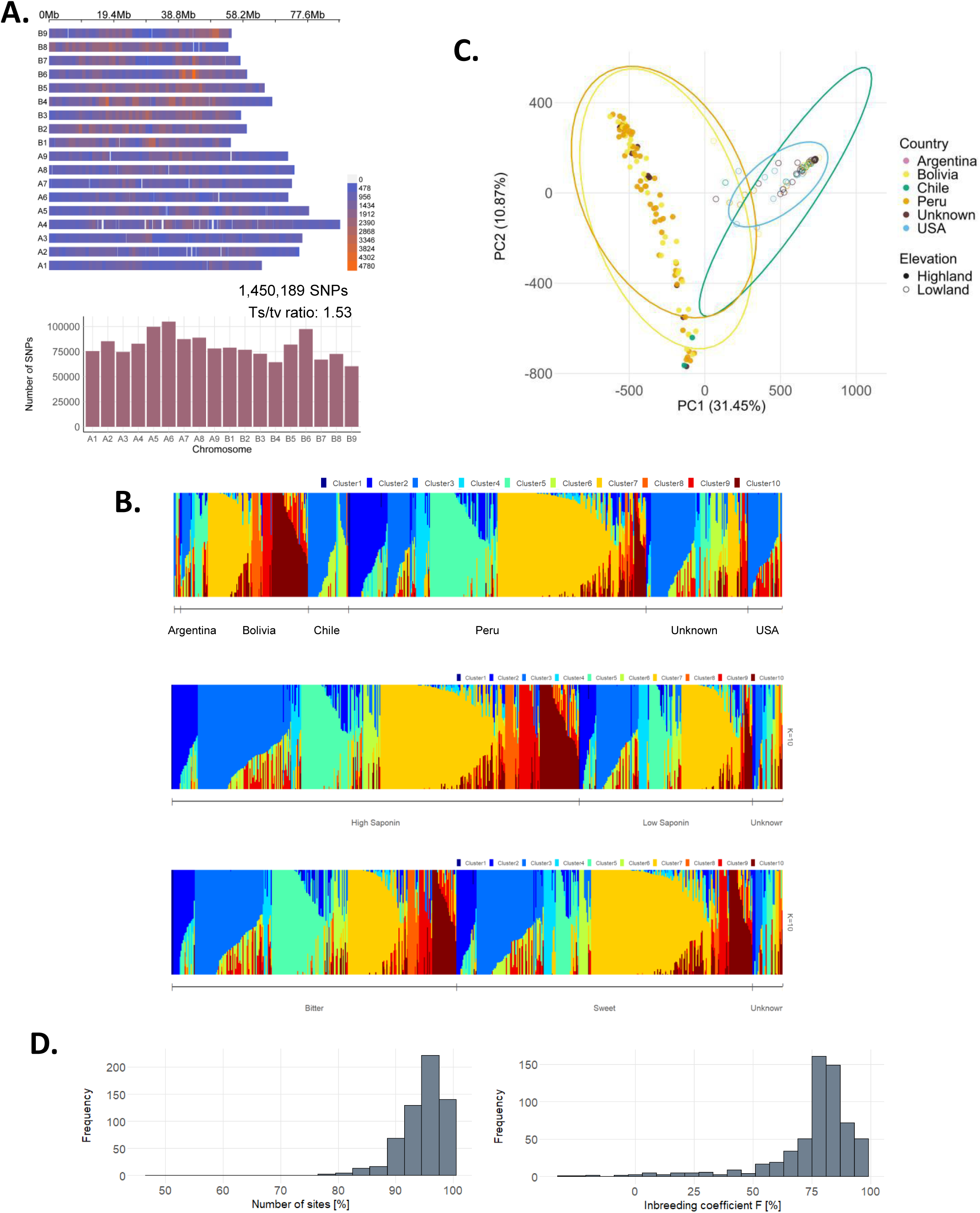
The quinoa diversity panel by its elevation. (**A**) SNP density in 1 Mb non-overlapping bins and number of SNPs per chromosome (60,417–104,938), based on 1,450,189 SNPs with a transition/transversion (Ts/Tv) ratio of 1.53. (**B**) The population structure plot grouped by elevation, sweet and bitter taste and average saponin content using K = 10 (model complexity that maximizes marginal likelihood). (**C**) Principal component analysis (PCA) of 166 quinoa accessions used for root and leaf GWAS colored by elevation and shaped by location. (**D**) Proportion of segregating sites and inbreeding coefficient (F [%]) indicate a high degree of inbreeding, consistent with a short breeding history typical of modern crops.

**Figure S2.**
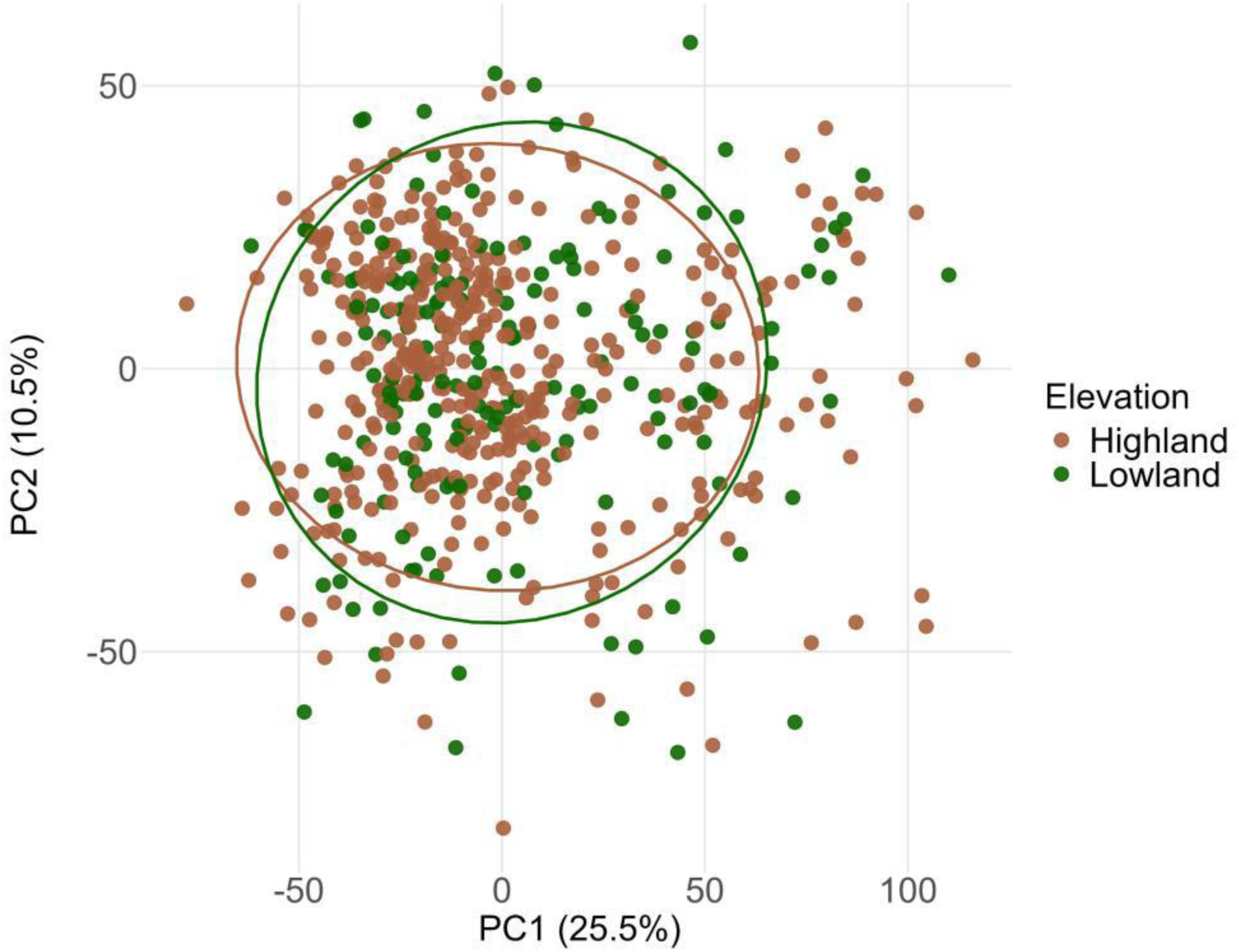
Principal component analysis of apolar seed compounds. 4,949 metabolic features from 573 accessions. Accessions are color-coded by elevation.

**Figure S3.**
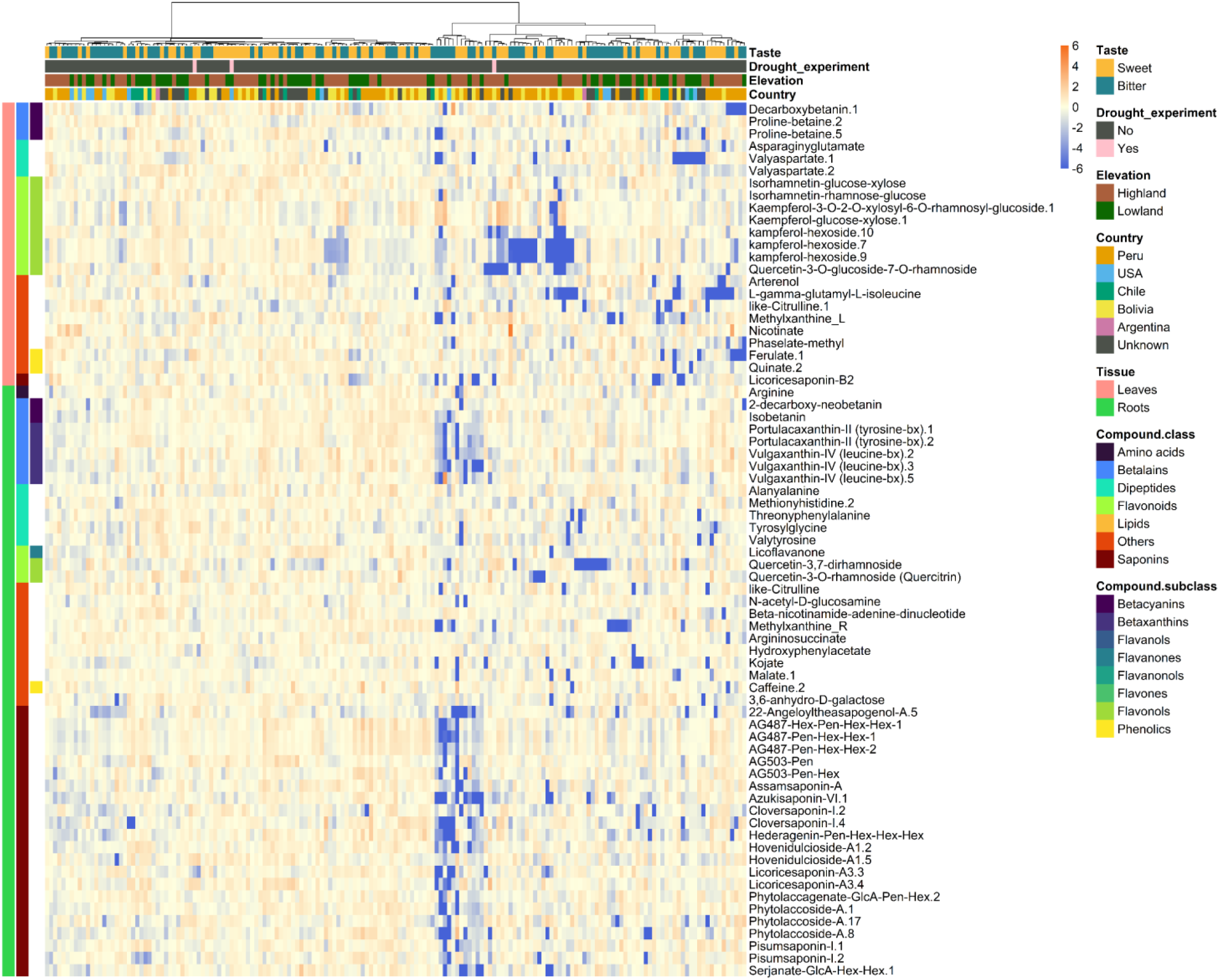
Secondary metabolic diversity of leaf and root tissue. Heatmap of 23 annotated leaf secondary metabolites and 48 root secondary metabolites with a genomic association. For leaves, 3 betalains, 3 dipeptides, 8 flavonoids, 1 saponin, and 8 other compounds, for roots, 1 amino acid, 7 betalains, 5 dipeptides, 3 flavonoids, 23 saponins, and 10 other metabolites. Accessions are color-coded by sweet and bitter taste, location and elevation of origin. Accessions which are used for the drought experiment are marked in pink (from left to right: Ames-13760, D-12393, PI-665276).

**Figure S4.**
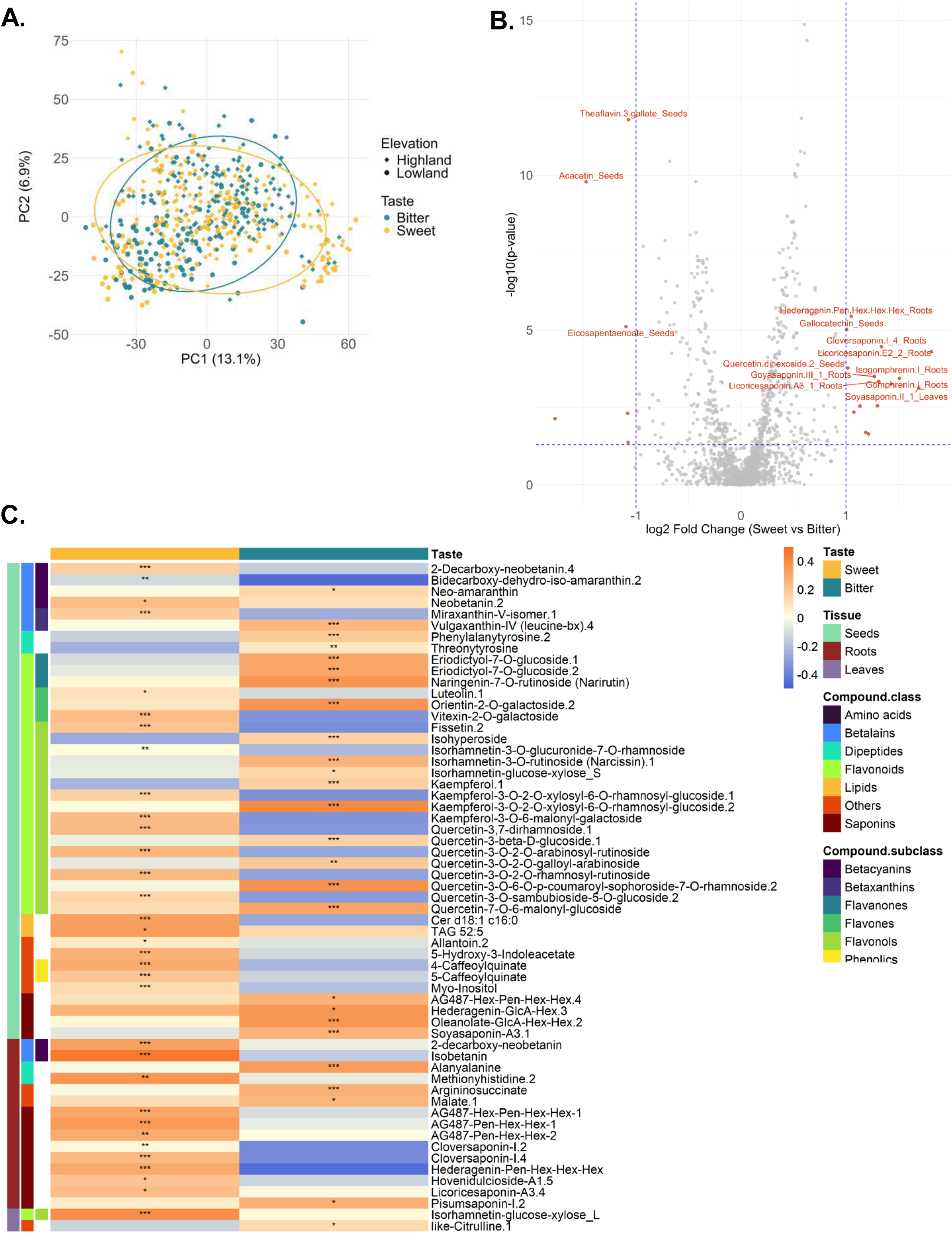
GWAS significant secondary and lipid compounds which correlate with sweet and bitter taste. (**A**) Principal component analysis of secondary metabolites of Fig. 2a grouped by elevation and taste. (**B**) Volcano plot of all annotated secondary metabolic features. Positive log_2_ fold change means the trait is higher in sweet accessions. Cut-off: *p* < 0.05, |log_2_ fold change| > 1. Names are displayed if *p* < 0.001, |log_2_ fold change| > 1. (**C**) Heatmap of 42 seed secondary and lipid features (6 betalains, 2 dipeptides, 23 flavonoids, 5 others, 4 saponins, 2 lipids), 15 root (2 betalains, 2 dipeptides, 2 others, 9 saponins) and 2 leaf secondary metabolites (1 flavonoid, 1 other) with a GWAS significant association. Significances were determined by either Student’s *t*-test or Wilcoxon test based on the data normal distribution (\**p* < 0.05, \*\**p* < 0.01, \*\*\**p* < 0.001).

**Figure S5.**
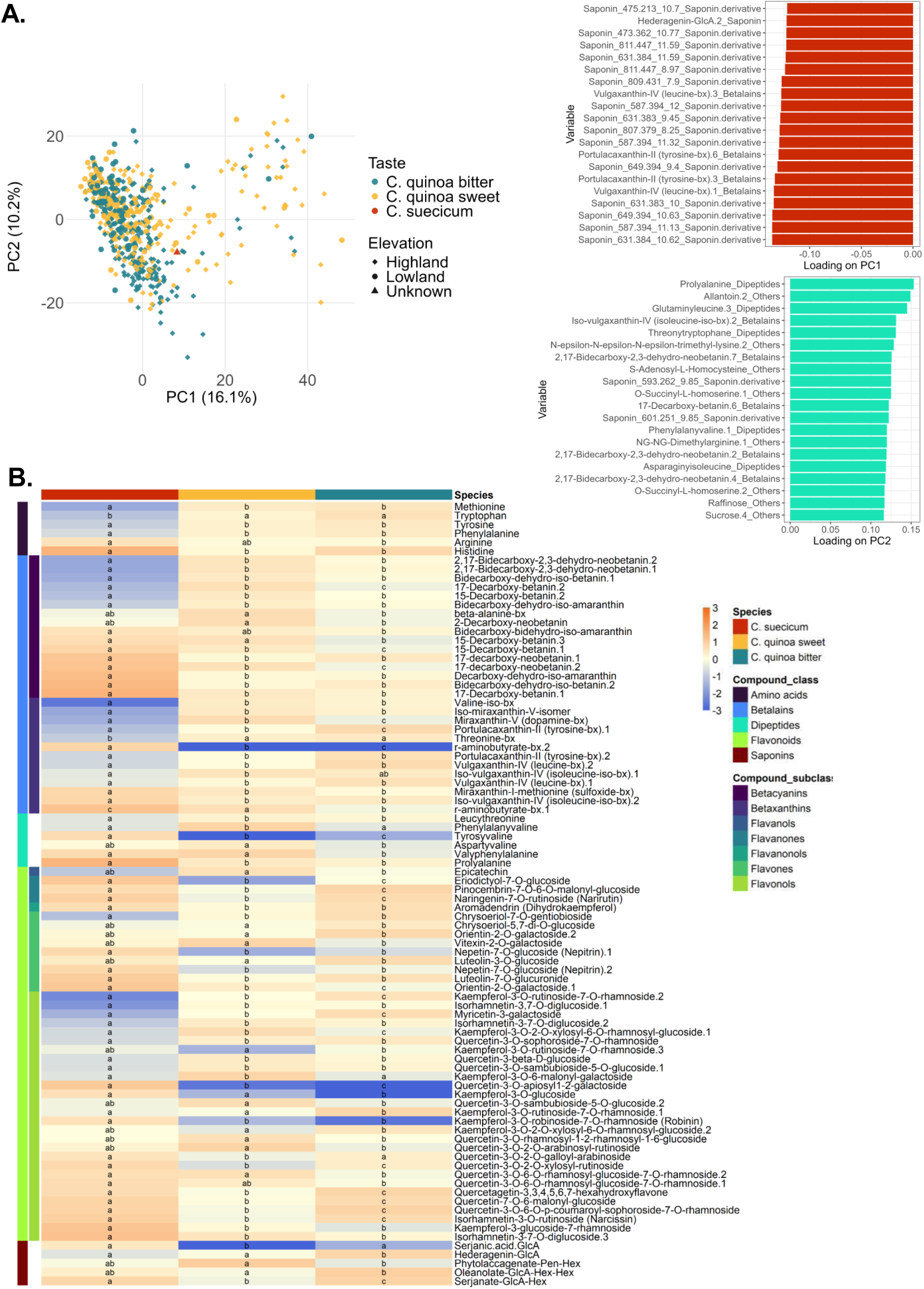
Wild variety *Chenopodium suecicum* (CHEN-100) compared with domesticated sweet and bitter *C. quinoa* varieties. (A) Principal component analysis of 828 common polar compounds (5 saponins, 71 saponin derivatives, 36 betalains, 10 dipeptides, 48 flavonoids, 31 other compounds, and 620 unknown features) and variable contribution plot of PC1 and PC2. (B) Heatmap of 88 compounds (6 amino acids, 29 betalains, 6 dipeptides, 42 flavonoids, 5 saponins) showing a significant difference across *C. suecicum*, and sweet and bitter *C. quinoa*. Letters indicate significances which were determined by either two-way-ANOVA or KRUSKAL-Wallis test based on the data

**Figure S6.**
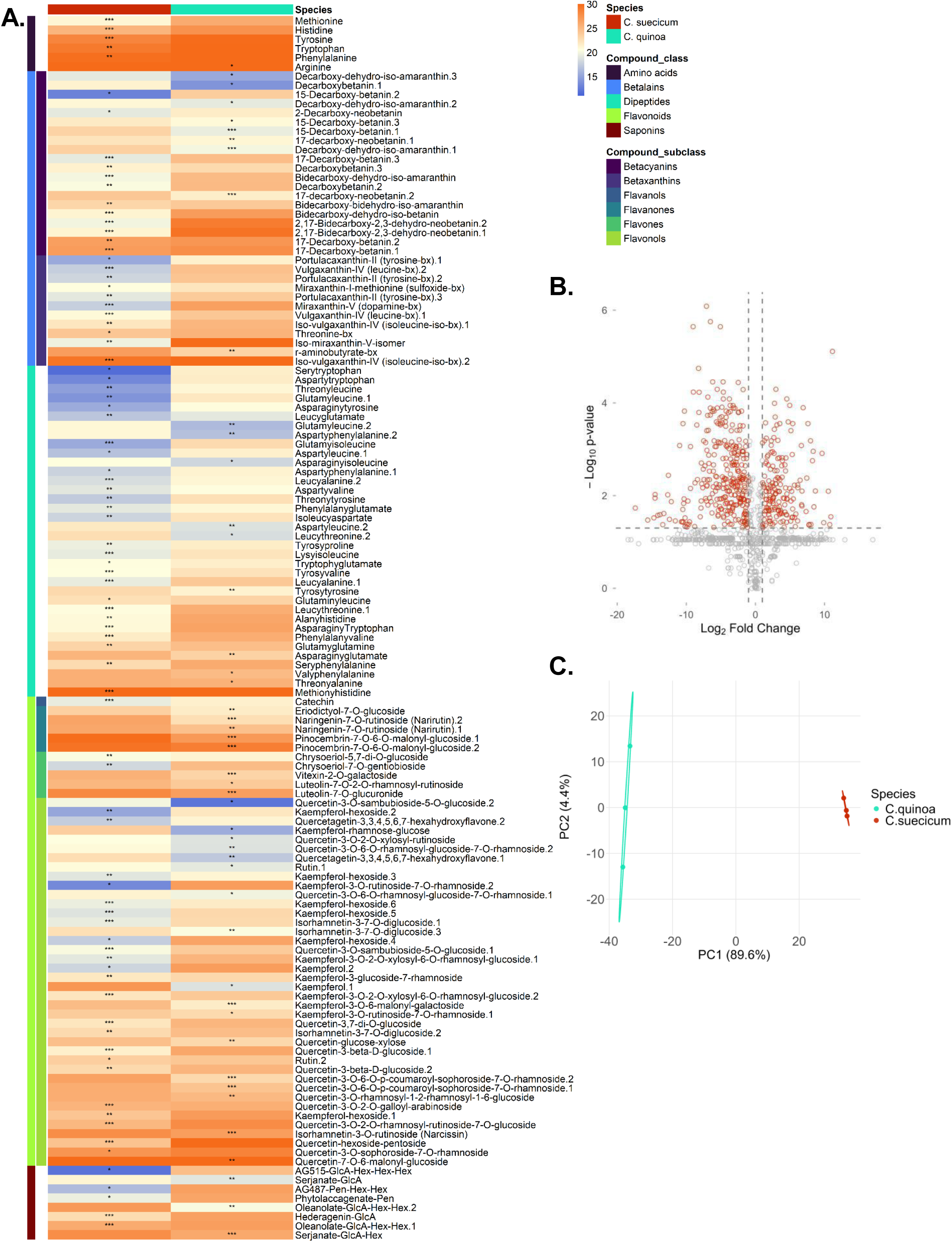
Wild variety *Chenopodium suecicum* (CHEN-100) compared with domesticated *C. quinoa* (CHEN-543). (**A**) Heatmap of 133 annotated secondary metabolites (6 amino acids, 32 betalains, 36 dipeptides, 51 flavonoids, 8 saponins) showing a significant difference across *C. suecicum*, and *C. quinoa*. Asterisks indicate significances which were determined by either Student’s *t*-test or Wilcoxon test based on the data normal distribution. (**B**) Volcano plot (cut-off: p-adj < 0.05, |log_2_FC| > 1) of 1465 compounds. (**C**) Principal component analysis of 106 common polar compounds (22 saponins, 8 betalains, 10 dipeptides, 27 flavonoids, and 39 other compounds).

**Figure S7.**
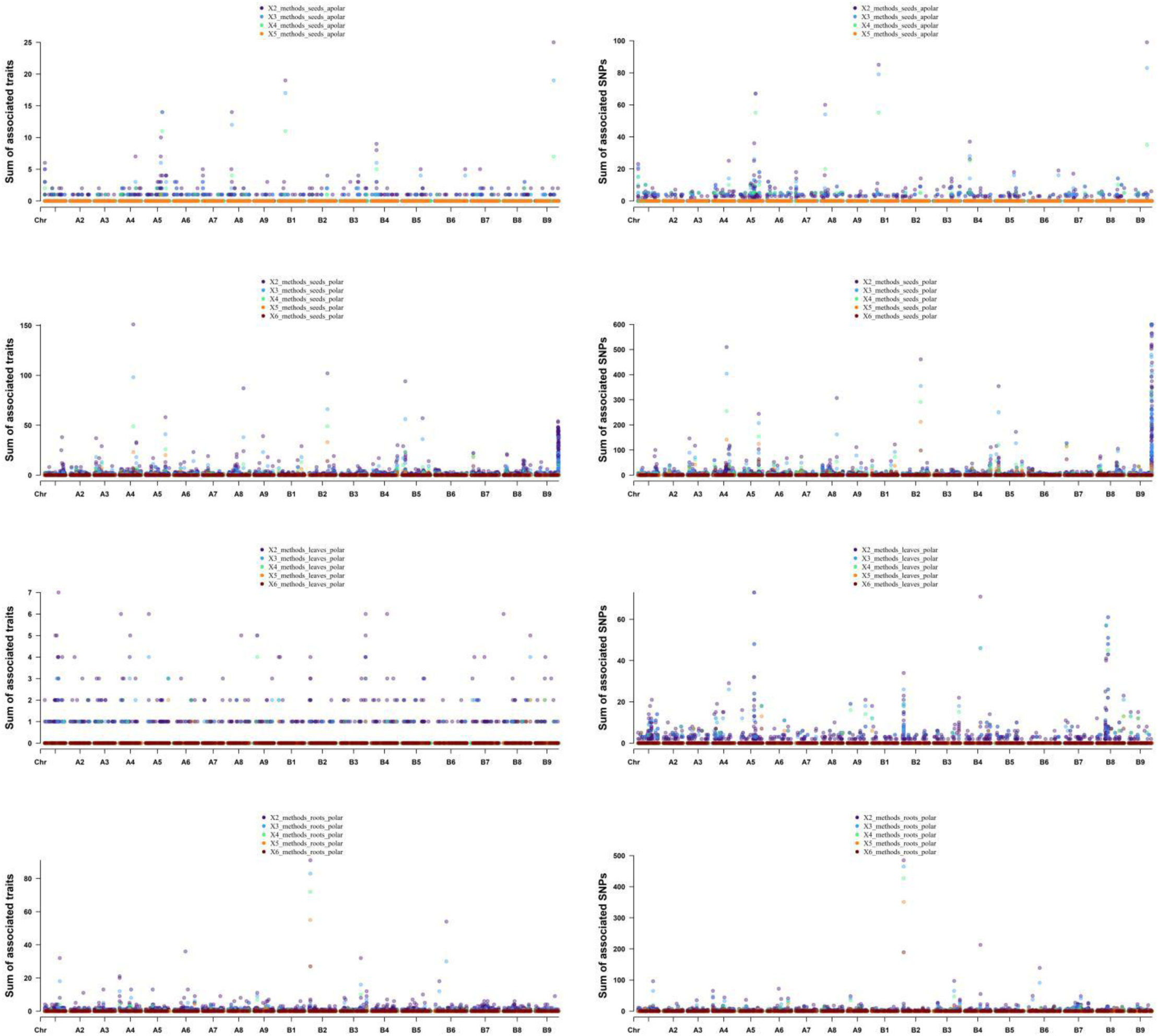
Overlaid Manhattan plots of genome-wide association studies (GWAS) of the total number of associated traits or SNPs for seed apolar and polar metabolites, and leaf and root polar secondary metabolites across six methods. Each dot represents a significant SNP detected by multiple GWAS models (BLINK, FarmCPU, EMMA-MLM, HE-MLM, MLMM, CMLM), colored by the number of methods in which the association was supported: two (purple), three (blue), four (green), five (orange), or six (red). For seed apolar metabolites, 412 associations were detected in ≥ two methods, 287 in ≥ three, 115 in ≥ four, one in ≥ five, and none in all six. For seed polar metabolites, 1,484 associations were detected in ≥ two methods, 1,164 in ≥ three, 624 in ≥ four, 309 in ≥ five, and 99 in all six. In leaves, 178 associations were detected in ≥ two methods, 96 in ≥ three, 44 in ≥ four, 20 in ≥ five, and five in all six. In roots, 381 associations were detected in ≥ two methods, 324 in ≥ three, 209 in ≥ four, 119 in ≥ five, and 46 in all six.

**Figure S8.**
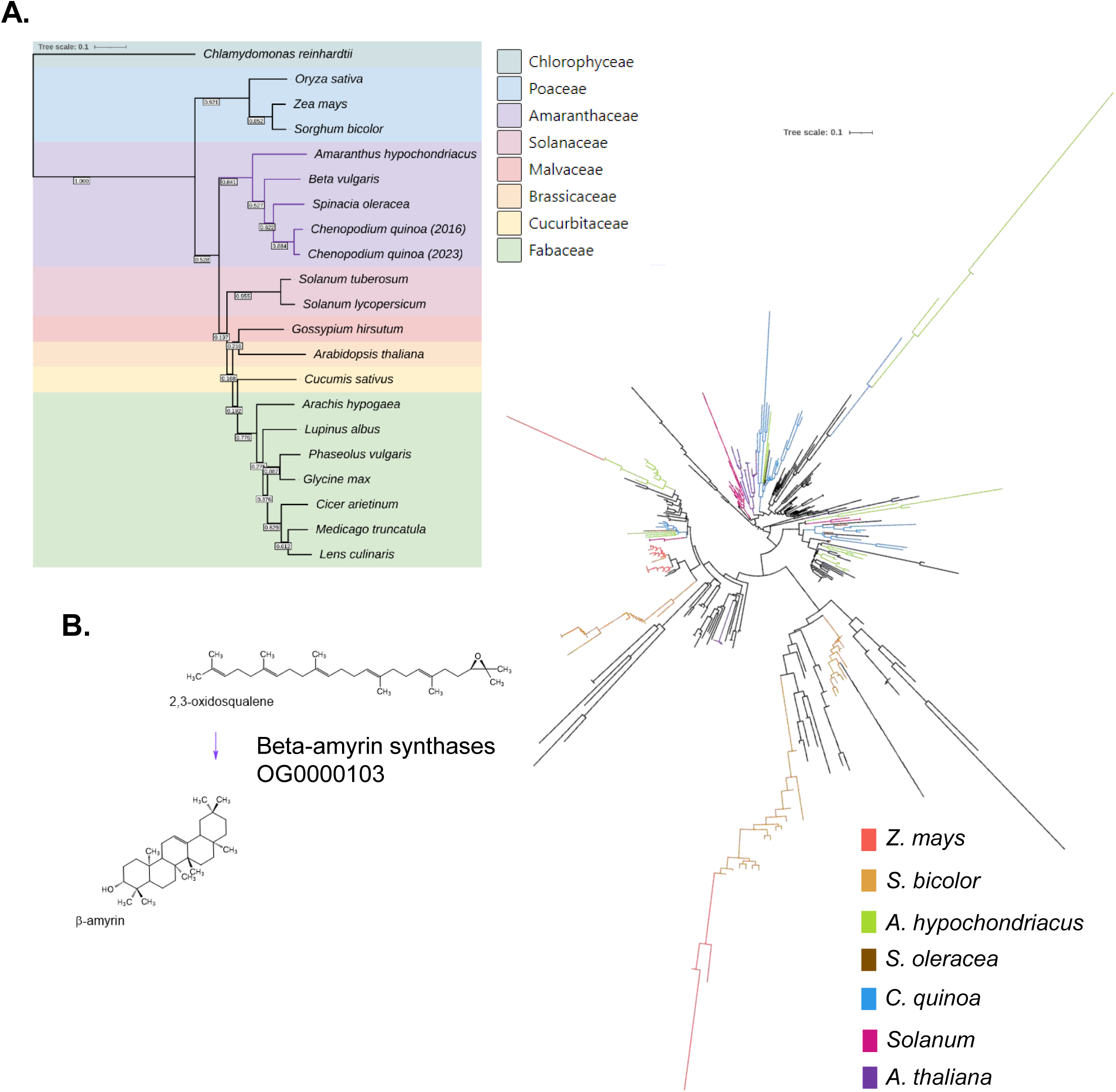
Identification of orthologues genes. OrthoFinder was used to discover orthologues across 20 crop species and eight families (**A**). In total 31,450 orthogroups were detected, OG0000103 containing beta-amyrin synthases serves as an example showing a high degree of duplication and speciation among species allowing conclusions about evolutionary lineages (**B**)

**Figure S9.**
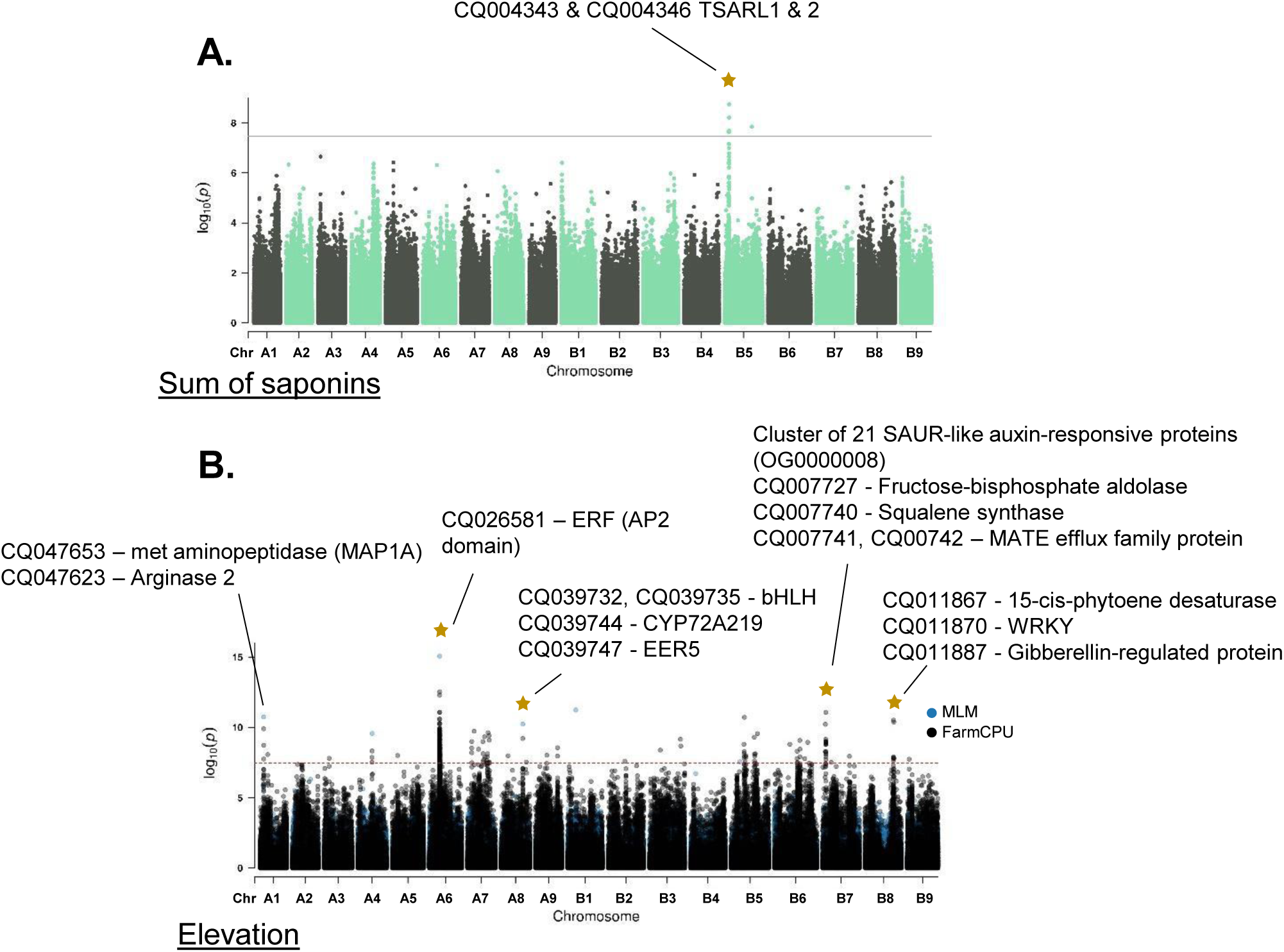
New genome-wide association panel maps to known loci. (**A**) The summed intensities of saponins measured by LC-MS mapped to the previously identified transcription factor triterpene saponin biosynthesis activating regulator 1 (TSARL1) and TSARL2 basic helix-loop-helix (bHLH) by Patirange et al., 2023. (**B**) GWAS results of mapping the elevation reveal four overlapping loci (asterisk) to previously identified loci of the principal component 1 (PC1), separating highland (Type I, blue) from lowland (Type II, red) accessions by Patirange et al., 2023. Here, five QTL containing 35 genes involved in hypoxia and plant development could be identified using farmCPU and MLM as GWAS models. ERF = ethylene response factor, AP2 = APETALA2, EER = enhanced ethylene response protein, SAUR = small auxin up-regulated RNA. The asterisk indicates similar associations.

**Figure S10.**
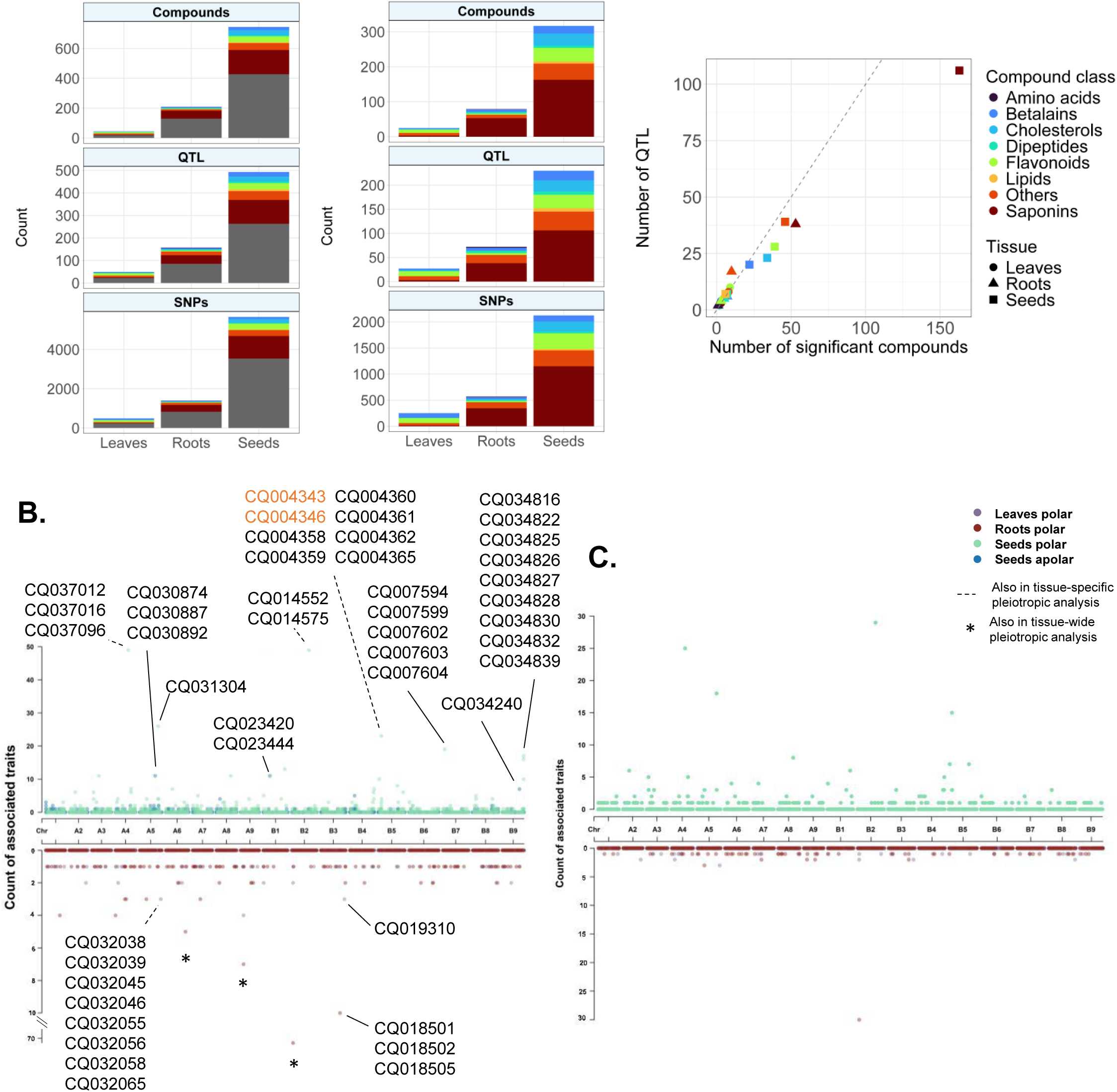
Sum of genomic associated annotated secondary metabolites. (**A**) Stacked bar plot with number of significant compounds, QTL in a 50 kb sliding window and SNPs using four genome-wide association study (GWAS) methods with and without untargeted compounds. Number of identified QTL in a 50 kb sliding window versus number of compounds with a GWAS association grouped by compound class and shaped by tissue without untargeted compounds. (**B**) Count of all metabolic features with significant genomic associations across tissues: 652 seed secondary metabolites, 119 seed lipid features, 44 leaf secondary metabolites, and 209 root secondary metabolites, detected in 573, 166, and 167 quinoa accessions, respectively. In total, 365, 82, 46, and 137 marker–trait associations (MTAs) were identified for seed secondary metabolites, seed lipids, and leaf and root secondary metabolites, respectively. Candidate genes are highlighted. (**C**) Count of annotated 317, 25 and 79 metabolic features of seed secondary metabolic and lipid features (upper), leaf and root secondary metabolic features (lower) of 573, 166 and 167 quinoa accessions, respectively, with a genomic association. In total 204, 26, 66 marker-trait association were detected for seed secondary metabolites and lipids, and leaf and root secondary metabolites.

**Figure S11.**
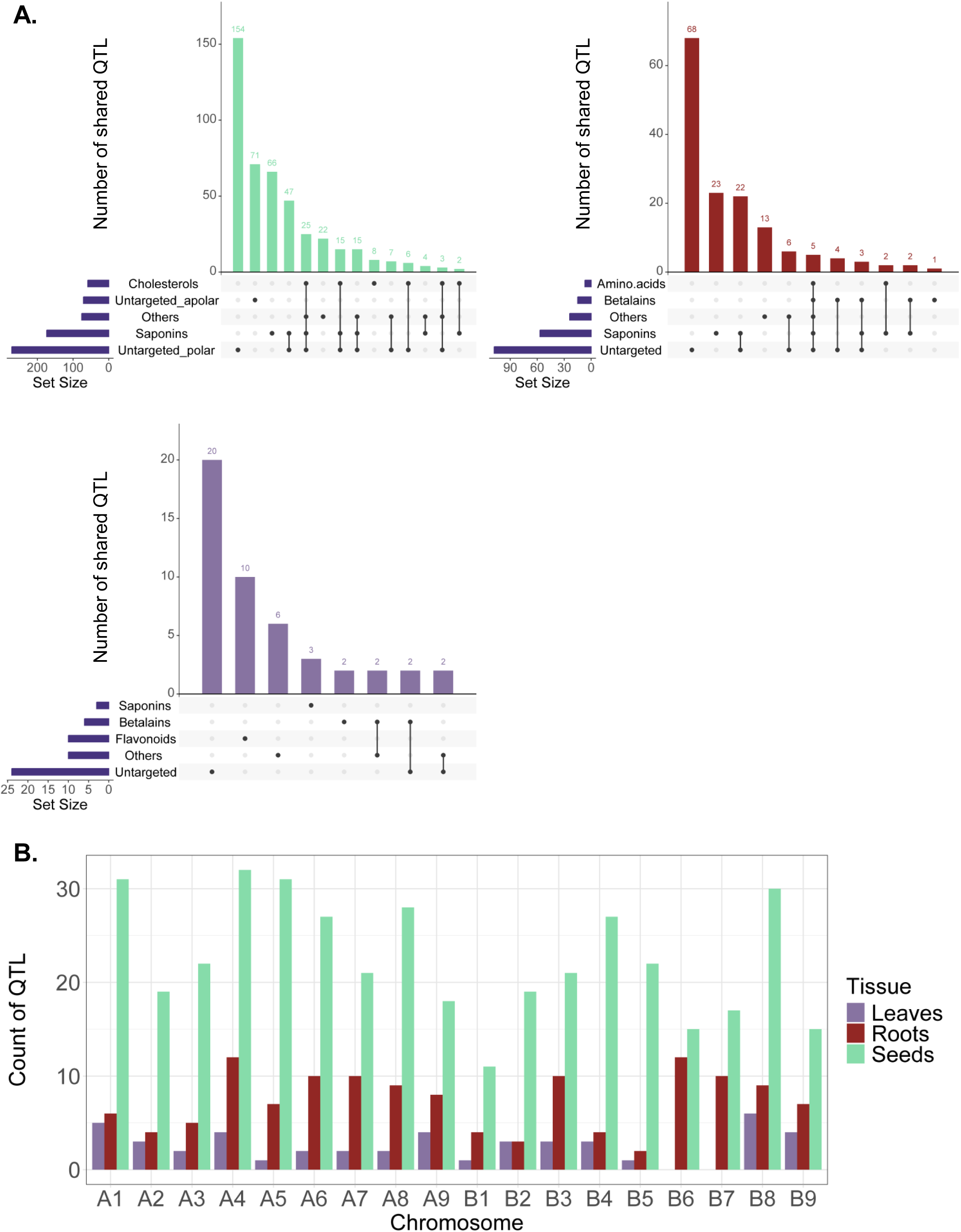
Total number of QTL in seeds, roots and leaves. (**A**) Intersections of QTL in a 50 kb sliding window in seeds (green), roots (red), and leaves (purple). In total 493, 157, and 49 QTL with 148, 46, and 6 shared QTL could be detected across compound classes for seeds, roots and leaves, respectively. (**B**) Count of QTL per chromosome of leaves, roots and seeds.

**Figure S12.**
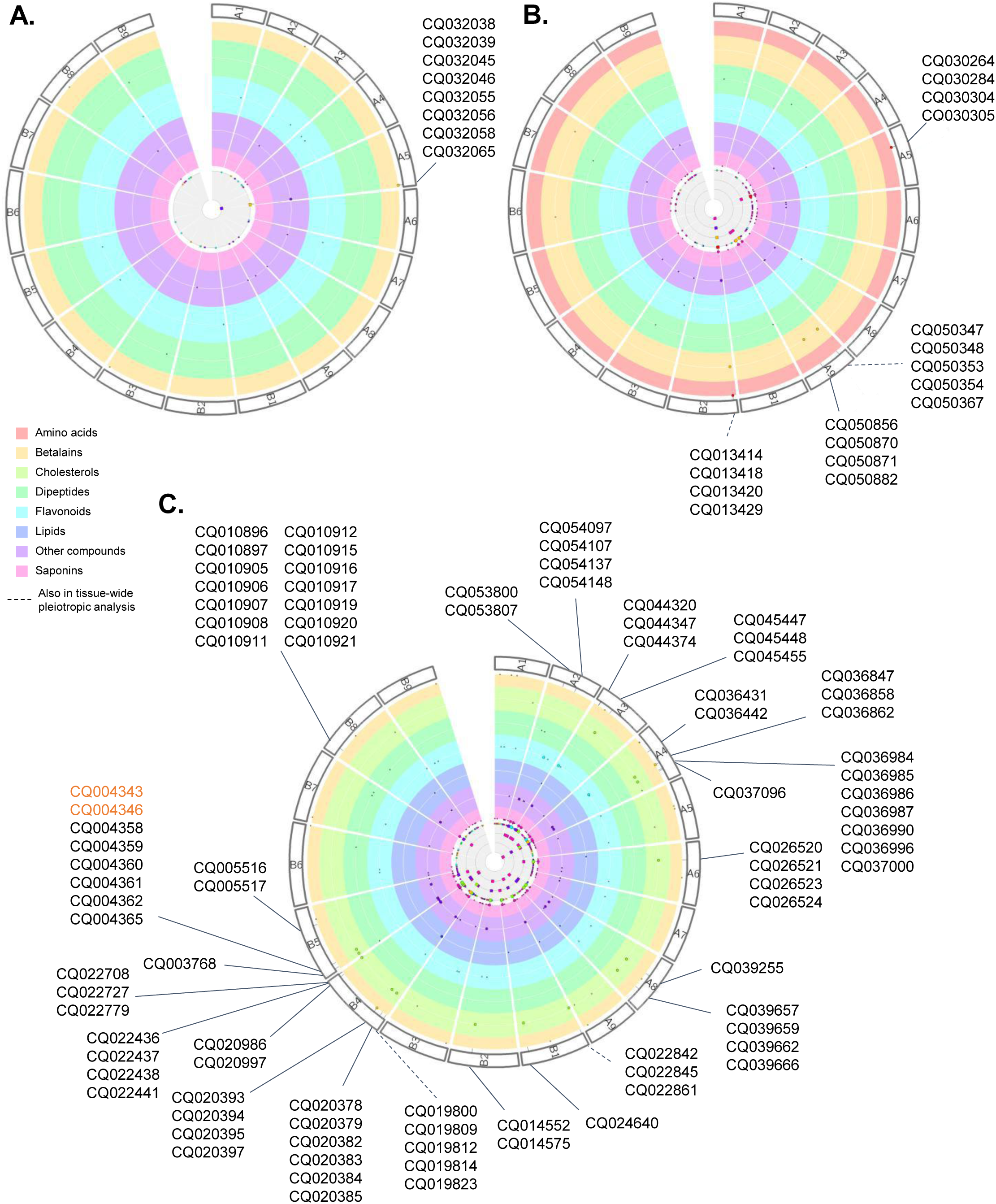
Pleiotropic analysis of annotated secondary metabolites and lipids in leaves, roots and seeds using ≥ four GWAS methods. Pleiotropic analysis of (A) leaves, (B) roots, and (C) seed secondary metabolites (A = amino acids, B = betalains, D = dipeptides, F = flavonoids, O = others, L = lipids, S = saponins) identifying 1, 4, and 24 common QTL, respectively.

**Figure S13.**
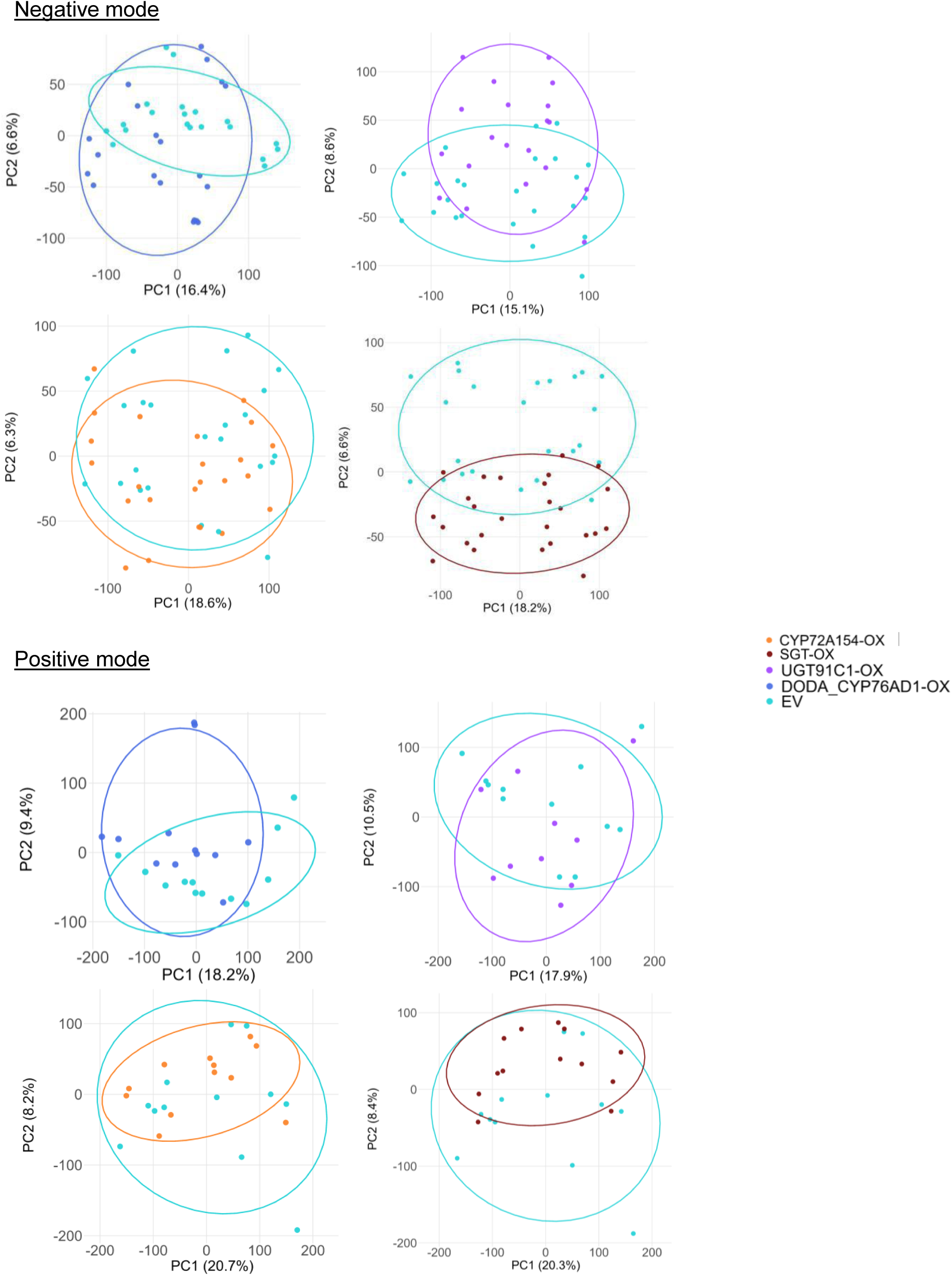
Transient overexpression in *Nicotiana benthamiana* leaves. Transient overexpression (OX) of CYP72A154, SGT, UGT91C1, DODA and CYP76AD1, and empty vector. Metabolic features were detected in negative and positive mode LC-MS.

**Figure S14.**
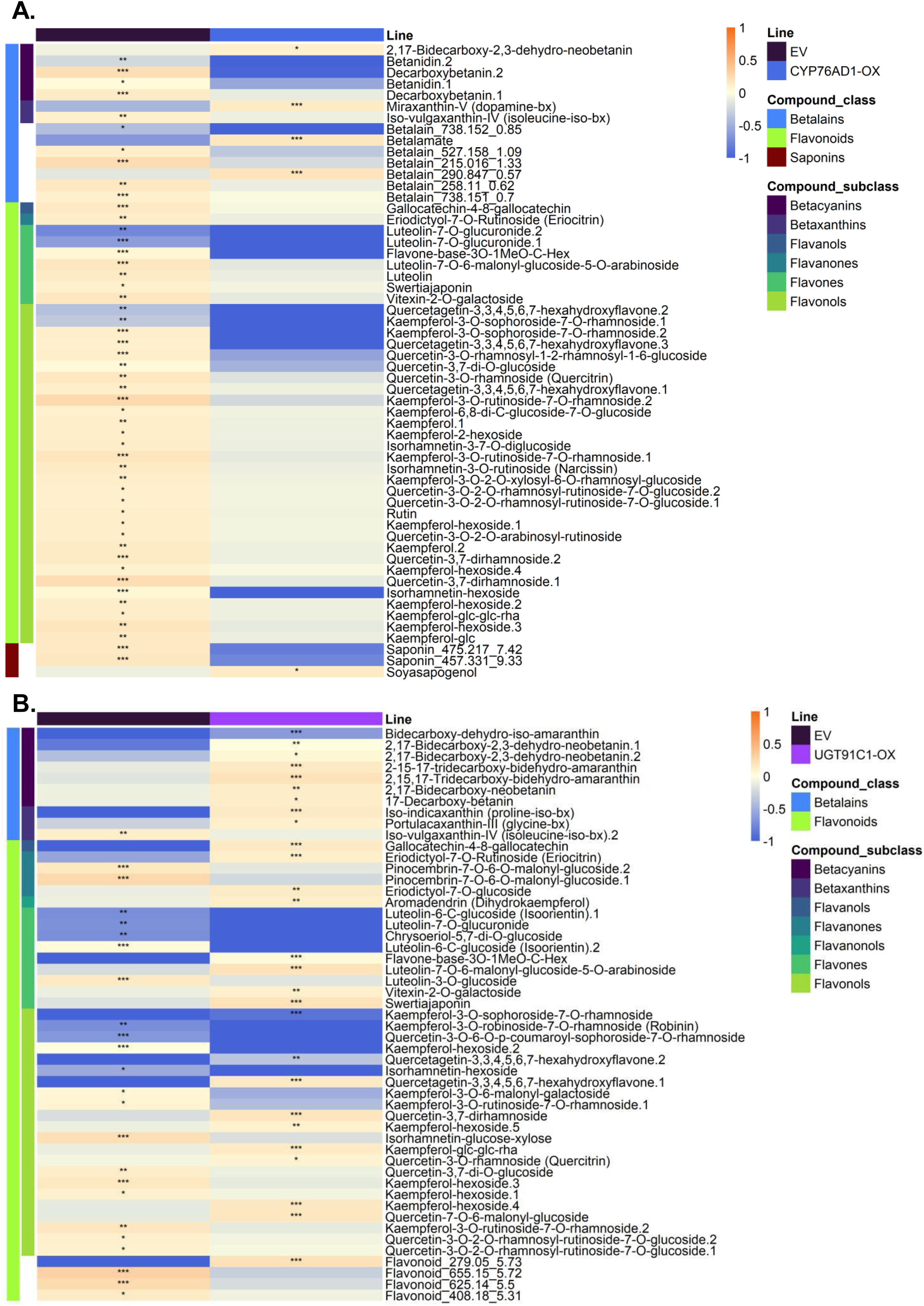
Transient overexpression in *Chenopodium quinoa* leaves. Heatmaps of transient overexpression (OX) of (**A**) CYP76AD1, and (**B**) UGT91C1 compared to empty vector.

**Figure S15.**
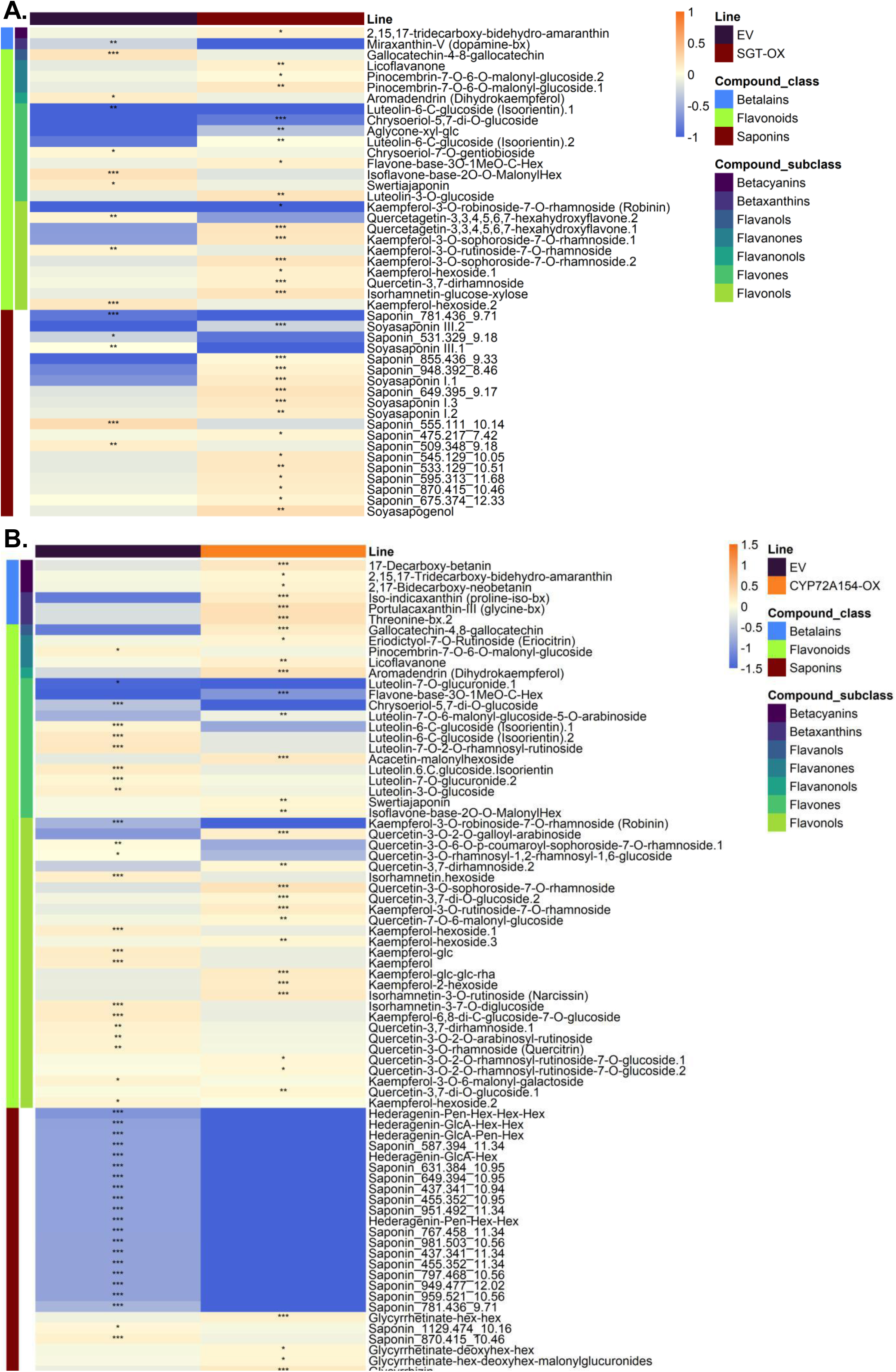
Transient overexpression in *Chenopodium quinoa* leaves. Heatmaps of transient overexpression (OX)

**Figure S16.**
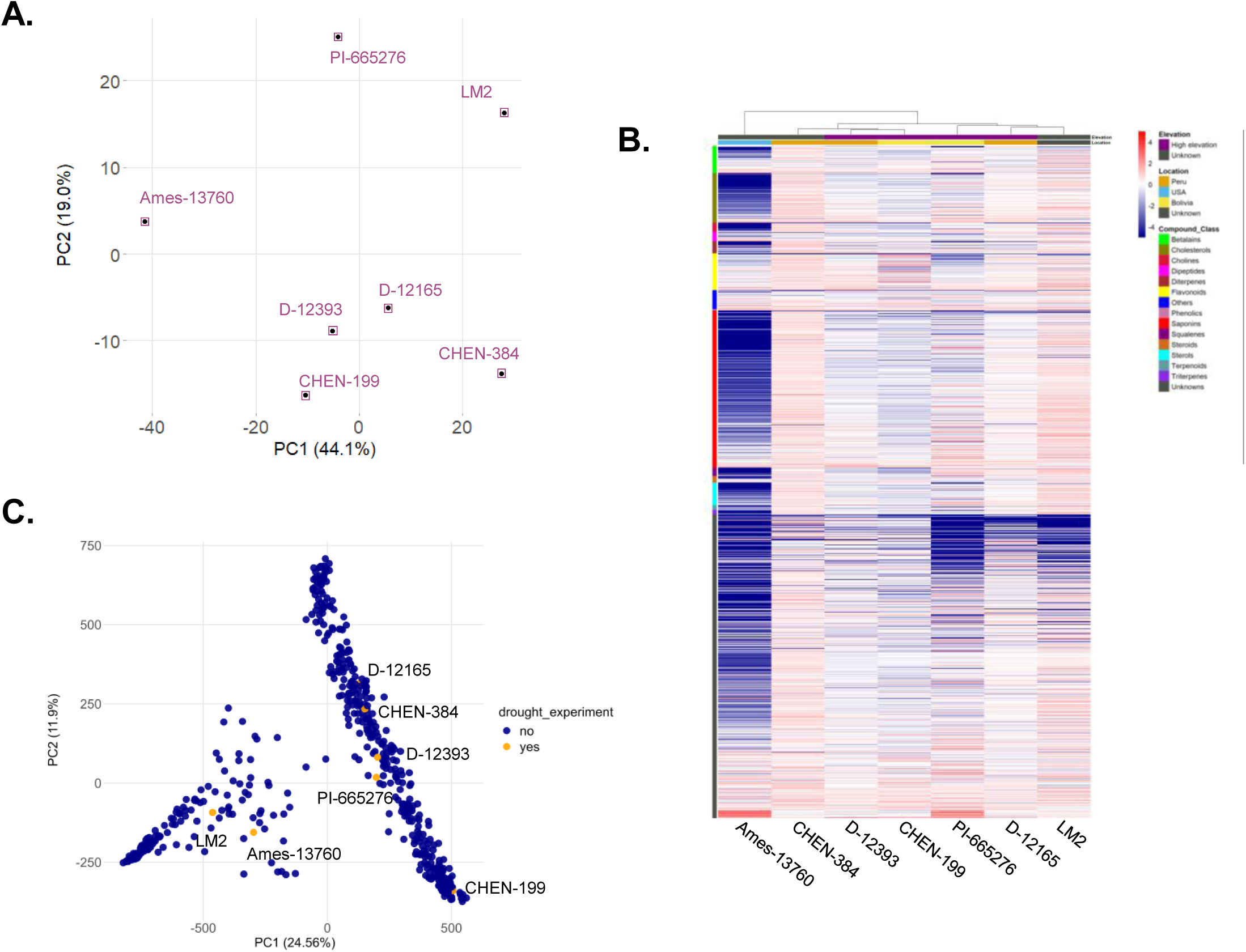
Seed metabolic diversity of 7 selected quinoa accessions for an integrative analysis. (**A**) Principal component analysis (PCA) of 7 accessions using 3029 metabolic features comprising 950 major peaks picked from the chromatograms and 2079 compounds putatively annotated (see material and methods). (**B**) Heatmap calculated using Euclidean distance measure and Ward clustering method. (**C**) PCA of 603 quinoa accessions. The accessions used for the drought experiment are coloured in orange.

**Figure S17.**
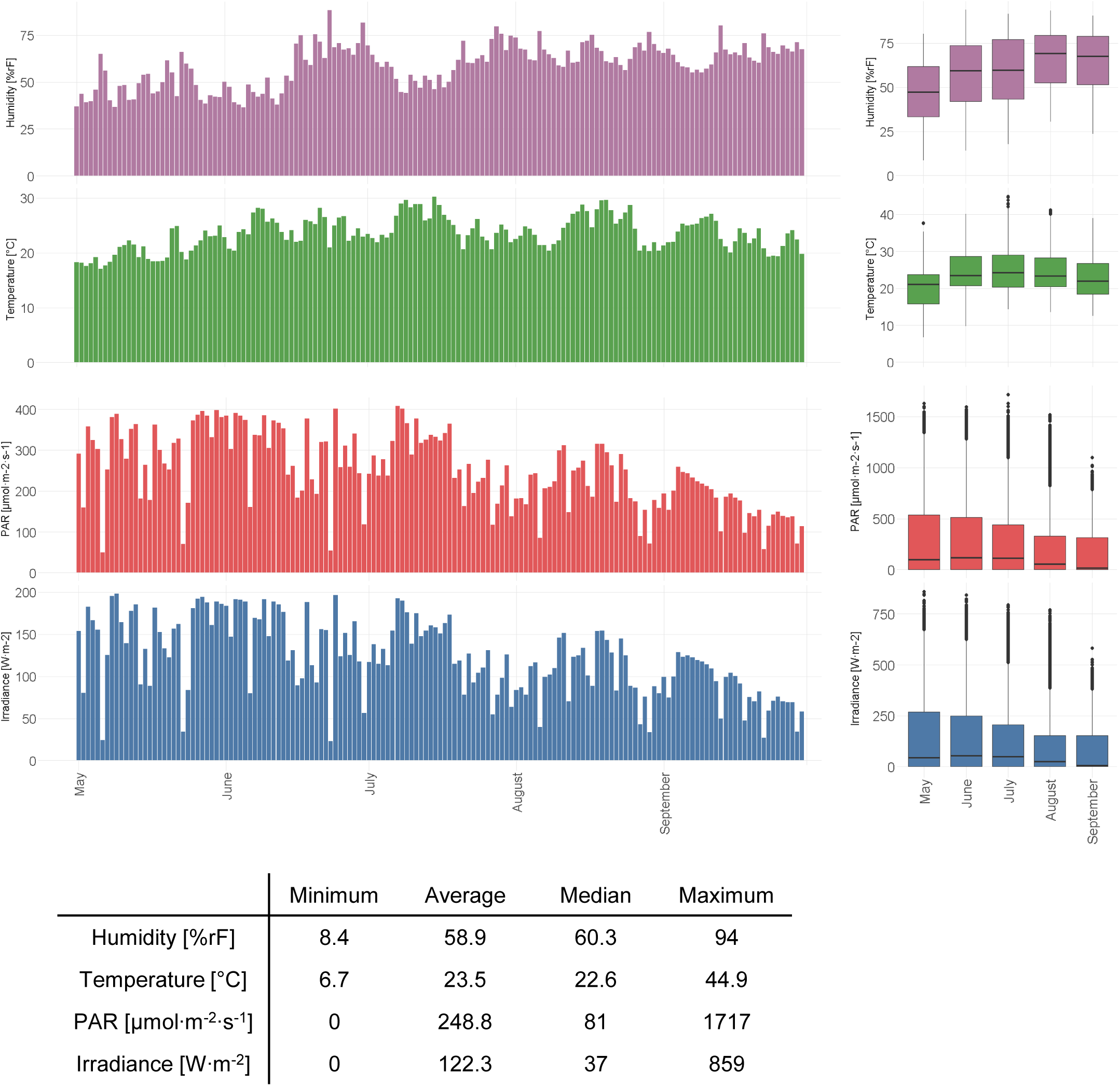
Climate conditions in the polytunnel. Barplot per day and boxplots per month of humidity [%rF], temperature [°C], photosynthetically active radiation (PAR) [µmol·m^-2^·s^-1^], and irradiance[W·m^-2^] from May to September 2023. The table shows the minimum, average, median and maximum values per condition.

**Figure S18.**
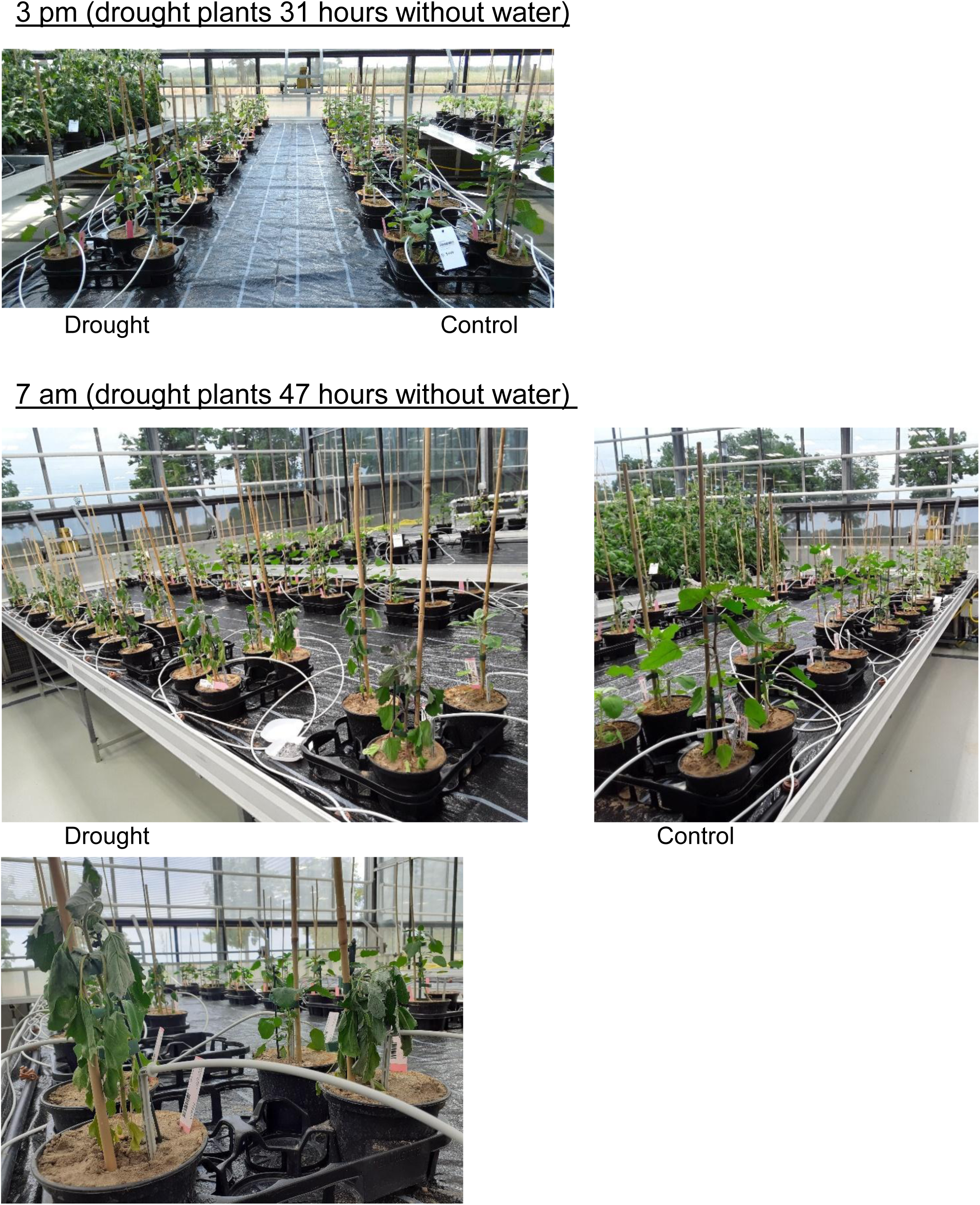
Quinoa drought experiment in the greenhouse. *Chenopodium quinoa* accessions under drought stress were irrigated every second day while control plants every day. The water supply was adjusted according to the turgescence of the control plants.

**Figure S19.**
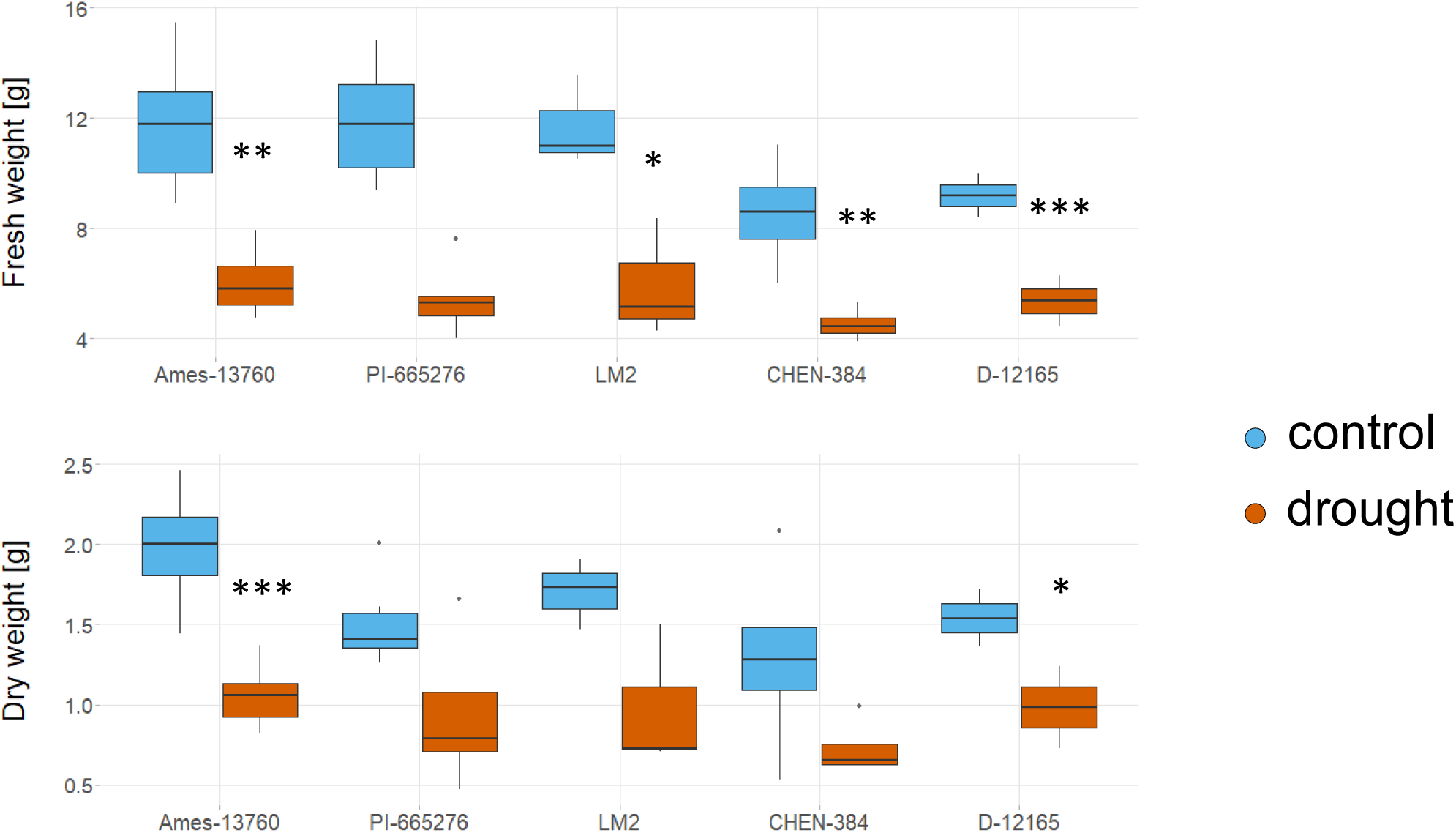
Quinoa drought experiment in the greenhouse. Fresh and dry weight of the *Chenopodium quinoa* accessions under control and drought stress conditions. Plants under drought were irrigated every second day while control plants every day. The water supply was adjusted according to the turgescence of the control plants.

**Figure S20.**
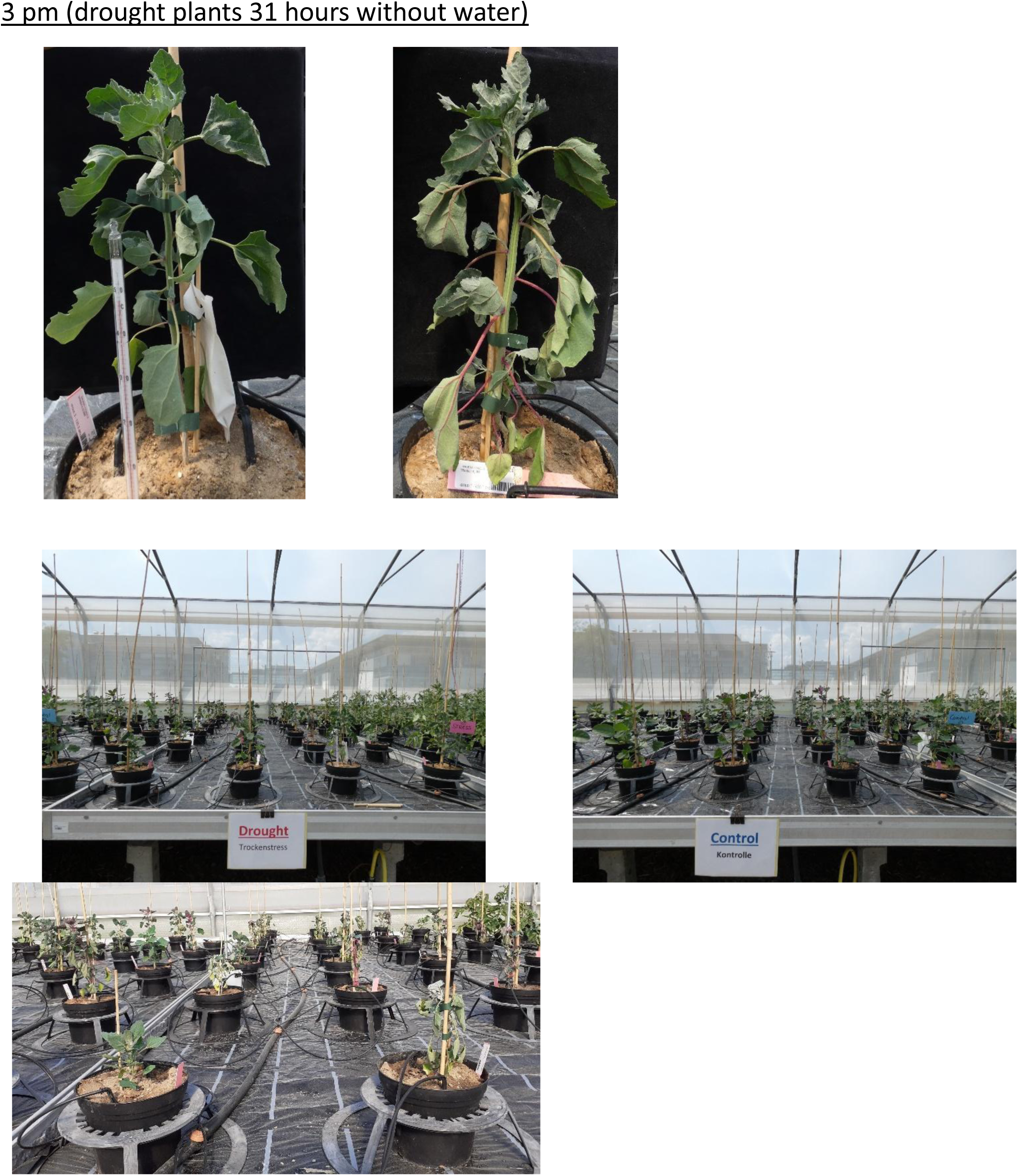
Quinoa drought experiment in the polytunnel. *Chenopodium quinoa* accessions under drought stress were irrigated every second day while control plants every day. The water supply was adjusted according to the turgescence of the control plants. Soil temperature reached 48 °C on 08/06/2023.

**Figure S21.**
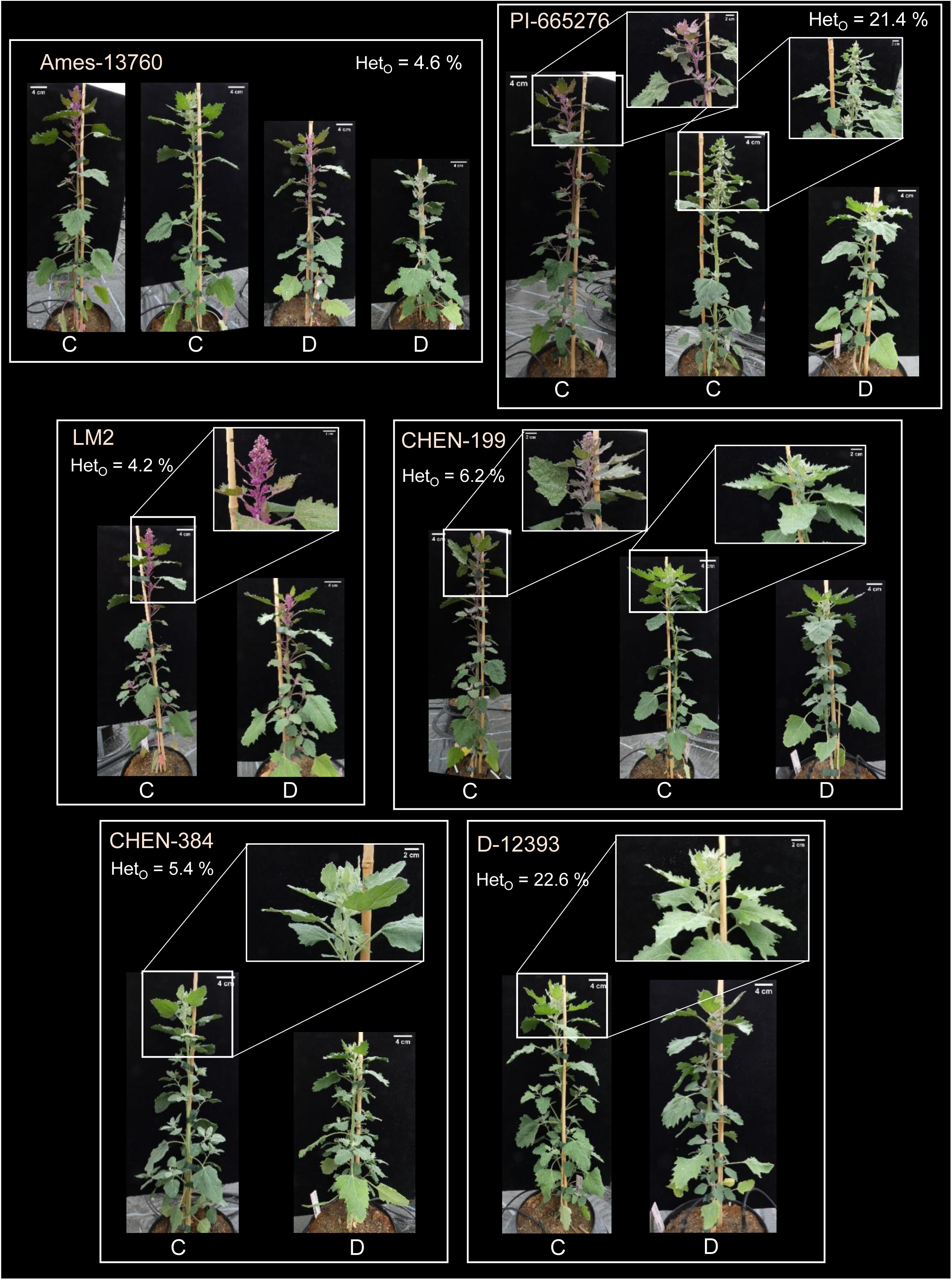
Phenotype of the *Chenopodium quinoa* accessions grown in the polytunnel. Ames-13760, PI-665276, LM2, CHEN-199, CHEN-384, D-12393 under control (C) and drought (D) conditions with calculated observed heterozygosity (Het_O_) in percent. Scale bar = 4 cm (whole plant), 2 cm (zoom in). Colors of names indicate clusters in PCA (Figure 8).

**Figure S22.**
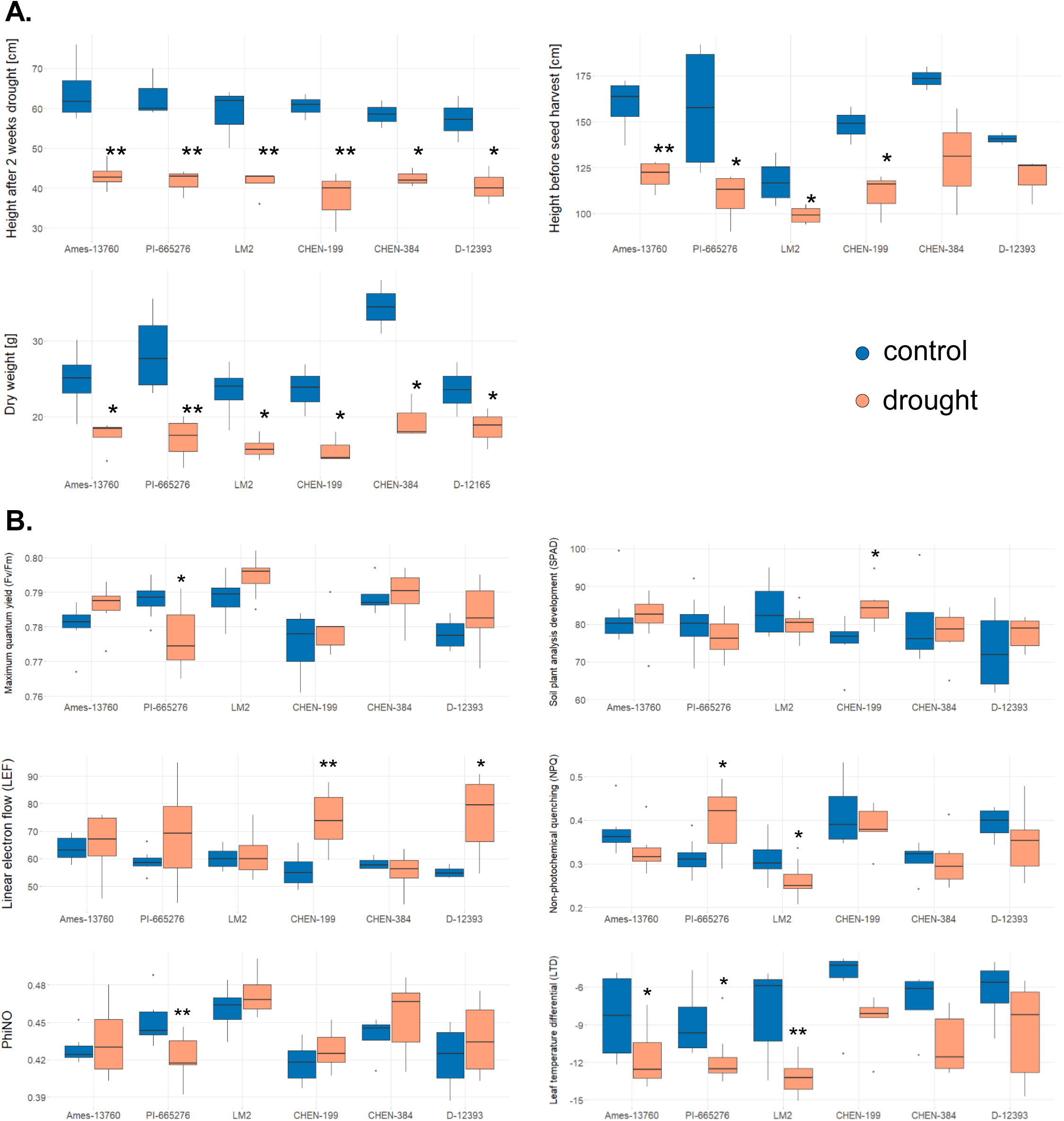
Phenotype of Chenopodium quinoa accessions grown in the polytunnel. (**A**) Height and dry weight of the *C. quinoa* accessions Ames-13760, PI-665276, LM2, CHEN-199, CHEN-384, D-12393 under control (blue) and drought (orange) conditions. (**B**) Photosynthetic activity of the *C. quinoa* accessions. Maximum quantum yield as Fv/Fm, soil plant analysis development (SPAD) value as an indicator of plant nitrogen status and relative chlorophyll, linear electron flow (LEF), nonphotochemical chlorophyll fluorescence quenching (NPQ) as a measure of dissipated heat or fluorescence, PhiNO-ratio as a measure of excited electrons that are lost in non-regulated processes and cause photodamage, leaf temperature differential (LTD), the temperature difference between the leaf and its environment. ***p < 0.001, **p < 0.001, *p < 0.01; Student’s t-test.

**Figure S23.**
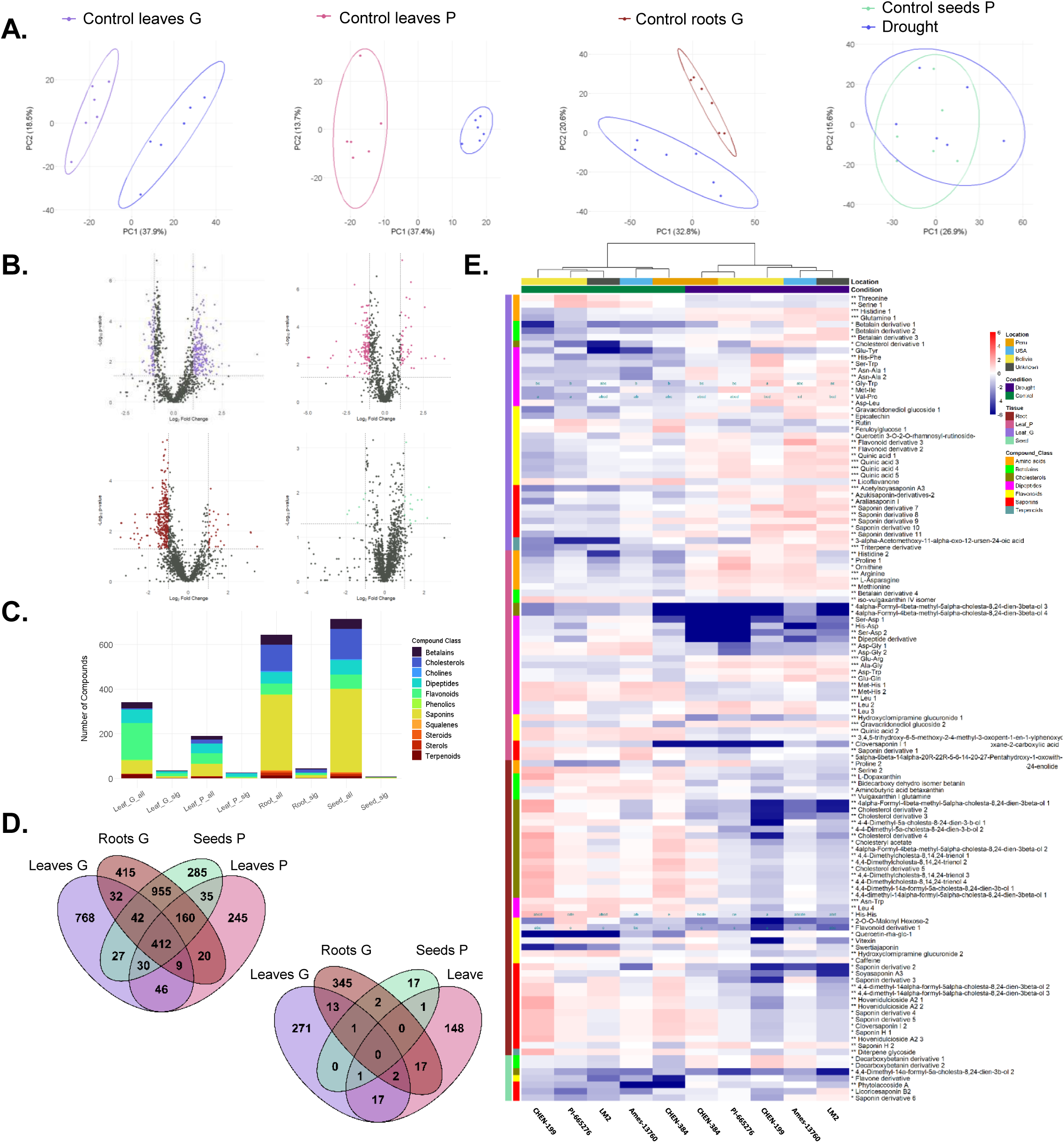
Secondary metabolic diversity of six quinoa accessions under drought conditions. Roots and leaves of the accessions CHEN-199, D-12165, Ames-13760, CHEN-384, PI-665276 and LM2 were harvested in the greenhouse (G) while leaves and seeds of the accessions CHEN-199, Ames-13760, CHEN-384, D-12393, PI-665276 and LM2 were harvested in the polytunnel (P). (**A**) Principal component analysis of 1365, 956, 2045 and 1945 metabolic features of leaves G and P, roots G and seeds P, respectively. Ellipse shows the 80 % confidence interval. (**B**) Volcano plot of leaves grown in the greenhouse (purple) and polytunnel (pink), roots (red) and seeds (green). Significance was calculated either by Student’s t-test or by Wilcoxon test depending on their normal distribution. Cut off: p < 0.05 and |log_2_ fold change| > 1. (**C**) Counts of compound classes per tissue without unknown compounds with either all identified compounds (“_all”, 549 leaf G, 370 leaf P, 858 root and 940 seed compounds) or only compounds with p < 0.05 (“_sig”, 117 leaf G, 138 leaf P, 105 root and 61 seed compounds). (**D**) Venn-diagram of all and only significant (p-value < 0.05, log2 fold change > |1|) common compounds in leaves G and P, roots (G) and seeds (P). (**E**) Heatmap of significant (p-value < 0.05, log2 fold change > |1|; total: 112 compounds; Leaves G: 35 compounds – 3 betalains, 1 cholesterols, 9 dipeptides, 12 flavonoids, 8 saponins, 2 terpenoids; Leaves P: 26 compounds – 2 betalains, 2 cholesterols, 15 dipeptides, 4 flavonoids, 3 saponins; Roots: 44 compounds – 4 betalains, 15 cholesterols, 3 dipeptides, 8 flavonoids, 13 saponins, 1 terpenoid; Seeds: 7 compounds – 2 betalains, 1 cholesterols, 1 flavonoids, 3 saponins) compounds of the five common accessions grown in the greenhouse and the polytunnel identified using either Student’s t-test or Wilcoxon test based on their normal distribution.

**Figure S24.**
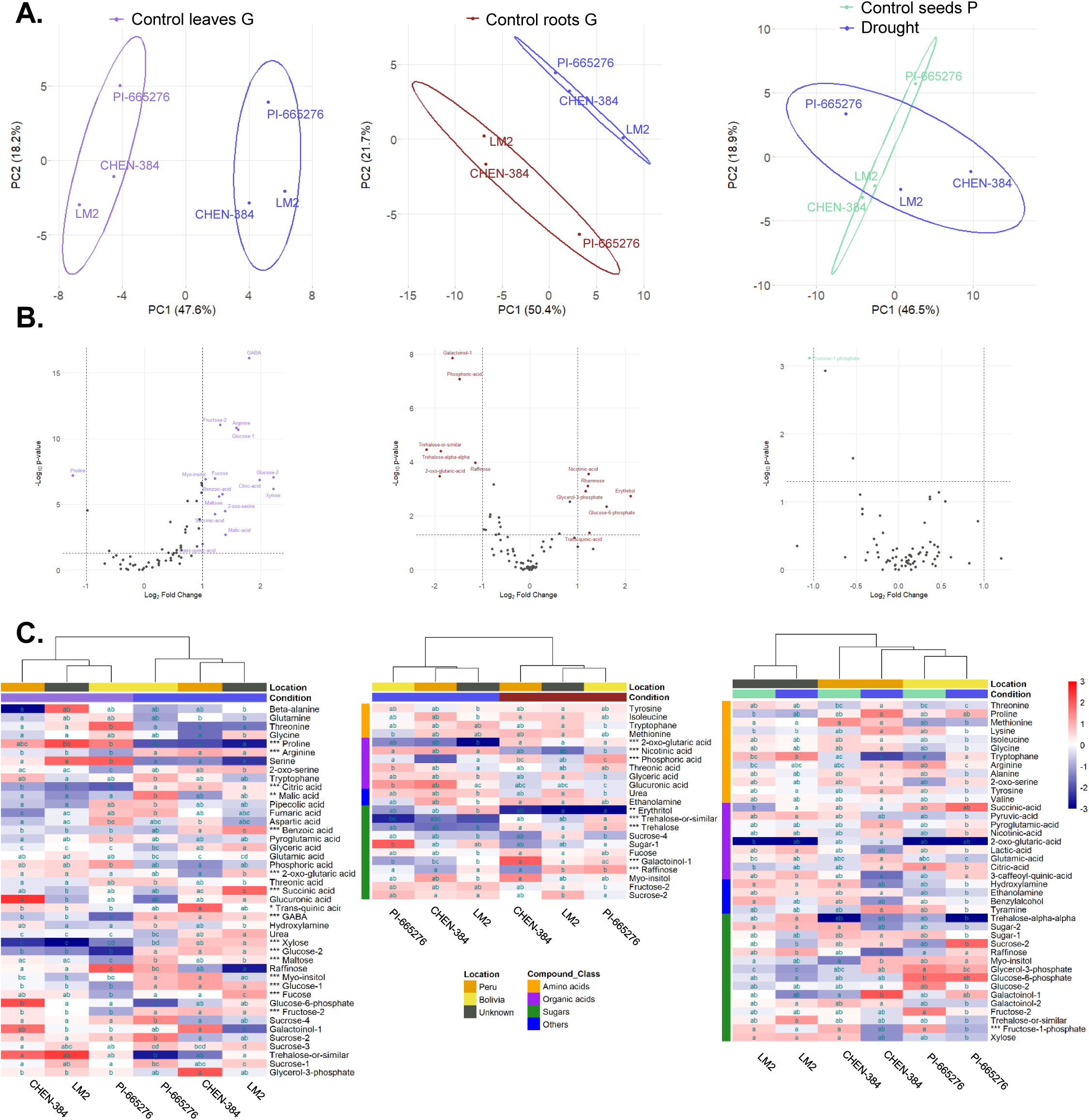
Primary metabolites of three quinoa accessions under drought conditions. Leaves (purple), roots (red) from the greenhouse and seeds (green) from the polytunnel of the accessions CHEN-384, PI-665276 and LM2 were utilized for GC-MS analysis. (**A**) Principal component analysis with the 80 % confidence interval and (**B**) volcano plot of control versus drought of all accessions calculated either by Student’s t-test or by Wilcoxon test depending on their normal distribution with cut off criterion of p < 0.05 and |log_2_ fold change| > 1 of 69 (18 amino acids, 20 organic acids, 24 sugars, 7 others), 67 (17 amino acids, 20 organic acids, 24 sugars, 7 others) and 70 (18 amino acids, 20 organic acids, 25 sugars, 7 others) primary metabolic features of leaves, roots and seeds, respectively. (**C**) Heatmap of significant compounds of the five common accessions grown in the greenhouse and the polytunnel (43 compounds of leaves, 23 compounds of roots, 40 compounds of seeds). Letters indicate significances calculated either by ANOVA and post hoc Tukey HSD test or Kruskal-Wallis test and post hoc Dunn’s test with p-value < 0.05. Asterisks next to the name indicate significances from (B; p-value < 0.001***, < 0.01**, < 0.05*).

**Figure S25.**
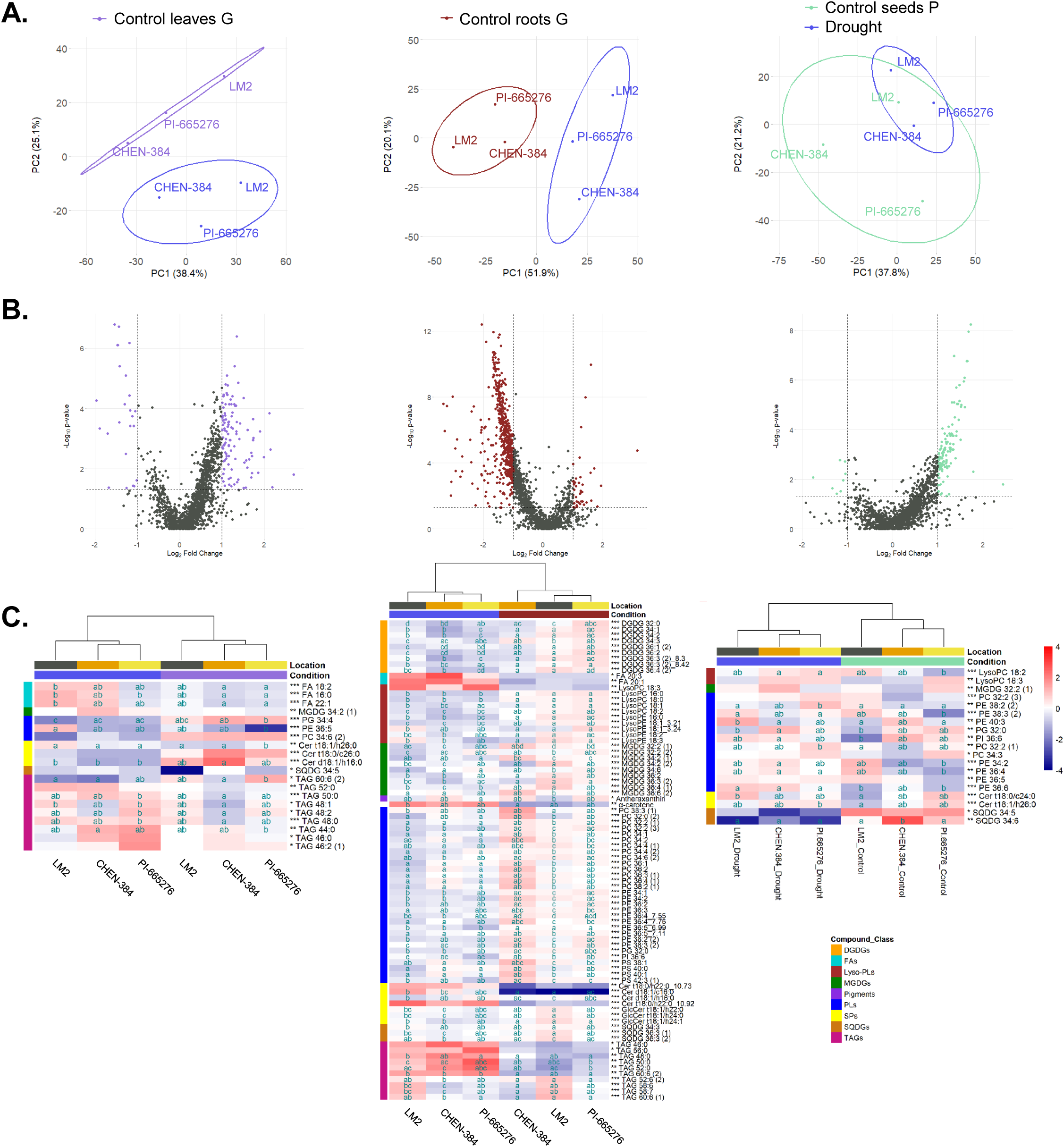
Lipids of three quinoa accessions under drought conditions. Leaves (purple), roots (red) from the greenhouse and seeds (green) from the polytunnel of the accessions CHEN-384, PI-665276 and LM2 were utilized for UPLC-MS analysis. (**A**) Principal component analysis with the 80 % confidence interval and (**B**) volcano plot of control versus drought of all accessions calculated either by Student’s t-test or by Wilcoxon test depending on their normal distribution with cut off criterion of p < 0.05 and |log_2_ fold change| > 1 of 1707 (7 pigments, 20 fatty acids, 224 annotated lipids, 627 less confident annotated lipids and 829 unknown), 1736 (7 pigments, 20 fatty acids, 224 annotated lipids, 637 less confident annotated lipids and 843 unknown) and 1605 (7 pigments, 20 fatty acids, 226 annotated lipids, 629 less confident annotated lipids and 768 unknown) lipid features of leaves, roots and seeds, respectively. (**C**) Heatmap of significant compounds (20 compounds of leaves: 4 fatty acids (FAs), 1 monogalactosyldiacylglycerol (MGDG), 3 phospholipids (PLs), 3 sphingolipids (SPs), 1 sulfoquinovosyldiacylglycerol (SQDG), 9 triacylglycerols (TAGs); 82 compounds of roots: 9 digalactosyldiacylglycerols (DGDGs), 2 FAs, 10 *lyso*-PLs, 9 MGDGs, 2 pigments, 30 PLs, 7 SPs, 3 SQDGs, 10 TAGs; 19 compounds of seeds: 2 *lyso*-PLs, 1 MGDG, 12 PLs, 2 SPs, 2 SQDGs). Letters indicate significances calculated either by ANOVA and post hoc Tukey HSD test or Kruskal-Wallis test and post hoc Dunn’s test with p-value < 0.05. Asterisks next to the name indicate significances from (B; p-value < 0.001***, < 0.01**, < 0.05*).

**Figure S26.**
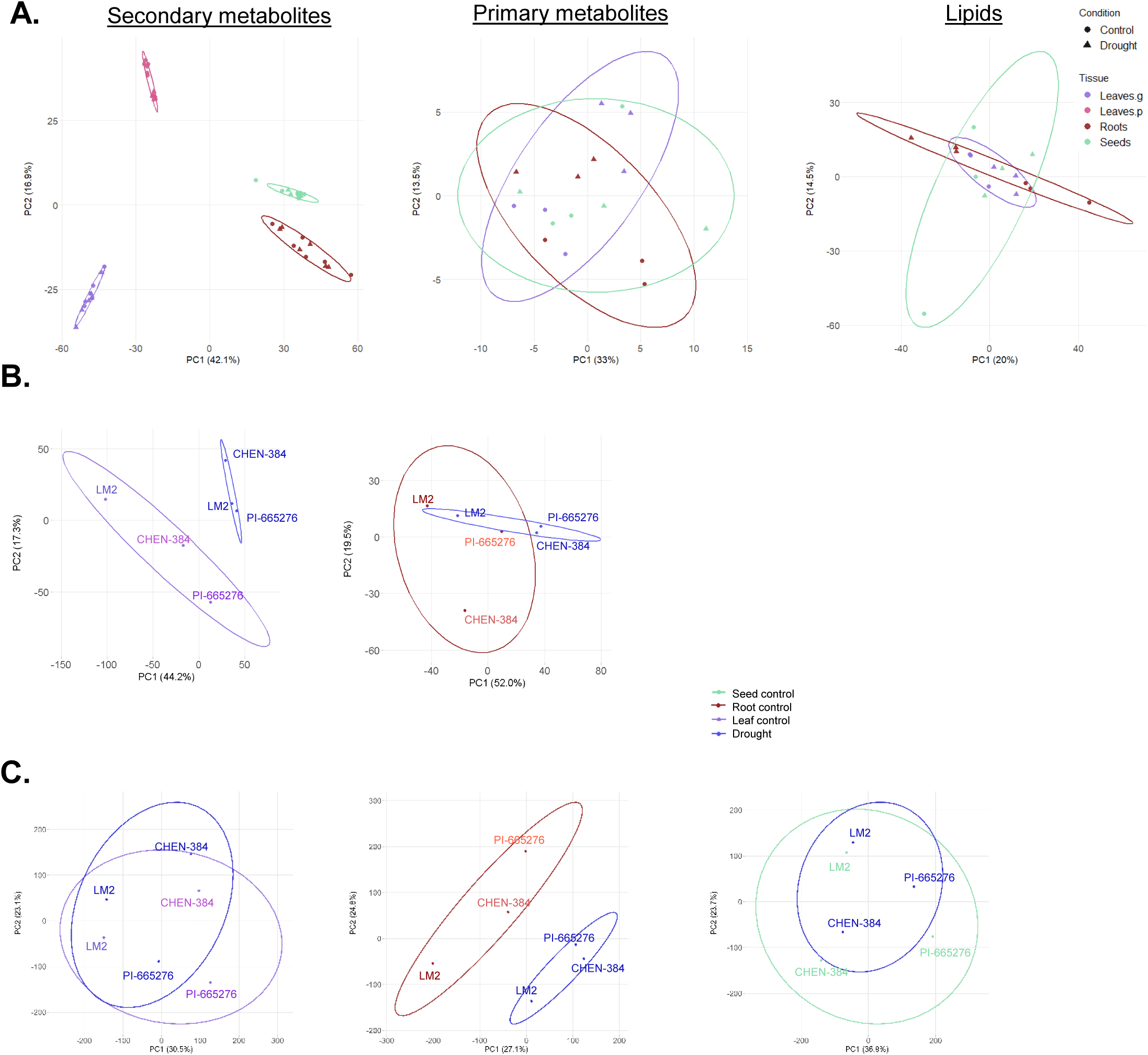
Metabolic, proteomic and transcriptomic diversity of quinoa accessions under drought conditions. (**A**) Metabolic diversity of control versus drought CHEN-384, LM2, PI-665276, CHEN-199, Ames-13760 and either D-12393 or D-12165 in leaves, seeds and roots. Principal component analysis of secondary, primary and lipid features from leaves of the greenhouse (g) or polytunnel (p), roots and seeds. (**B**) Principal component analysis of 6574 proteins of quinoa leaves (purple) and 2001 proteins of quinoa roots (red) of the accessions CHEN-384, PI-665276 and LM2. (**C**) Principal component analysis of 47732 genes of quinoa leaves (purple), 50332 genes of quinoa root (red) and 46557 genes of quinoa seeds (green). Ellipse shows the 80 % confidence interval.

**Figure S27.**
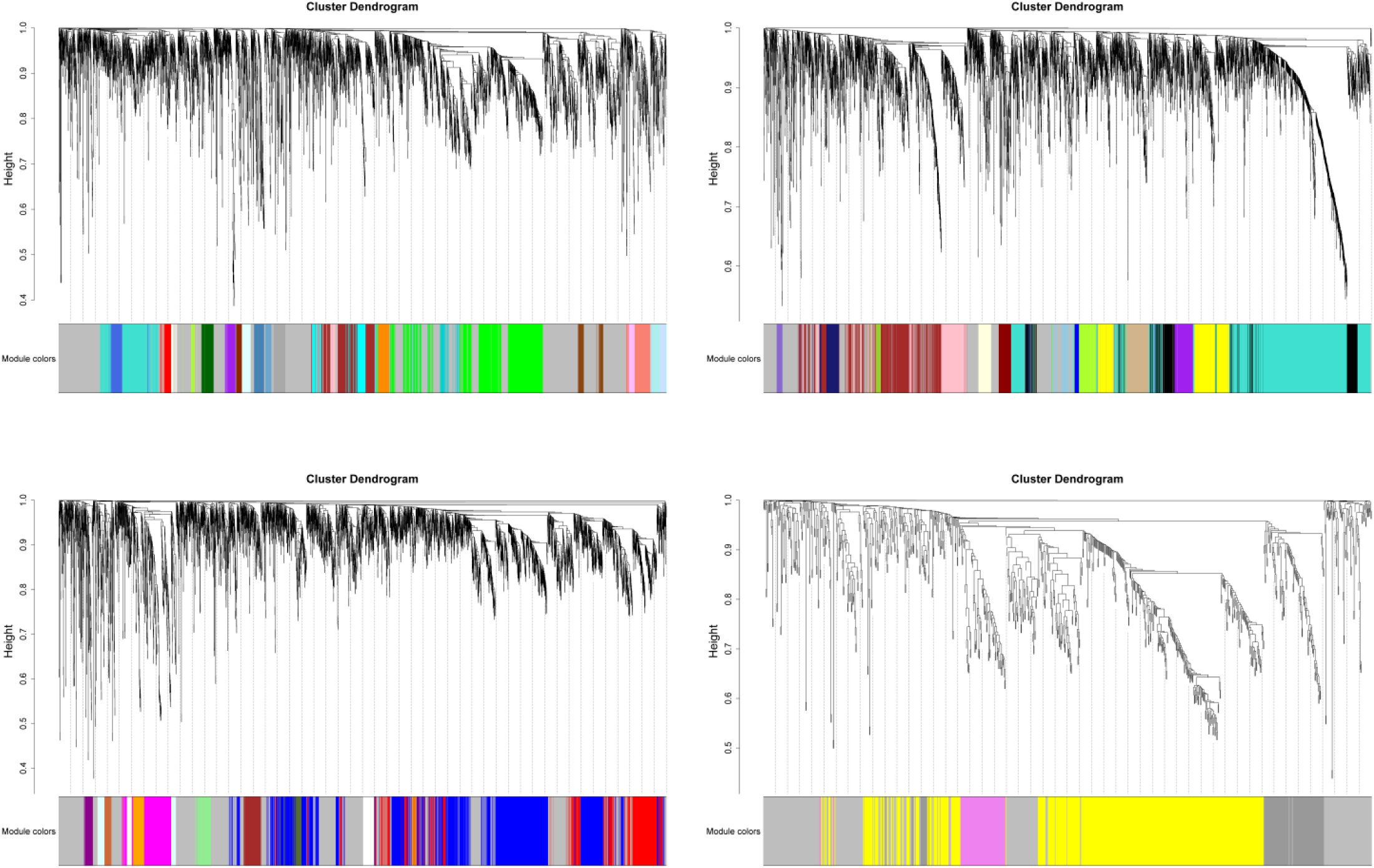
Feature dendrograms of weighted gene co-expression network analysis (WGCNA) identifying 46 modules of four block wise clusters. First cluster comprised 4998 features, the second 4975, the third 4863 and the fourth 972.

**Figure S28.**
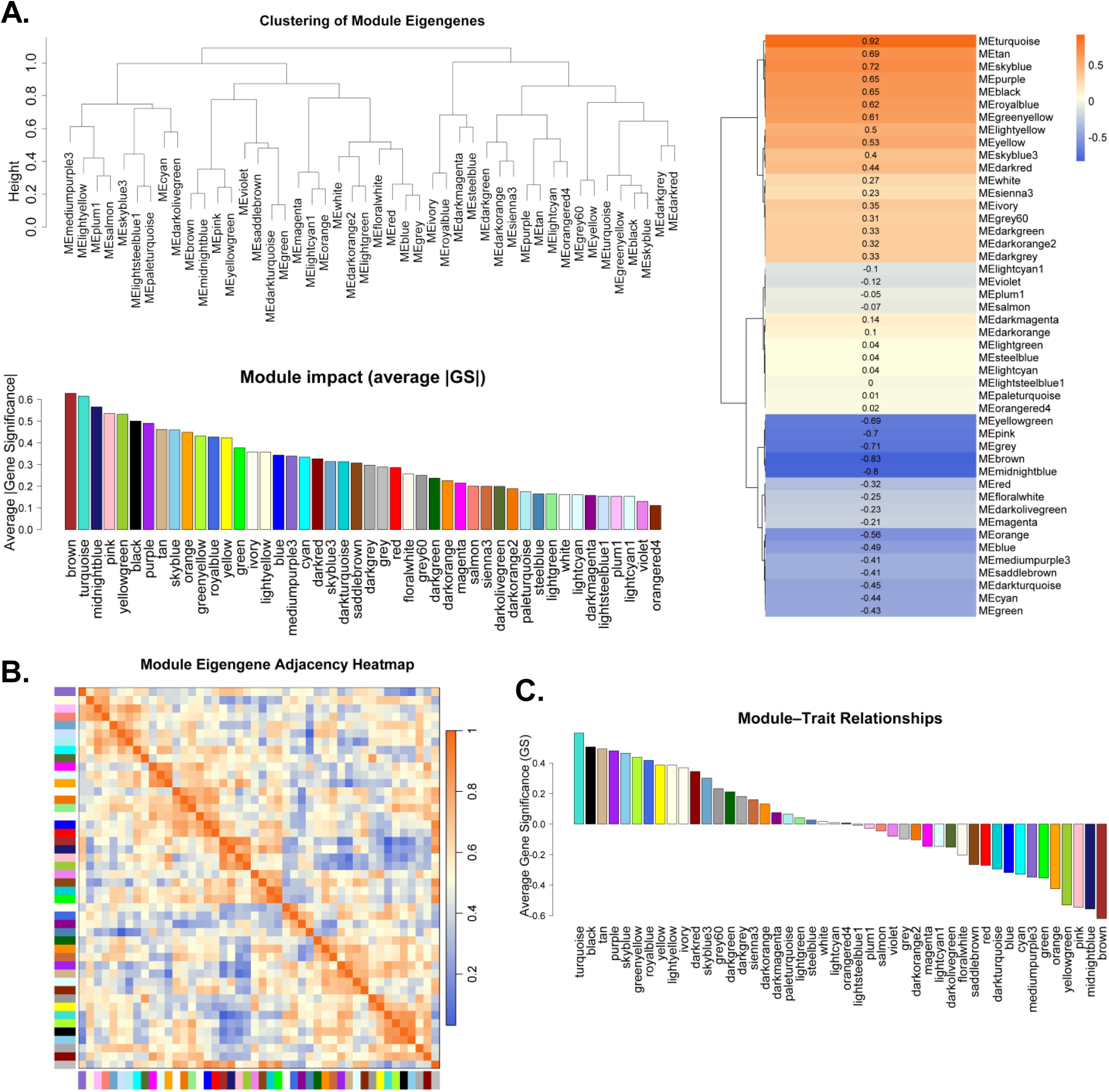
Module overview of weighted gene co-expression network analysis (WGCNA). WGCNA identified 46 modules across 15,809 proteomic, metabolomic and transcriptomic features. (**A**) Dendrogram of the clustering of module eigengenes with module impact as average |gene significance| (GS) as bar plot and as heatmap representation. (**B**) Heatmap of module eigengene adjacency. (**C**) Module-trait relationship of weighted gene co-expression network analysis (WGCNA) of 15,809 metabolic, transcriptomic and proteomic features across leaf, root and seed tissues of CHEN-384, LM2, and PI-665276.

**Figure S29.**
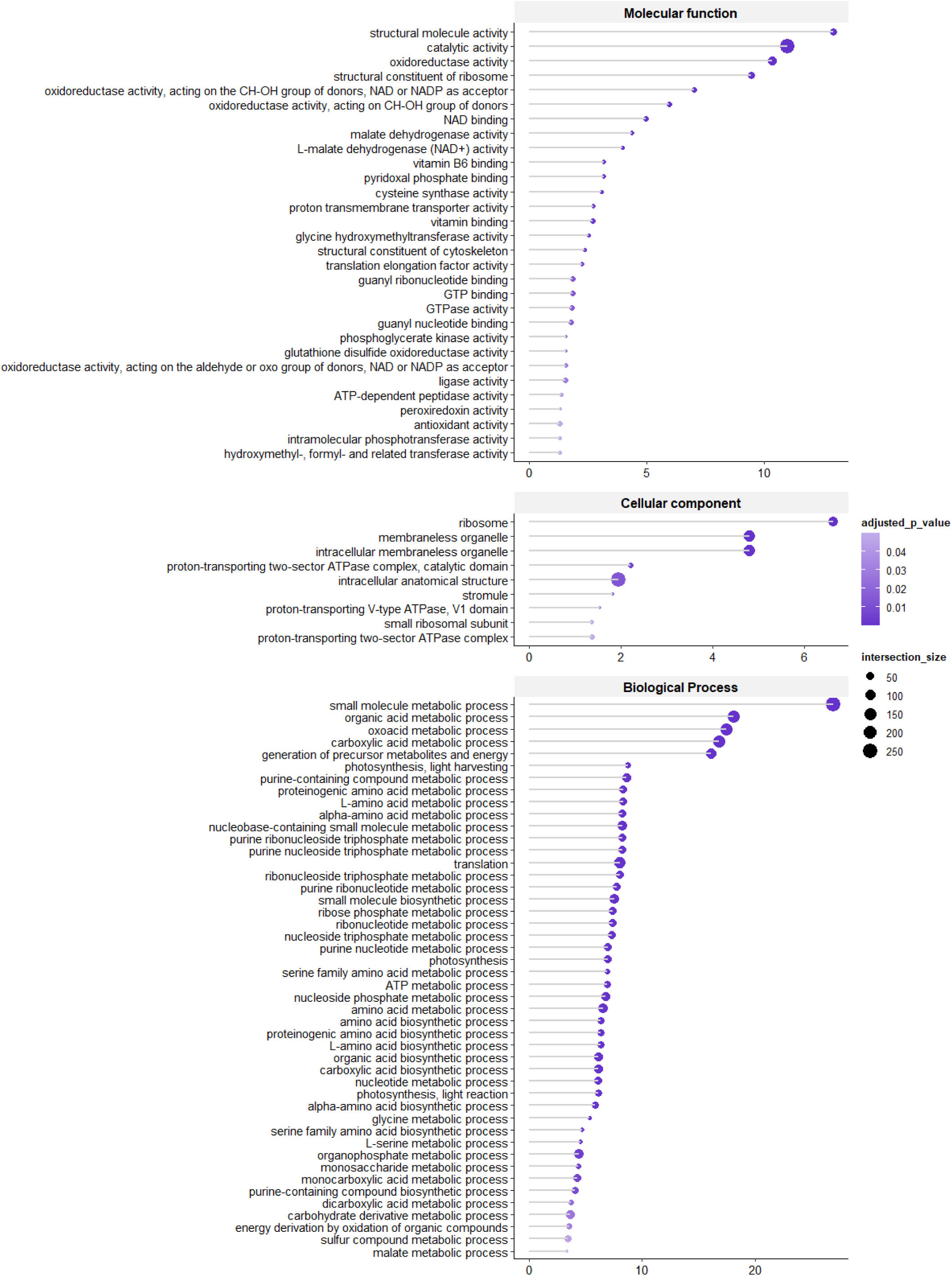
Gene ontology assignment of the leaf proteome of PI-665276, LM2, and CHEN-384 grown in the greenhouse under control and drought conditions. Significant proteins of leaves grown under drought and control conditions calculated by either Student’s *t*-test or by Wilcoxon test based on their normal distribution. Cut-off criterion: *p*-value < 0.05, |log_2_ fold change| > 1. For GO analysis g:Profiler was used with a cut-off criterion of adjusted *p*-value < 0.05 for molecular function and cellular component and < 0.001 for biological process.

**Figure S30.**
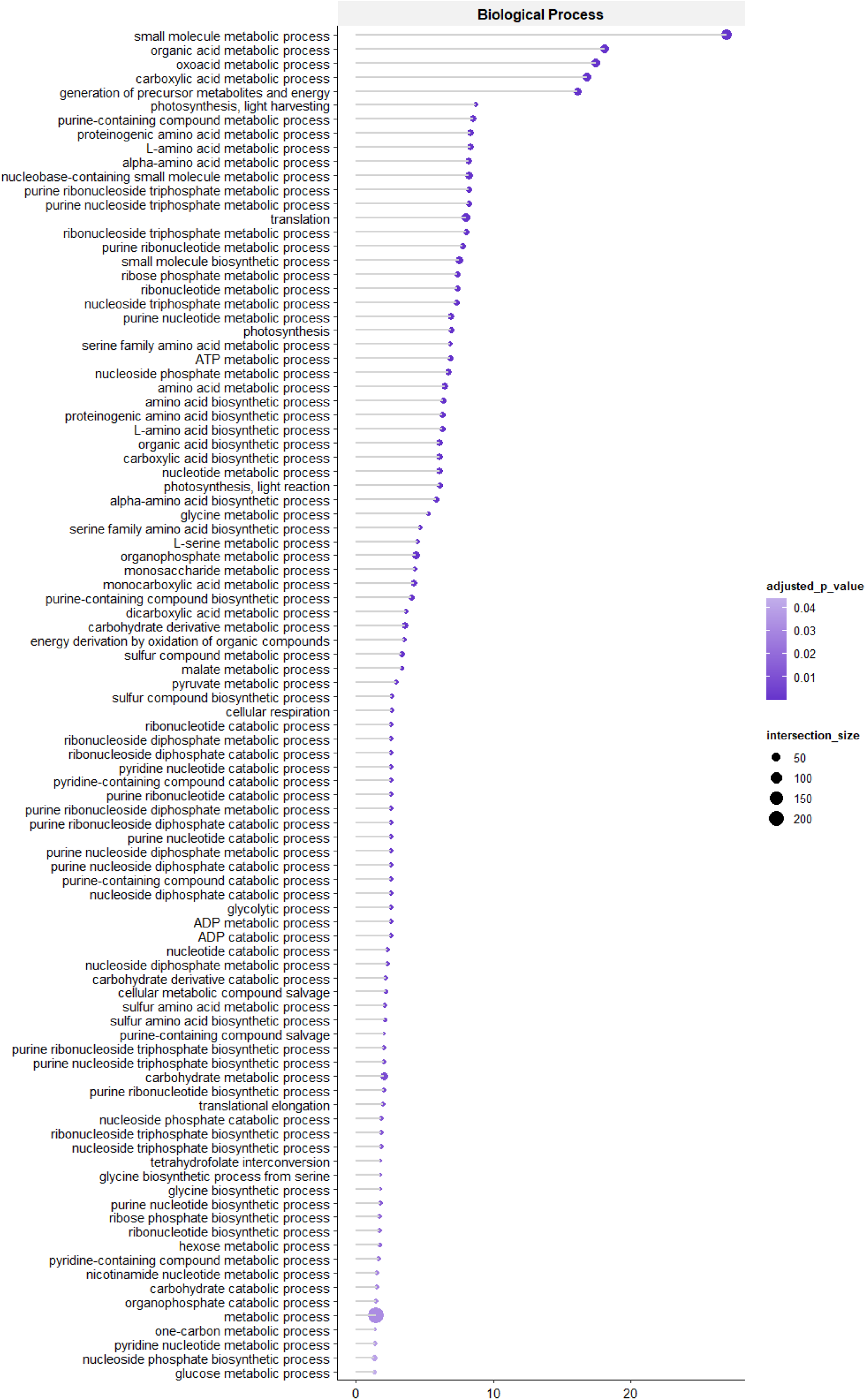
Significant leaf proteins of PI-665276, LM2, and CHEN-384 with an assignment of a biological process during gene ontology analysis. Significant proteins of leaves grown under drought and control conditions calculated by either Student’s *t*-test or by Wilcoxon test based on their normal distribution. Cut-off criterion: *p*-value < 0.05, |log_2_ fold change| > 1. For GO analysis g:Profiler was used with a cut-off criterion of adjusted *p*-value < 0.05.

**Figure S31.**
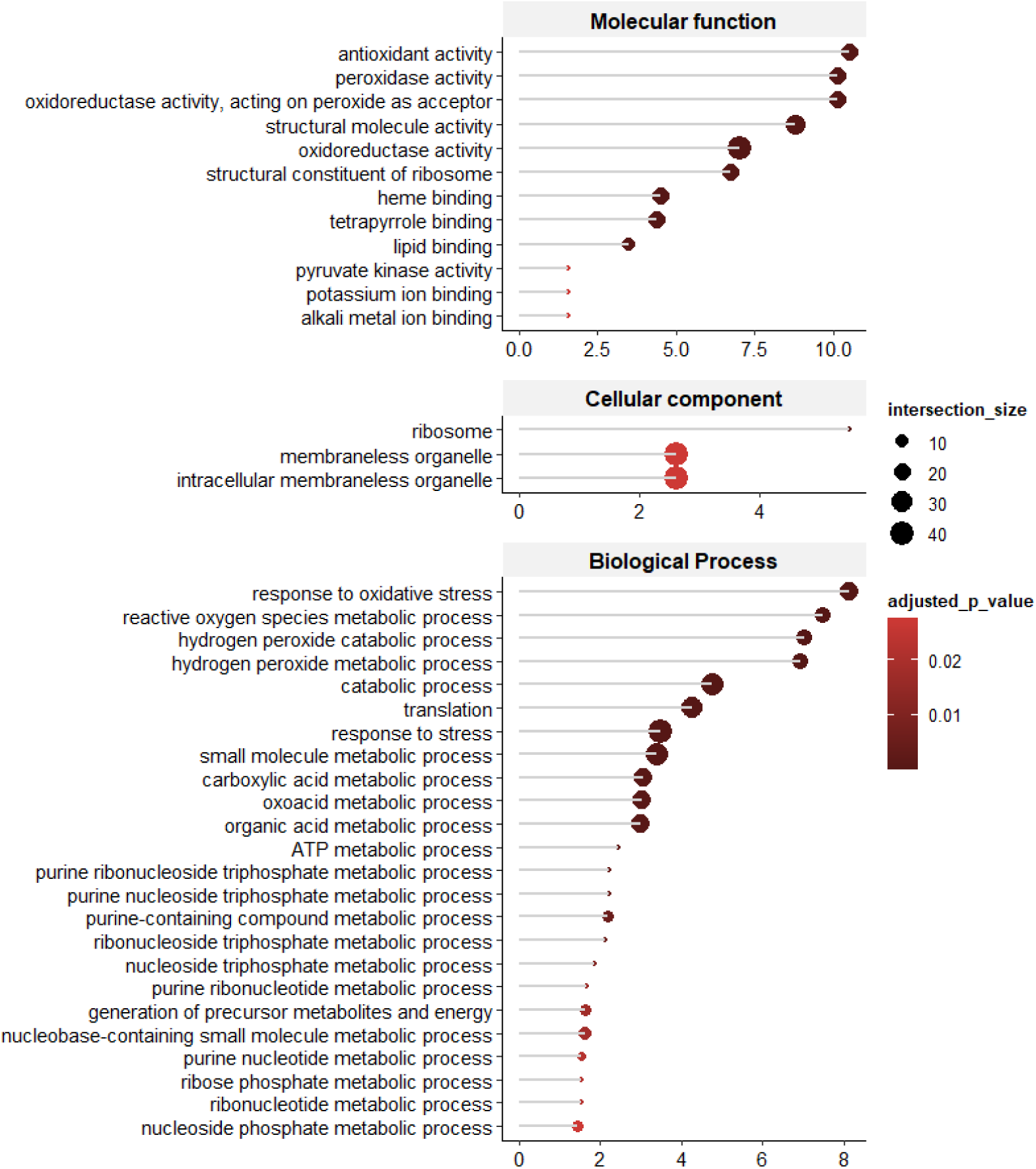
Gene ontology assignment of the root proteome of PI-665276, LM2, and CHEN-384 grown in the greenhouse under control and drought conditions. Significant proteins of leaves grown under drought and control conditions calculated by either Student’s *t*-test or by Wilcoxon test based on their normal distribution. Cut-off criterion: *p*-value < 0.05, |log_2_ fold change| > 1. For GO analysis g:Profiler was used with a cut-off criterion of adjusted *p*-value < 0.05.

**Figure S32.**
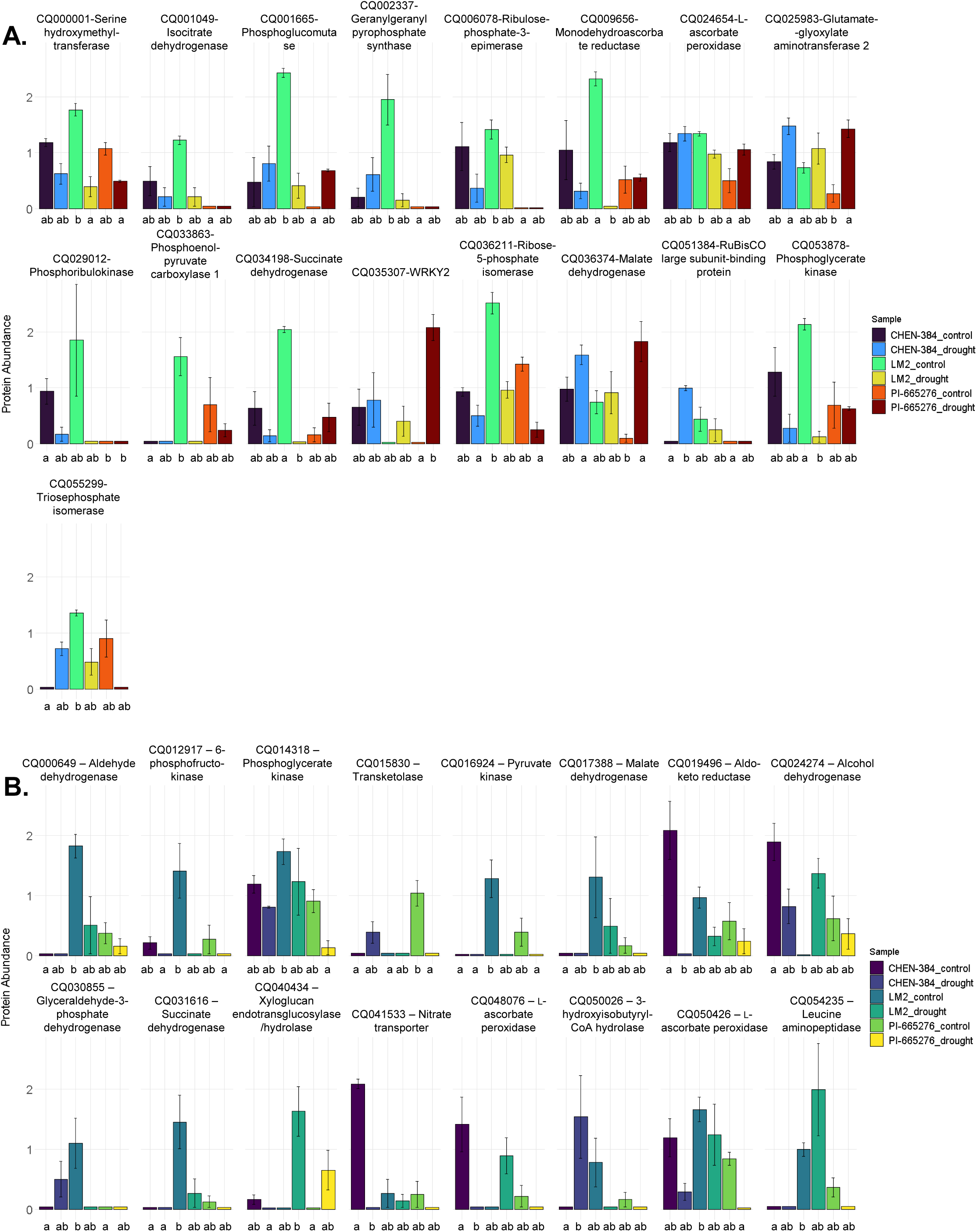
Selected differential expressed proteins of leaves and roots of PI-665276, LM2, and CHEN-384 grown in the greenhouse under control and drought conditions. Proteins involved in drought response in (**A**) leaves and (**B**) roots. Significances were determined using either two-way-ANOVA with post-hoc Tukey HSD test or Kruskal-Wallis test with post hoc Dunn’s test based on the data normal distribution.

**Figure S33.**
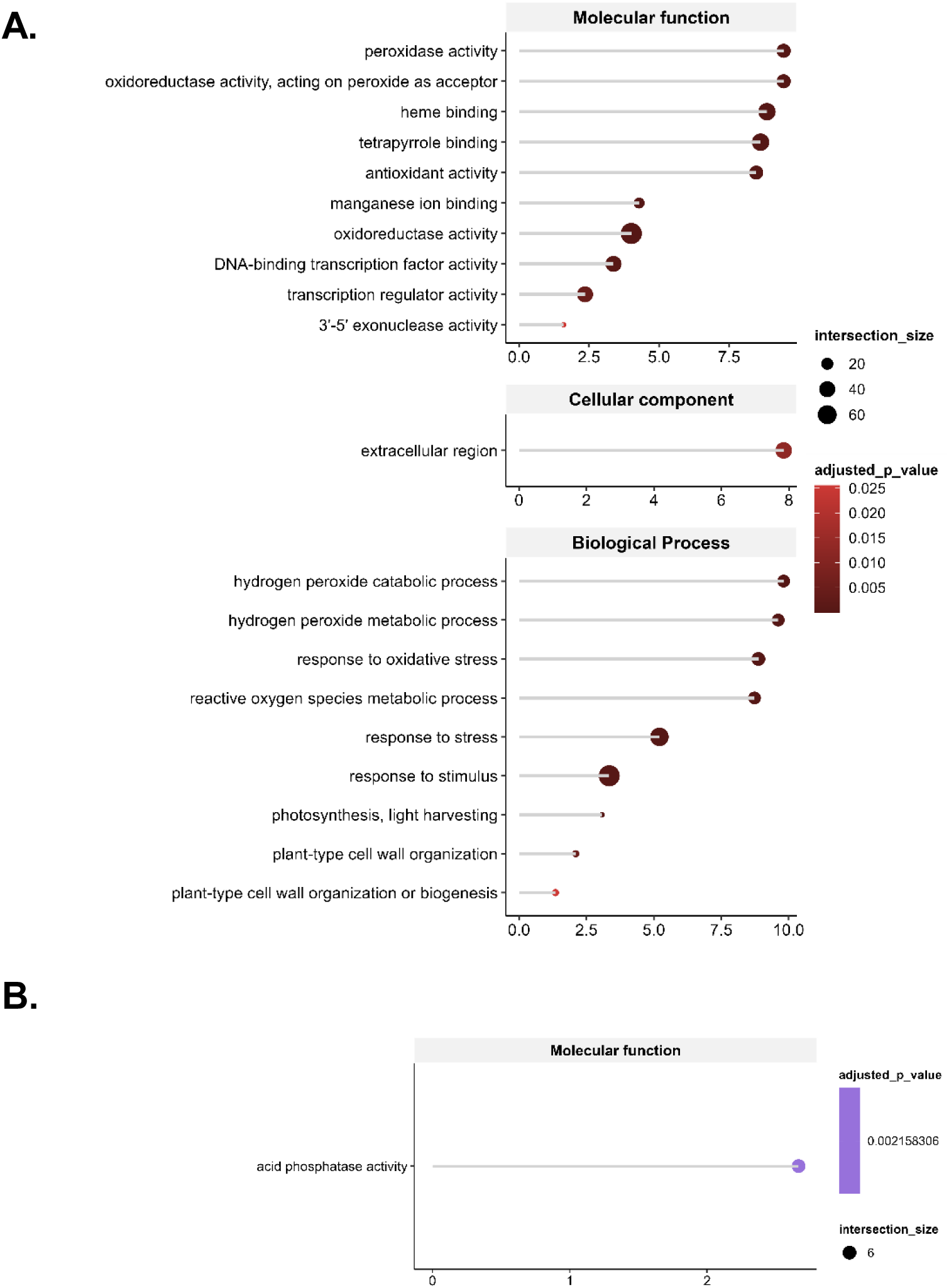
Gene ontology assignment of the root and leaf transcriptome of PI-665276, LM2, and CHEN-384 grown in the greenhouse under control and drought conditions. Significant transcripts of A) roots and B) leaves grown under drought and control conditions calculated by either Student’s *t*-test or by Wilcoxon test based on their normal distribution. Cut-off criterion: *p*-value < 0.05, |log_2_ fold change| > 1. For GO analysis g:Profiler was used with a cut-off criterion of adjusted *p*-value < 0.05.

